# A *Xanthomonas* effector increases virulence quantitatively by inducing at least three minor susceptibility genes

**DOI:** 10.1101/2025.07.31.667891

**Authors:** Brice Charleux, Carine Gris, Sébastien Carrère, Alvaro L. Pérez-Quintero, Aurélie Le Ru, Caroline Bellenot, Ivanna Fuentes, Zoë E. Dubrow, Adam J. Bogdanove, Laurent D. Noël, Corinne Audran

## Abstract

The transcription activator-like effector Tal12a is widely conserved among *Xanthomonas campestris* pv. *campestris* strains that cause black rot in Brassica crops. However, its role in disease remains unclear. To investigate how Tal12a contributes to pathogenesis, we combined transcriptomic profiling of cauliflower leaves infected with Tal12a-expressing strains, prediction of TALE-binding elements, and heterologous expression assays in *Nicotiana benthamiana.* The aim of this approach was to identify candidate susceptibility genes. Artificial TALEs were further employed to validate the contribution of these candidate targets to disease development. Tal12a enhanced both virulence and bacterial growth in cauliflower. Transcriptome analysis revealed 380 genes that were induced upon infection, nine of which were prioritised as candidates. Seven of these were confirmed as direct Tal12a targets, while the induction of the sugar transporter genes *BoSWEET13* and *BoSWEET14c* likely occurred indirectly. Functional assays demonstrated that these two *SWEET* genes and the *BoIAA7c* gene, which encodes auxin-dependent transcriptional regulator, contribute to disease development. These findings identify the first susceptibility genes in cauliflower and reveal that Tal12a promotes disease through a complex transcriptional reprogramming, involving direct and indirect target induction, with several genes functioning as minor susceptibility factors.

## INTRODUCTION

Phytopathogenic bacteria have evolved sophisticated molecular strategies to manipulate host physiology and establish disease. Among the virulence factors employed, Transcription Activator-Like Effector proteins (TALEs) represent a unique class of type III effectors translocated into the plant cell via the bacterial type-III secretion system (Boch & Bonas, 2010). TALEs are targeted to the nucleus thanks to their C-terminal nuclear localization signals (Yang & Gabriel, 1995; Szurek *et al*., 2002) where they act as eukaryotic transcription factors by binding to DNA, and increasing the expression of host genes (Hutin *et al*., 2015a). TALEs are deployed predominantly by *Xanthomonas* and *Ralstonia* species where they directly upregulate expression of plant genes, some of which promote colonization and infection spread and are referred to as *Susceptibility* (*S*) genes (Kay & Bonas, 2009; Boch & Bonas, 2010; de Lange *et al*., 2013). TALE proteins bind to specific plant DNA sequences through their central DNA binding domain. This domain is composed of a series of nearly identical repeats, each characterized by two critical amino acids known as Repeat-Variable Diresidues (RVDs), which define the specificity of DNA base recognition (Moscou & Bogdanove, 2009; Boch *et al*., 2009). The combination of RVDs within a TALE determines its binding to specific DNA sequences known as effector binding elements (EBEs). As a consequence, the analysis of RVD sequences using bioinformatics tools, such as TALvez, PrediTALE or TALE-NT enables the prediction of potential EBE for a given TALE within a genome (Doyle *et al*., 2012; Pérez-Quintero *et al*., 2013; Erkes *et al*., 2019). Building on this knowledge, researchers have developed artificial TALEs (arTALEs) which are engineered to recognize custom DNA sequences. These synthetic effectors have become widely used tools to specifically induce host gene expression and test the biological function of candidate genes during infection (Weber *et al*., 2011; Streubel *et al*., 2013).

Among the well-characterized *S* genes, the clade III members of the *Sugars Will Eventually be Exported Transporters* (*SWEET*) gene family have received considerable attention due to their targeting by various TALEs in numerous pathosystems and their key biological role during infection (Gupta *et al*., 2021; Baruah et *al*., 2025). For instance, TALE-mediated transcriptional upregulation of any of the rice genes *OsSWEET11, OsSWEET13* and *OsSWEET14* is essential for disease symptom development and enhanced bacterial proliferation (Yang *et al*., 2006; Streubel *et al*., 2013; Zhou *et al*., 2015). Failure to induce *SWEET* gene expression by TALEs yields rice plants essentially resistant to *Xanthomonas oryzae* pv. *oryzae* (*Xoo*) (Antony *et al*., 2010; Liu *et al*., 2011; Hutin *et al*., 2015b; Eom *et al*., 2019; Oliva *et al*., 2019; Schepler-Luu *et al*., 2023). Beyond rice, TALE-mediated activation of *SWEET* genes has also been reported in cotton and cassava, indicating that the exploitation of *SWEET* genes represents a convergent strategy employed by various *Xanthomonas* species to facilitate infection (Cohn *et al*., 2014; Cox *et al*., 2017). Other examples of *S* genes activated by TALEs include *OsSULTR3;6*, a putative sulfate transporter gene associated with susceptibility to bacterial leaf streak in rice (Cernadas *et al*., 2014) and TaNCED-5BS, a gene encoding a key enzyme involved in abscisic acid biosynthesis and plant defence inhibition during bacterial streak disease in wheat (Peng *et al*., 2019). In addition, several transcription factors have been identified as TALE targets, including *CsLOB1* in citrus (Hu et al., 2014) as well as *OsTFX1* and *OsERF123* in rice, where they contribute to disease susceptibility (Sugio *et al*., 2007; Tran *et al*., 2018).

Black rot, caused by *Xanthomonas campestris* pv. *campestris* (*Xcc*), is a major vascular and seed-transmitted disease that affects all cultivated brassica crops worldwide (Vicente & Holub, 2013; Dubrow & Bogdanove, 2021). After infection of hydathodes at the leaf margins, *Xcc* spreads through the xylem vessels and colonizes the mesophyll (Vicente & Holub, 2013; An *et al*., 2020). Unlike other *Xanthomonas* species, the presence of *tal* genes in pathogenic *Xcc* strains is highly variable, with some strains harboring several TALEs, while others are completely devoid of them. A recent study demonstrated that inhibition of *tal* genes by CRISPR interference (CRISPRi) in the *Xcc* strain CN08 resulted in decreased disease symptoms in cauliflower (Zárate Chaves *et al*., 2023). Interestingly, a reduction in *Xcc* pathogenicity was also reported in the cauliflower pathosystem with the double deletion mutant Δ*tal12a/15a* in strain Xca5 (Denancé *et al*., 2018). Among the TALEs identified, Tal12a, which comprises 12 RVDs, is the most widespread TALE effector in an *Xcc* diversity panel of 23 strains (Denancé *et al*., 2018).

The present study aimed to determine the contribution of Tal12a to *Xcc* virulence in cauliflower and to identify its host targets. We showed that Tal12a enhances bacterial pathogenicity and *in planta* proliferation. We identified 380 candidate target genes of Tal12a in cauliflower by a transcriptomic approach. Among these, seven genes were confirmed as direct targets. We demonstrated that *BoSWEET13*, *BoSWEET14c* and *BoIAA7c* play a role in host susceptibility. These findings suggest that Tal12a manipulates both auxin signaling and sugar transport pathways to promote disease susceptibility in cauliflower.

## RESULTS

### *tal12a* contributes to the virulence of *Xcc* in cauliflower

To investigate the biological importance of *tal12a* in the CN08 strain, we evaluated the pathogenicity of a spontaneous mutant lacking this gene (CN08Δ*tal12a*) and a complemented strain carrying *tal12a* on the pSKX1 vector (CN08Δ*tal12a* pSKX1-*tal12a*). Western blot analysis confirmed the absence/presence of the 12-repeat-long TALE corresponding to Tal12a in strains CN08Δ*tal12a* and CN08Δ*tal12a* pSKX1-*tal12a*, respectively while the expression of the other three TALEs (15e, 18a and 22a repeat-long TALEs) remained unchanged (Fig. **S1a**). *Xcc* strains were wound-inoculated in the leaf midvein of susceptible cauliflower (cv. Clovis) and symptoms were monitored at four and seven days post-inoculation (dpi) (Fig. **1a,b**). The CN08Δ*tal12a* mutant caused significantly less chlorosis and necrosis on cauliflower than the wild type. Symptom development was fully restored upon complementation with *tal12a*, demonstrating Tal12a contribution to the virulence of strain CN08.

**Fig. 1.**
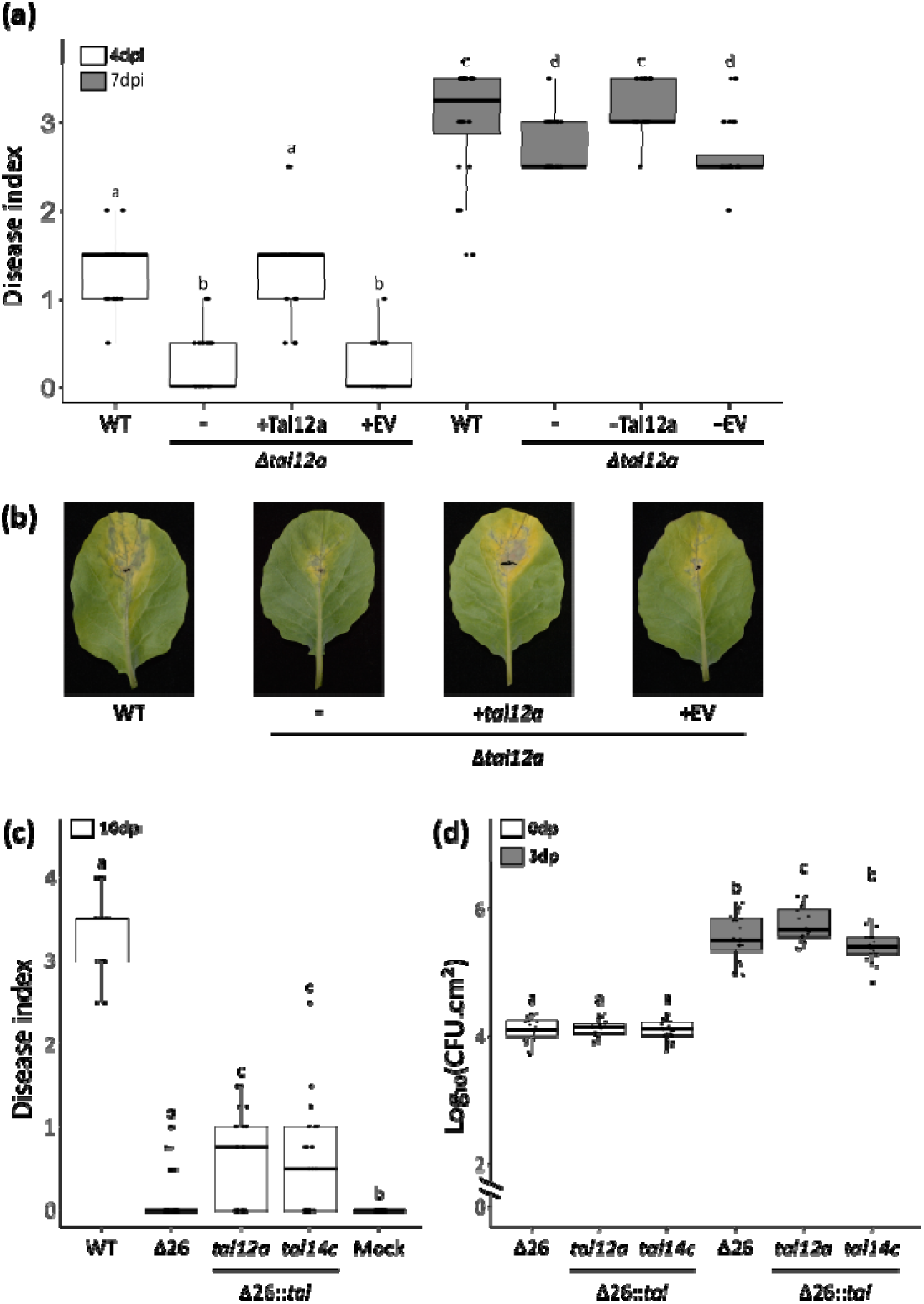
*tal12a* promotes bacterial virulence and proliferation on cauliflower. (**a**) Boxplot representation of disease symptoms caused by *Xcc* CN08 strain at 4 and 7 days post inoculation (dpi), respectively. Pathogenicity assays were performed using the WT *Xcc* strain CN08, its *tal12a* deletion mutant CN08Δ*tal12a* and the mutant complemented with either the empty vector pSKX1 (+EV) or the *tal12a* gene (+*tal12a*). (**b**) Representative photographs for the median disease symptoms caused by *Xcc* CN08 strain derivatives at 7 dpi. (**c**) Boxplot representation of disease symptoms caused by *Xcc* at 10 dpi. Pathogenicity was evaluated in the WT *Xcc* strain 8004 or its Δ26 polymutant derivative, which lacks 26 genes encoding type III effectors, either alone or complemented with *tal12a* or *tal14c* integrated into the genome. (**d**) *Xcc* Δ26 strain titers in cauliflower leaves were quantified at 0 and 3 days after mesophyll infiltration with a bacterial suspension at 10^6^ CFU.mL^−1^. Results represent four independent experiments with six plants per strain. Statistical groups were determined using the nonparametric Kruskal–Wallis test (p < 0.05) and are indicated by different letters. (**a-c**) Cauliflower leaves (cv. Clovis) were wound inoculated by piercing the central vein with a needle dipped in a bacterial suspension at 10^8^ CFU.mL^−1^. The disease index was as follows: 0, no symptoms; 1, weak chlorosis around wounded sites; 2, strong chlorosis; 3, first necrotic symptoms; 4, large necrotic lesions as described in Luneau *et al*. (2022). Results correspond to three independent experiments with at least six plants per strain. The statistical groups were determined using the nonparametric Kruskal–Wallis test (p < 0.05) and are indicated by different letters.

To specifically assess the contribution of *tal12a* in *Xcc* virulence on cauliflower by a gain-of-function experiment, we used a disarmed derivative of *Xcc* strain 8004 named 8004Δ26T3E (hereafter referred to as Δ26; European patent EP20305940.7). This polymutant strain possesses a functional type III secretion system, naturally lacks endogenous *tal* genes, and has been engineered through the successive deletion of twenty-six genes encoding type-III effector proteins. The Δ26::*tal12a* and Δ26::*tal14c* strains harbor a stable genomic integration of *tal12a* or *tal14c*, a 14-repeat *tal* gene used as a control to assess the specificity and function of *tal12a* targets (Denancé *et al*., 2018). Expression of the Tal12a and Tal14c proteins was confirmed by Western blot analysis in the Δ26::*tal12a* and Δ26::*tal14c* strains, respectively (Fig. **S1b**). These two strains showed no detectable fitness deficit *in vitro* relative to the parental Δ26 strain, as measured by maximum growth rates (µmax) in MME and MOKA culture media (Fig. **S2**). At 10 dpi, the Δ26 strain failed to induce disease symptoms, similar to the mock-inoculated plants, while the wild-type 8004 strain caused severe chlorotic and necrotic symptoms (Fig. **1c**). Interestingly, Δ26 strains expressing either tal12a or tal14c exhibited increased disease severity compared to the Δ26 strain (p < 0.01), suggesting that each TALE can partially restore virulence. To further evaluate their contribution to bacterial proliferation, bacterial titers were measured at 3 dpi (Fig. **1d**). A significant increase in bacterial growth was observed for the Δ26::*tal12a* strain but not Δ26::*tal14c* compared to Δ26 (Fig. **1d**). Together, these results establish *tal12a* as a virulence factor of *Xcc*, enhancing both symptom development and bacterial colonization in cauliflower.

### Tal12a increases directly or indirectly the expression of 380 cauliflower genes

To identify the cauliflower genes whose expression is specifically activated by Tal12a, we conducted a transcriptomic analysis using RNA sequencing on leaves infected with the Δ26::*tal12a* strain, comparing them to leaves infected with the Δ26 strain at 1 dpi. The Δ26::*tal14c* strain was included to evaluate the specificity of Tal12a targets. The properties of the RNAseq libraries are given in Table **S1**. As expected from transcriptional activators, only a small number of genes were found to be downregulated following infection with the Δ26::*tal12a* strain (10 genes) and the Δ26::*tal14c* strain (38 genes). This suggests that these are likely to be the result of indirect TALE-dependent regulation (Fig. **2a**). The expression of 380 genes was increased (fold change ≥ 2; FDR < 0.01) by Δ26::*tal12a* (Fig. **2a**, Table **S8**). For comparison, the expression of 713 genes was upregulated (fold change ≥ 2; FDR < 0.01) by Δ26::*tal14c* (Table **S8**). Among these, 43 genes were induced by both TALEs. A hypergeometric test confirmed that this overlap was statistically significant (p-value < 0.001) and unlikely to occur by chance. Gene Ontology (GO) enrichment analysis of these 43 genes revealed a strong and significant enrichment in categories related to “jasmonic acid” (JA), “wound responses” and “regulation of defense responses” (Fig. **2b**). These results suggest that TALEs may modulate disease-associated stress responses by converging on shared downstream pathways. This could be due to the activation of shared target genes or functionally redundant paralogues, or to plant perception mechanisms that respond similarly to TALEs, regardless of their DNA-binding specificities and target gene identities.

**Fig. 2.**
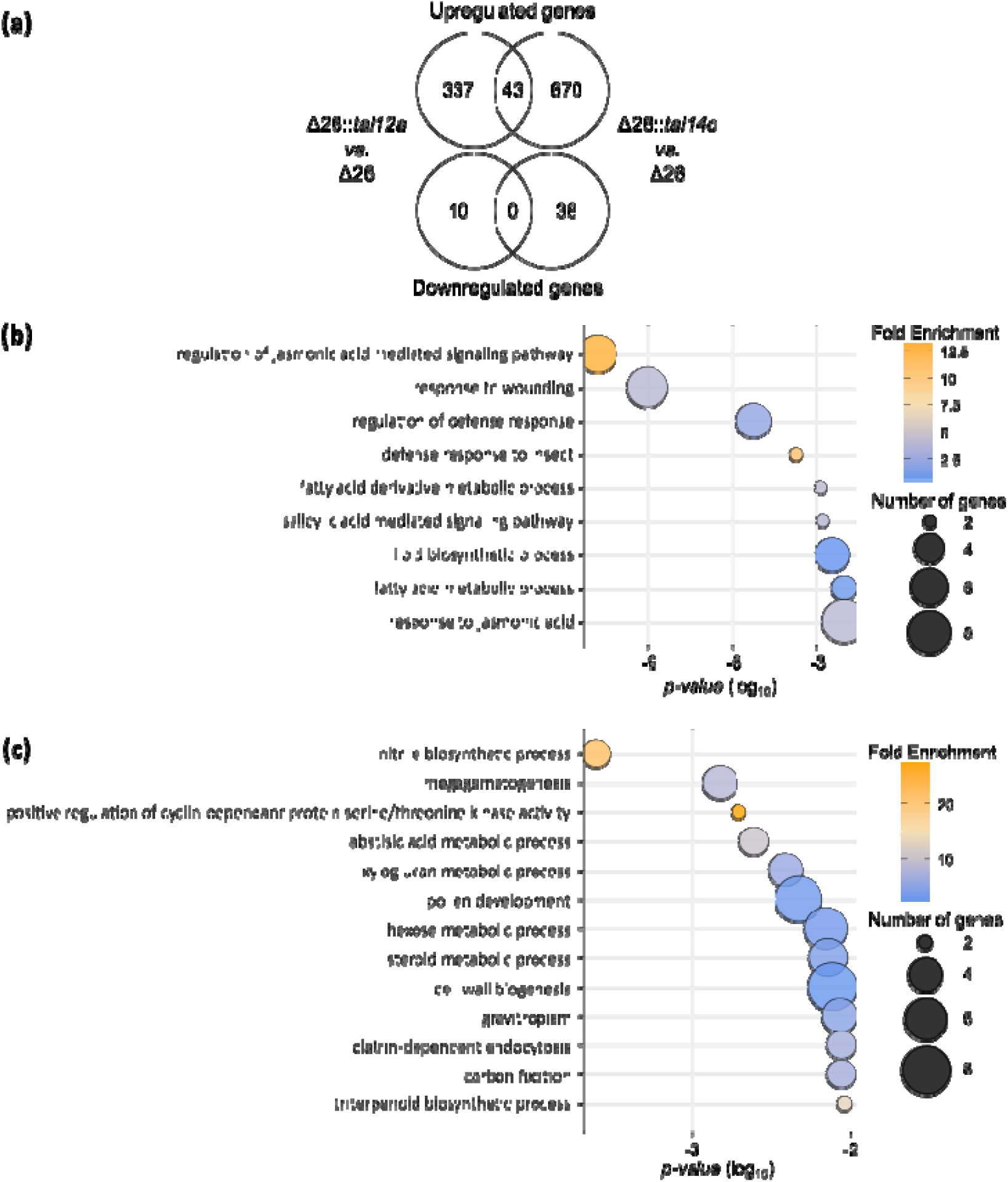
Tal12a induces expression of 380 cauliflower genes. Transcriptomic analysis was performed on cauliflower leaves infected by Δ26 strain derivatives expressing either Tal12a (Δ26::*tal12a*) or Tal14c (Δ26::*tal14c*) at 1 dpi and compared to strain Δ26. (**a**) Venn diagram showing genes whose expression is regulated by Tal12a and Tal14c (FDR < 0.01, Log_2_ fold change > 2). (**b,c**) Gene ontology (GO) analysis highlights the biological processes enriched among genes regulated by both *tal12a* and *tal14c* (**b**) and by *tal12a* only (**c**). The *x*-axis represents the enrichment probability of a given GO term. The size of the circle represents gene count. The color scale represents the enrichment of the GO term in the up-regulated gene list compared to the background population of genes.

GO-term enrichment analysis for genes specifically upregulated by Tal12a revealed distinct biological processes compared to those associated with Tal14c (Fig. **2c**, Table **S2**). GO terms such as "megagametogenesis", “ABA metabolic pathway”, “pollen development”, and cell-wall-related categories (“xyloglucan metabolic process”, “pollen development”, “hexose metabolic process”, “cell wall biogenesis”) were significantly enriched among Tal12a-regulated genes, indicating that Tal12a impacts physiological pathways distinct from Tal14c. Notably, the term “pollen development” was associated with the expression of two clade III *SWEET* genes, *BoSWEET13* and *BoSWEET14c* (Zhang et al., 2019). Interestingly, *BoSWEET13* is the most upregulated (greatest fold-change) gene in the transcriptomic analysis when comparing Δ26::*tal12a* to Δ26 (Table **1**). These genes are homologous to *OsSWEET13* and *OsSWEET14* in rice, which are known *S* genes induced by TALEs such as PthXo2, PthXo3, AvrXa7, TalC and TalF from *Xoo* (Teper et al., 2023). Therefore, *SWEET* genes are considered to be promising candidates for the *S* gene that could mediate the virulence function of Tal12a in cauliflower.

**Table 1.** Cauliflower genes induced by Tal12a at 1 dpi and selected for downstream analysis.

| Gene rank<sup>a</sup> | Gene ID<sup>b</sup> | Log<sub>2</sub>FC | FDR | Product | Name | Predictions |  |  |  |
| --- | --- | --- | --- | --- | --- | --- | --- | --- | --- |
|  |  |  |  |  |  | TALVEZ |  | PrediTAL |  |
|  |  |  |  |  |  | score<sup>c</sup> | rank<sup>d</sup> | score<sup>c</sup> | rank<sup>d</sup> |
| 1 | Bo9g097170 | 8.95 | 3,0x10<sup>-127</sup> | Putative sugar transporter SWEET | *BoSWEET13* |  |  |  |  |
| 2 | Bo3g023170 | 8.89 | 1.5x10<sup>-99</sup> | Putative cytochrome P450 | *BoCYP450* | 10.8 | 33 | 0.55 | 125 |
| 12 | Bo3g171090 | 6.55 | 1.5x10<sup>-27</sup> | Putative transcription factor AP2- EREBP family | *BoERF043* |  |  | 0.54 | 176 |
| 15 | Bo6g112570 | 6.37 | 3.9x10<sup>-113</sup> | Putative transcription factor Homobox- WOX family | *BoATHB-X* | 9.9 | 158 | 0.54 | 163 |
| 48 | Bo1g103470 | 4.80 | 8.8x10<sup>-44</sup> | Putative transcription factor interactor and regulator AUX- IAA family | *BoIAAa* | 9.7 | 207 |  |  |
| 50 | Bo5g093110 | 4.69 | 8.1x10<sup>-68</sup> | Putative transcription factor interactor and regulator AUX- IAA family | *BoIAAc* | 9.7 | 216 |  |  |
| 77 | Bo2g013210 | 4.07 | 2.2x10<sup>-64</sup> | Putative Heat shock chaperonin-binding, STI1/HOP, DP domain-containing protein | *BoTIC40* | 10.2 | 96 | 0.55 | 153 |
| 98 | Bo3g081600 | 3.78 | 4.6x10<sup>-18</sup> | Putative transcription factor interactor and regulator AUX- IAA family | *BoIAAb* | 9.7 | 212 |  |  |
| 114 | Bo7g109890 | 3.52 | 8.2x10<sup>-07</sup> | Putative sugar transporter SWEET | *BoSWEET14c* |  |  |  |  |

Gene rank, Log<sub>2</sub>FC and FDR values are from RNAseq data, leaf infection and harvest at 1 dpi. Predictions are from TALVEZ and PrediTAL softwares.
<sup>a</sup> Rank sorted by gene induction level from RNAseq data.
<sup>b</sup> Gene ID corresponds to *Brassica oleracea* public annotation.
<sup>c</sup> Score indicated by prediction software. Presence of a score means prediction of putative target, the higher the better. No putative targets are indicated by absence of score.
<sup>d</sup> Rank sorted by prediction software, based on the putative targets found.

### A set of 29 genes identified as candidate direct targets of Tal12a in cauliflower

In order to identify direct targets of Tal12a in cauliflower, promoters of genes significantly upregulated (FDR < 0.01) by Tal12a were inspected for predicted effector binding elements (EBE) using TALVEZ and PrediTALE (Pérez-Quintero *et al*., 2013; Erkes *et al*., 2019). We restricted our analysis to the 300 predictions with the highest EBE scores. Among the 380 genes upregulated by Tal12a, 29 genes harbored high-score EBE predictions within their promoter regions, as identified by at least one of the two prediction tools (Fig. **3a**). Notably, none of the 29 candidate direct targets of Tal12a overlapped with those identified for Tal14c (Table **S3**). This list was refined using the features often found in direct TALE targets: (i) high EBE prediction scores, (ii) close proximity of EBEs to the TATA box, (iii) strong transcriptional induction. Considering gene annotations, seven candidate genes were prioritized for further characterization (Table **1** and Table **S3**). These genes encode a cytochrome P450 gene (*Bo3g023170*), a translocon protein TIC40 located on the inner chloroplast membrane (*Bo2g013210*) and five genes encoding transcription factors or related proteins (*ERF043* (*Bo3g171090*), *ATHB-X* (*Bo6g112570*), and three IAA7 isoforms (*Bo1g103470*, *Bo3g081600*, and *Bo5g093110*). In addition, given the well-established role of *SWEET* genes as major *S* genes in several pathosystems (Streubel *et al*., 2013; Cohn *et al*., 2014; Teper *et al*., 2023), we also studied *BoSWEET13* (*Bo9g097170*) and *BoSWEET14c* (*Bo7g109890*), which were identified in our RNA-seq analysis. These two genes are specifically expressed in the presence of Tal12a, despite the absence of an EBE in their respective promoters. This suggests that Tal12a may induce their expression through an indirect regulatory mechanism.

**Fig. 3.**
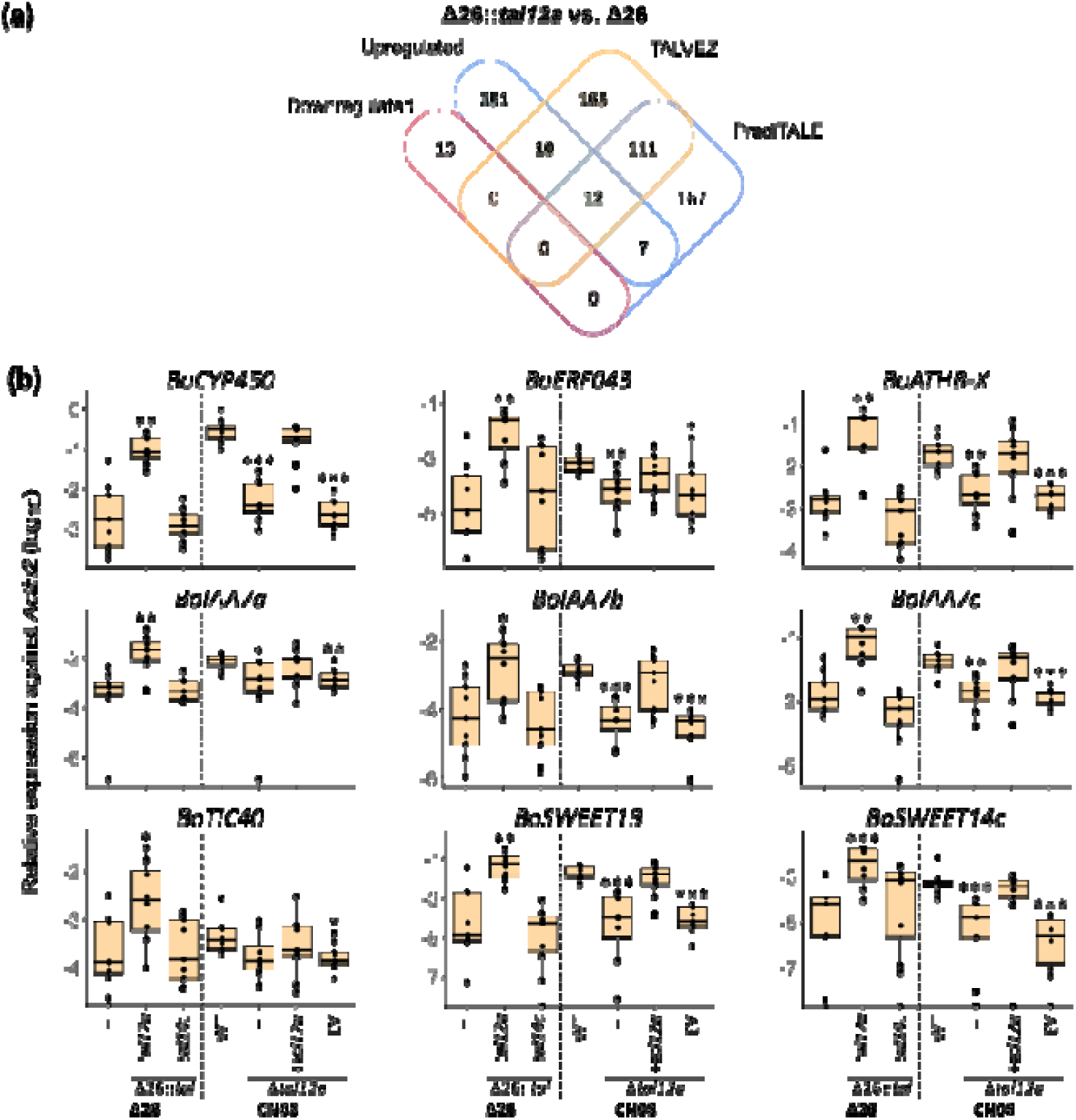
Tal12a specifically modulates the expression of cauliflower genes. (**a**) Venn diagram showing the intersections between genes differentially expressed in response to Tal12a (FDR<0.01, |log_2_FC| >2) and genes predicted to contain a Tal12a EBE using the PrediTALE and TALVEZ tools. The top 300 predicted targets were retained for comparison (Pérez-Quintero *et al*., 2013; Erkes *et al*., 2019). (**b**) Gene expression of nine putative Tal12a target genes following inoculation: *BoCYP450*, *BoERF043*, *BoIAA7a*, *BoIAA7b*, *BoIAA7c*, *BoTIC40*, *BoSWEET13*, *BoSWEET14c* and *BoATHB-X*. Cauliflower leaves were infected with various *Xcc* strains derivatives and sampled one day post-inoculation. The CN08Δ*tal12a* mutant was transformed with either the empty vector pSKX1 (EV) or with *tal12a* (+*tal12a*). The Δ26 polymutant strain and its derivatives expressing *tal12a* or *tal14c* (Δ26::*tal12a* or Δ26::*tal14c*, respectively) were inoculated. Each strain was tested in three independent experiments, each comprising three plants. Gene expression was measured by qRT-PCR. Data points represent at least eight measurements per condition. Statistical analysis was performed using a Wilcoxon test, comparing CN08 WT with its derivatives and Δ26 with its derivatives (*: p<0.05; **: p<0.01; ***: p<0.001).

To independently validate Tal12a-mediated induction of these nine genes, we quantified their expression in cauliflower by qRT-PCR following inoculation with *Xcc* strains carrying or lacking *tal12a*. Consistent with the RNA-seq results, gene expression analysis at 1 dpi revealed that introducing *tal12a* in the Δ26 polymutant strain significantly upregulated all nine candidate genes (Fig. **3b**). No upregulation of these genes has been observed for the condition Δ26 containing *tal14c*. In parallel, to evaluate the broader contribution of TAL effectors to the induction of these genes, we tested *Xcc* strains CN08 and Xca5 silenced for *tal* gene expression using a CRISPRi strategy targeting the conserved ribosome binding site (RBS) of *tal* genes as described in Zárate Chaves *et al*. (2023). Compared to both wild-type (WT) CN08 and control (Ctl) strains, eight out of nine Tal12a target genes showed a statistically significant decrease in expression in the CN08 CRISPRi-RBS strain relative to the wildtype and control strains (Fig. **4**). Similarly, expression of five out of nine target genes was significantly reduced in the Xca5 CRISPRi-RBS strain relative to WT Xca5 and control strains (Fig. **4**). These results demonstrate that the nine candidate genes are regulated in a TALE-dependent manner, although some degree of strain-to-strain variation is observed. The subtle differences observed between strains may result from incomplete CRISPRi-mediated silencing or variations in TALE accumulation levels. Since CN08 and Xca5 express four and three tal genes, respectively, and share *tal12a* and *tal22a* (Denancé et al., 2018), the CN08Δ*tal12a* mutant was used to determine the specific contribution of Tal12a to the regulation of the nine candidate genes. This deletion resulted in a statistically significant reduction in expression of the *BoCYP450*, *BoERF043*, *BoIAA7a*, *BoIAA7b*, *BoIAA7c*, *BoSWEET13*, *BoSWEET14c*, and *BoATHB-X* genes, while the expression of *BoTIC40* tended to decrease, but this reduction was not statistically significant (Fig. **3b**). Expression levels were restored to wild-type levels upon complementation with a plasmid carrying *tal12a* (Fig. **3b**). The various results obtained using different *Xcc* strains confirm that Tal12a promotes the expression of the selected genes.

**Fig. 4.**
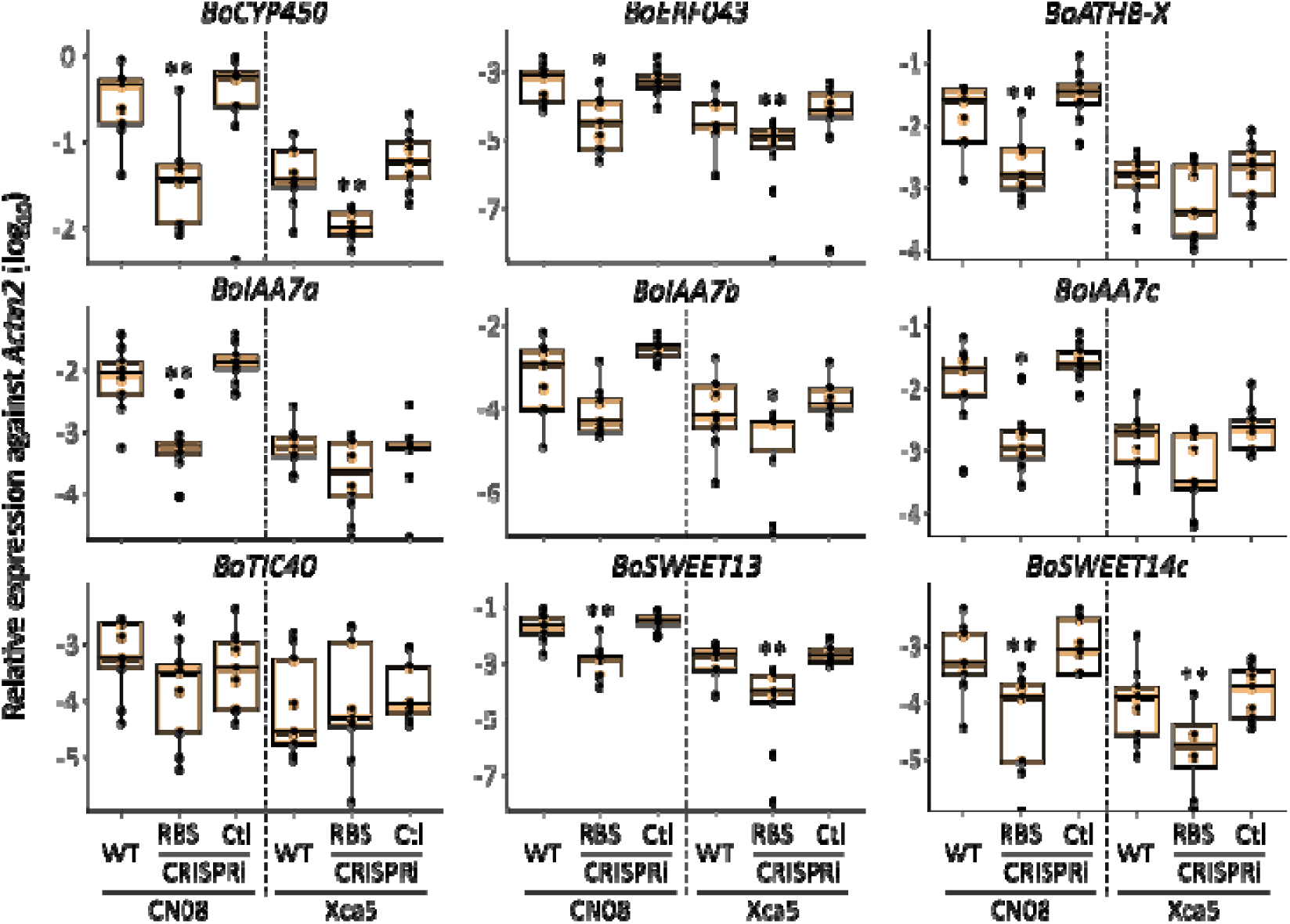
Silencing of *tal12a* in *Xcc* modulates the expression of potential target genes in cauliflower. Gene expression of nine potential Tal12a target genes (*BoCYP450*, *BoERF043*, *BoIAA7a*, *BoIAA7b*, *BoIAA7c*, *BoTIC40*, *BoSWEET13*, *BoSWEET14c*, and *BoATHB-X*).Cauliflower leaves were infected with various *Xcc* strains expressing or not Tal12a and sampled at 1 dpi. CN08 and Xca5 CRISPRi (RBS) and control (Ctl) strains are derivatives of the wild-type (WT) CN08 and WT Xca5 (Zárate Chaves *et al*., 2023). Each strain was tested in three independent experiments, each involving three plants. Data points represent at least eight measurements per condition. Statistical analysis was performed using a Wilcoxon test, with significance set at p<0.05, comparing WT with CRISPRi derivatives

### Transactivation assays in *N. benthamiana* revealed both direct and indirect regulation of Tal12a targets

To assess the direct regulation of the nine genes by Tal12a, promoter fragments (307–920 bp), including the predicted EBEs when present (Table **S4**), were fused upstream of a promoterless GUS reporter gene. These fusions were transiently delivered in *N. benthamiana* by *Agrobacterium tumefaciens* while Tal12a was delivered using the *Xcc* Δ26::*tal12a* strain.

The transactivation of the *GUS* gene expression was assessed by quantifying GUS activity in leaves at 2 dpi using ImageJ (Fig. **5**). All the promoter constructs tested showed negligible or low basal GUS activity in *N. benthamiana* leaves in the absence of *Xcc* strain. Significant increases in GUS activity were observed for the *BoCYP450*, *BoERF043*, *BoIAA7a*, *BoIAA7b*, *BoIAA7c*, *BoTIC40* and *BoATHB-X* promoter constructs upon co-delivery with Δ26::*tal12a*, relative to the Δ26 strain alone. On the contrary, no differences in GUS activity were observed for the *BoSWEET13* and *BoSWEET14c* constructs. These results support the EBE prediction and suggest that Tal12a directly regulates *BoCYP450*, *BoERF043*, *BoIAA7a*, *BoIAA7b*, *BoIAA7c*, *BoTIC40*, and *BoATHB-X*. However, Tal12a-mediated upregulation of *BoSWEET13* and *BoSWEET14c* appears to be indirect.

**Fig. 5.**
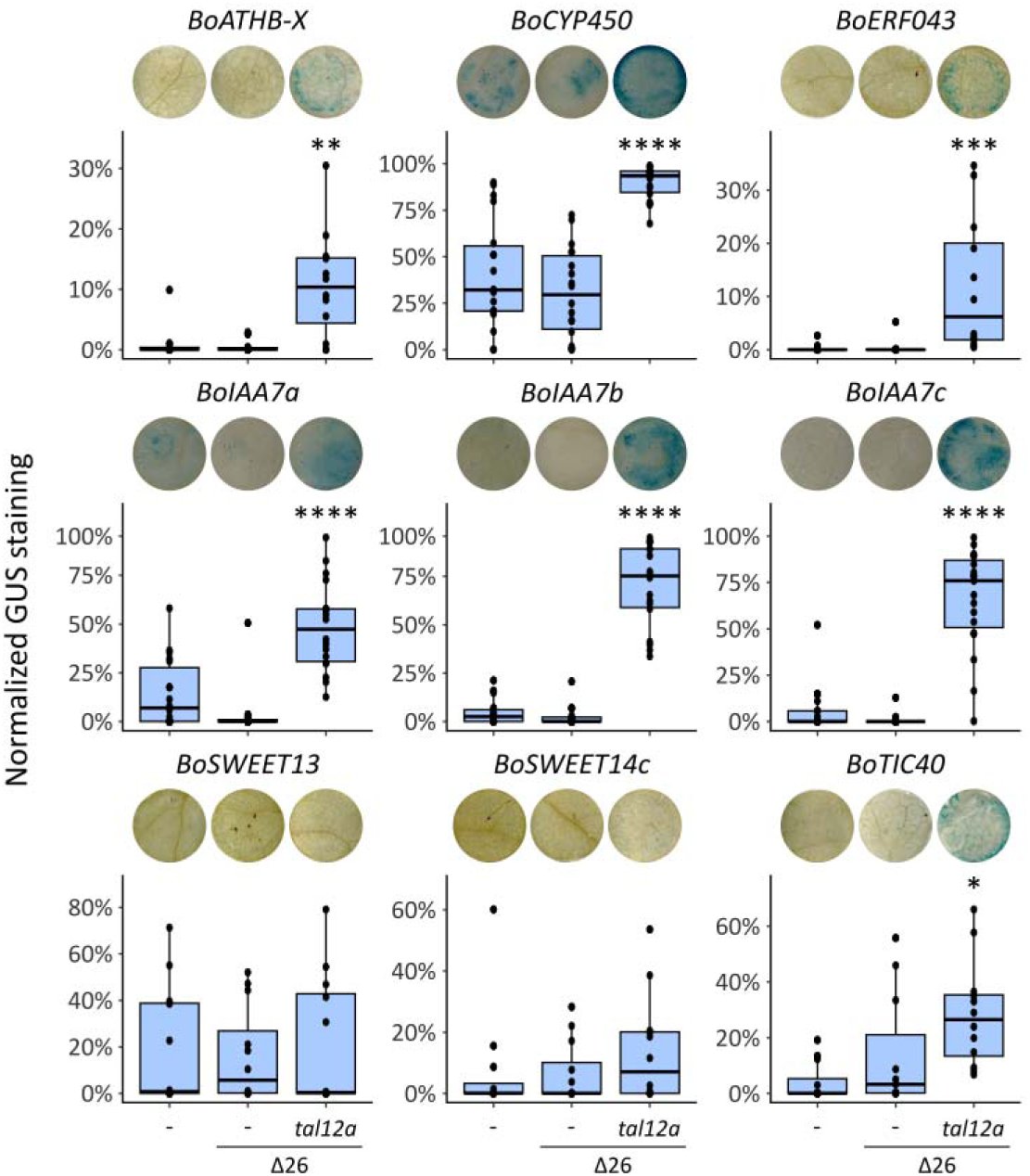
Tal12a-mediated transcriptional transactivation of candidate *cauliflower* susceptibility gene promoters in *N. benthamiana*. Promoter regions of nine Tal12a target genes (*BoCYP450*, *BoERF043*, *BoIAA7a*, *BoIAA7b*, *BoIAA7c*, *BoTIC40*, *BoSWEET13*, *BoSWEET14c*, and *BoATHB-X*) were fused to a *gus* reporter gene and delivered into *N. benthamiana* leaves via *Agrobacterium*-mediated transformation, either alone or in co-infiltration with the *Xcc* Δ26::*tal12a* strain or the Δ26 strain. GUS expression levels were quantified at 2 dpi using ImageJ. The experiment was independently repeated four times, with three plants per experiment, resulting in at least 11 data points per condition. Statistical significance was assessed using a Wilcoxon test, comparing the Δ26::*tal12a* condition to the Δ26 control for each promoter (*: p<0.05; **: p<0.01; ***: p<0.001)

### *BoIAA7c*, *BoSWEET13* and *BoSWEET14c* are minor *S* genes promoting disease symptoms in cauliflower

We assessed the contributions of *BoIAA7c*, *BoSWEET13*, and *BoSWEET14c* in cauliflower susceptibility to *Xcc* by testing whether their individual transcriptional activation by artificial TALEs (arTALEs) could mimic the virulence function of Tal12a. Two arTALEs were designed for each target promoter (Fig. **6a**, Table **S7**) and introduced into the Δ26 *Xcc* strain. To verify the functionality of the arTALEs, the transcript levels of *BoIAA7c*, *BoSWEET13*, and *BoSWEET14c* were quantified by RT-qPCR at 1 dpi in cauliflower leaves inoculated with Δ26 strains expressing the corresponding arTALE. As expected, plasmid-mediated expression of *tal12a* in the Δ26 strain (Δ26+Tal12a) significantly increased the transcript levels of all three candidate genes compared to the Δ26 carrying the empty vector (Δ26+EV) (Fig. **6b**). Consistent with their expected activity, each arTALE specifically targeting the promoter of *BoIAA7c*, *BoSWEET13*, or *BoSWEET14c* robustly induced expression of its target gene, reaching transcript levels comparable to or higher than those obtained with Tal12a (Fig. **6b**). Interestingly, arTALE13.2 also induced *BoSWEET14c* expression, although in silico analysis did not predict an EBE for arTALE13.2 in the *BoSWEET14c* promoter. To investigate the impact of arTALE-mediated activation of those three genes on disease development and bacterial fitness *in planta*, we used a simple mesophyll infiltration assay in cauliflower leaves. Symptoms were quantified using a disease index scale ranging from 0 to 4 (Fig. **S4**). At 7 dpi, the Δ26+Tal12a strain caused severe chlorosis and necrosis, confirming the contribution of Tal12a to virulence in this tissue, whereas the Δ26 strain carrying the empty vector (Δ26+EV) induced only mild chlorosis (Fig. **6c**). Interestingly, Δ26 strains expressing arTALE7.1, arTALE7.2, arTALE13.1, arTALE13.2, arTALE14.1, or arTALE14.2 also triggered more severe disease symptoms compared to the EV control. These results indicate that *BoIAA7c*, *BoSWEET13* and *BoSWEET14c* contribute to symptom development on cauliflower. However, no significant increase in bacterial proliferation was observed at 3 dpi with any of the arTALEs, in contrast to strain expressing Tal12a (Fig. **6d**). Notably, expression of arTALE7.1 was even associated with a reduction in bacterial population despite the increased symptom severity. The basis for this apparent uncoupling between symptom development and bacterial growth remains unclear. Taken together, these results provide evidence that *BoIAA7c*, *BoSWEET13* and *BoSWEET14c* contribute to the virulence of Tal12a in *Xcc*-cauliflower interactions.

**Fig. 6.**
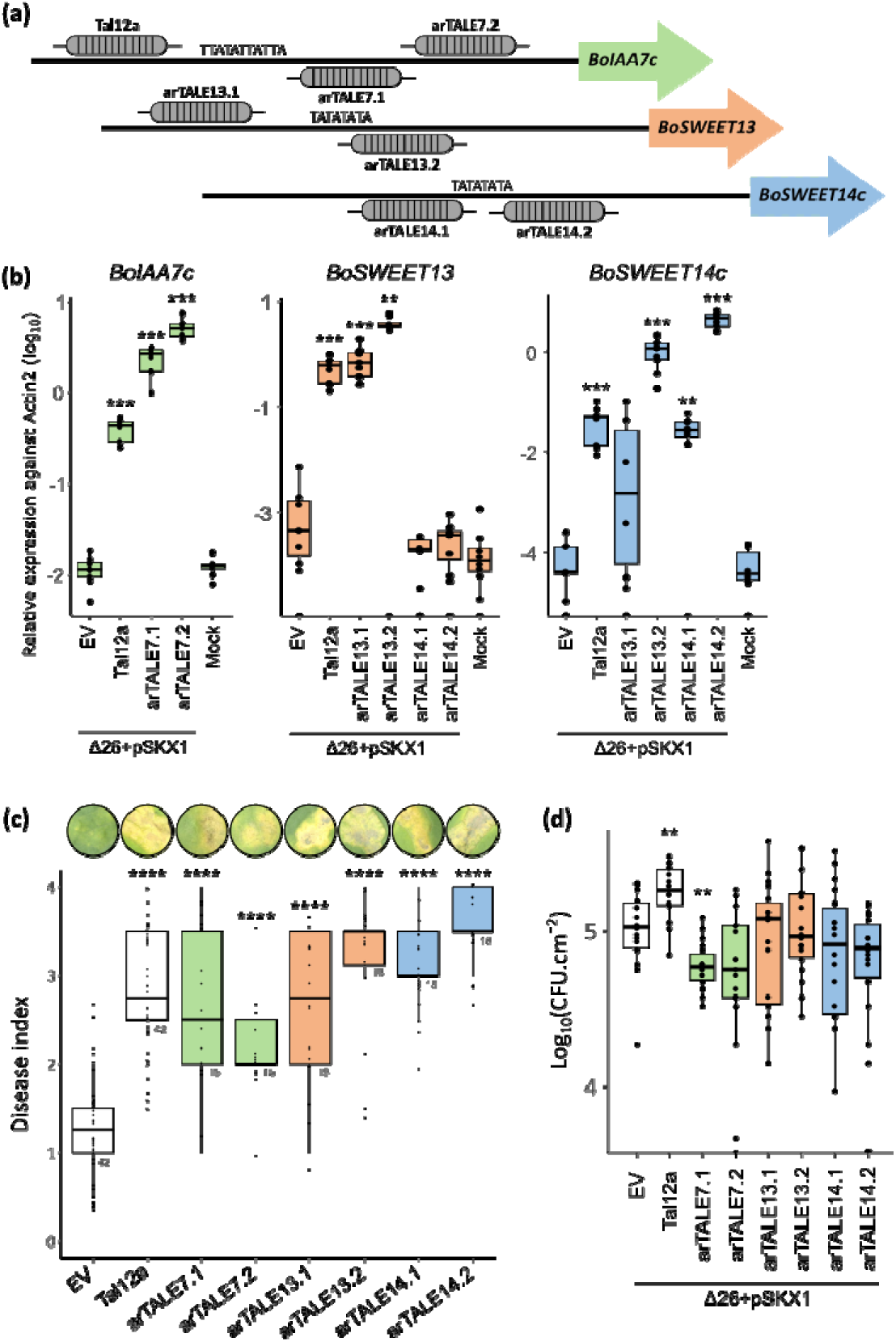
Tal12a and arTALE inducing expression of *BoIAA7c*, *BoSWEET13* and *BoSWEET14c* expression promote symptom development in cauliflower mesophyll. (**a**) ArTALE target sites designed based on the promoter sequences of *BoIAA7c*, *BoSWEET13* and *BoSWEET14c*. (**b**) Expression of *BoIAA7c*, *BoSWEET13* and *BoSWEET14c* were quantified by qRT-PCR in cauliflower leaves infiltrated with the Δ26 strain carrying either the empty vector pSKX1 (EV), the *tal12a* gene (+Tal12a) or individual arTALE constructs at 10 CFU·mL^-^¹ for each strain. Samples were collected at 1 dpi. The experiment was repeated three times independently, using three plants per replicate, yielding at least nine biological data points per condition. (**c**) Boxplot representation of disease symptoms at 7 dpi following infection with the *Xcc* Δ26 strain complemented with either the empty vector pSKX1 (EV), *tal12a* (+Tal12a) or arTALEs. Inoculations were performed on cauliflower leaves by syringe infiltration with bacterial suspensions at 10 CFU·mL^-^¹. Each condition was tested in six independent experiments using three plants per strain. The disease index is indicated in Fig. S4. (**d**) *Xcc* titers in cauliflower leaves were measured at 0 and 3 dpi following mesophyll infiltration with a bacterial suspension at 10^6^ CFU.mL^−1^. Three independent experiments were performed, each involving six plants per strain. Statistical significance in panels (**b**), **(c)** and (**d**) was assessed using a Wilcoxon test, comparing the different strain against the Δ26 + empty vector (EV) control (*: p<0.05; **: p<0.01; ***: p<0.001; ****: p<0,0001).

To explore a potential synergy between these three *S* genes, we co-delivered a 1:1 mix of Δ26 strains expressing arTALEIAA7.1 and either arTALE13.1, arTALE13.2, or arTALE14.2 in cauliflower mesophyll. Induction of the expected genes was confirmed for each combination of arTAL (Fig. **7a**). Co-inoculation of arTALEIAA7.1 and arTALE13.1, which induced simultaneous expression of *BoIAA7c* and *BoSWEET13* resulted in increased symptoms severity (DI = 3.5) compared with the empty vector control (DI = 1). Interestingly, co-expression of *BoIAA7c* with either *BoSWEET14c* alone (upon arTALEIAA7.1/arTALE14.2 co-delivery) or *BoSWEET14c* with *BoSWEET13* (upon arTALE7.1/arTALE13.2 co-delivery) was consistently associated with increased symptom severity relative to the empty vector, and to an even greater extent than with Tal12a (p = 1,6.10^-4^ and p = 1,74.10^-5^, respectively; Fig. **7b**). These results support the hypothesis that *BoSWEET13*, *BoSWEET14c* and *BoIAA7c* function as minor *S* genes, and that the co-activation of *BoIAA7* with either *BoSWEET13* or *BoSWEET14c* has an additive effect on disease symptom severity.

**Fig. 7.**
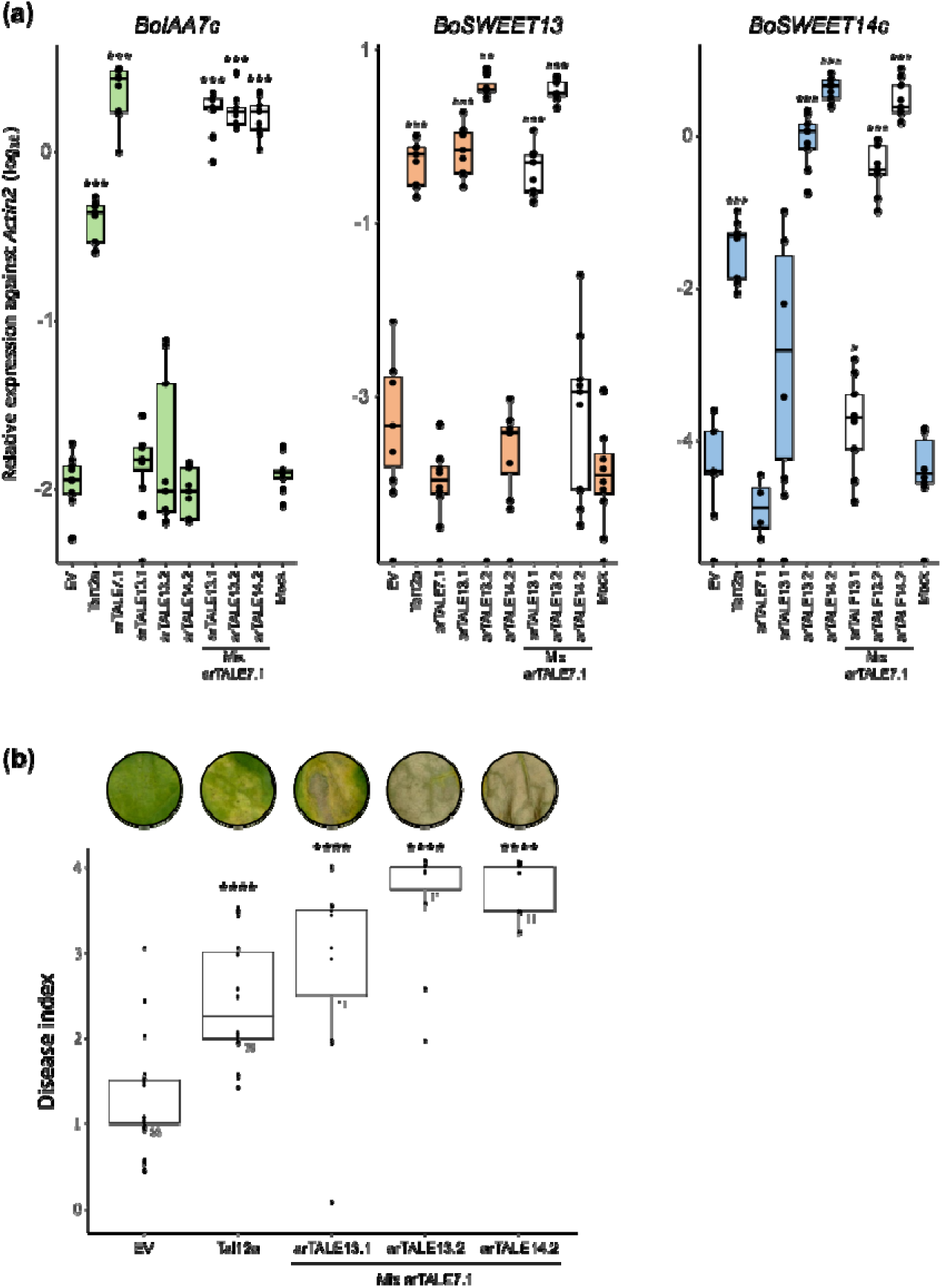
Co-induction of *BoIAA7c, BoSWEET13* and *BoSWEET14c* genes by arTALEs mix enhances disease symptom development in cauliflower mesophyll. (**a**) Gene expression of *BoIAA7c*, *BoSWEET13* and *BoSWEET14c* was assessed in the mesophyll one day after infiltration with *Xcc* suspensions (10^8^ CFU.mL^−1^) of Δ26 derivatives carrying either the empty vector pSKX1 (EV), the *tal12a* gene (+Tal12a) or a 1:1 mix of arTALE constructs. The experiment was repeated three times, with three plants per experiment, resulting in a minimum of 9 data points. (**b**) Boxplot representation of disease symptoms at 7 dpi by the Δ26 strain carrying either the empty vector pSKX1 (EV), the *tal12a* gene (+Tal12a) or a 1:1 mix of arTALE constructs. Five independent experiments were performed with one to three plant per strain. *Xcc* strains were inoculated by mesophyll infiltration of a cauliflower leaf (cv. Clovis) with a syringe filled in a bacterial suspension at 10^8^ CFU.mL^−1^. The disease index is indicated in Fig. S4. Statistical significances were assessed using a Wilcoxon test, comparing the different strain against the Δ26 + empty vector control (*: p<0.05; **: p<0.01; ***: p<0.001).

## DISCUSSION

### A detectable contribution of Tal12a to *Xcc* virulence on cauliflower

The first comprehensive analysis on *Xcc tal* genes was conducted in 2018 and identified Tal12a as a predominant TALE protein present in nine out of 38 strains representative of *Xcc* genetic diversity (Denancé *et al*., 2018). Tal12a contribution to virulence was previously shown only in combination with Tal15a or Tal22a on radish (Kay *et al*., 2005) and with Tal15a on cauliflower (Denancé *et al*., 2018). However, both studies failed to show any stand-alone Tal12a virulence functions. Here, we demonstrate that Tal12a alone promotes both bacterial proliferation and disease symptom development on cauliflower by gain-of-function in the *Xcc* strain 8004Δ26 as well as symptom development on cauliflower by loss-of-function in strain CN08. In contrast, Tal14c enhanced symptom development without affecting bacterial growth. These results thus illustrate that *Xcc tal* genes likely function as minor virulence determinants in contrast to other pathogens where major *tal* virulence functions have long been identified such as in *Xanthomonas oryzae* pv. *oryzae*, *Xanthomonas oryae* pv. *oryzicola*, *Xanthomonas citri* pv. *citri*, *Xanthomonas axonopodis* pv. *manihotis*, *Xanthomonas phaseoli* pv. *manihotis* (Streubel *et al*., 2013; Cernadas *et al*., 2014; Cohn *et al*., 2014; Hu *et al*., 2014; Cox *et al*., 2017). In *Xcc*, virulence seems to rely primarily on Xop type III effectors, like *XopAC* (Guy *et al*., 2013) or *XopJ6* (Lauber *et al*., 2024) while *tal* genes play minor yet significant roles in virulence.

### Tal12a induces specific transcriptional reprogramming while converging with Tal14c on shared downstream responses

Our transcriptomic analysis revealed that Tal12a alters processes beyond classical defense, including developmental and hormonal pathways such as megagametogenesis, ABA metabolism, and pollen development suggesting a specific effect of Tal12a different from Tal14c. Despite upregulating a distinct repertoire of direct target genes, Tal12a and Tal14c induce expression of a shared set of 43 genes, none of which appear to be direct targets of Tal12a. This overlap likely reflects indirect regulation through convergent downstream signaling pathways or common stress responses. Notably, GO enrichment analysis of these shared genes reveals categories associated with jasmonic acid signaling, wounding, and regulation of defense responses. This could reflect a case of functional convergence, where different TALEs manipulate distinct upstream regulators that ultimately feed into similar host processes. This scenario is reminiscent of TALEs within a single *Xoo* strain exhibiting functional redundancy by targeting the same genes or host pathways (Streubel *et al*., 2013; Doucouré *et al*., 2018).

### *BoIAA7c*, *BoSWEET13* and *BoSWEET14c* are minor *S* genes in cauliflower

Considering that Tal12a consists of only twelve repeats, it is theoretically more prone to bind and induce a larger number of genes. However, there were only 29 potential direct targets, both upregulated in the presence of Tal12a and harboring a candidate EBE in the promoter region. Among these, only one or few may act as *S* genes mediating Tal12a virulence functions, while others could be off-targets. We also identified two *SWEET* genes as indirect Tal12a targets. This contrasts with other cases of TALE-mediated *SWEET* induction in rice, cassava and cotton where activation is direct, via an EBE in the promoter (Zhang *et al*., 2022). Notably, TALE-independent induction of *SWEET* gene expression has also been reported upon infection by *Pseudomonas*, *Botrytis* and *Sinorhizobium* (Chen *et al*., 2010). The mechanisms by which these microbes naturally deprived of *tal* genes induce *SWEET* expression have not been characterized. These findings highlight the fundamental utility of *SWEET* gene upregulation as a common parasitic/symbiotic strategy. Our findings suggest that *Xcc* may have evolved an alternative TALE-dependent mechanism to induce *SWEET* gene expression. We demonstrated that *BoIAA7c*, *BoSWEET13* and *BoSWEET14c* function as minor *S* genes, contributing to symptom development without significantly enhancing bacterial growth. The precise role of *BoIAA7c* in disease progression remains unclear, although it is known that auxin signaling facilitates bacterial infection (Kunkel & Johnson, 2021). Interestingly the *Pseudomonas syringae* type III effector AvrRpt2 promotes the degradation of IAA27/AXR2 in Arabidopsis, thereby enhancing auxin signaling and suppressing plant immunity to facilitate infection (Cui *et al*., 2013). Thus, *BoIAA7c*, *BoSWEET13* and *BoSWEET14c* act as minor *S* genes, contributing quantitatively to susceptibility to *Xcc* in cauliflower, these genes might represent a useful resource for breeding durable resistance by loss-of-susceptibility.

### *BoIAA7c* with *BoSWEET13* or *BoSWEET14c* function additively to mediate Tal12a virulence functions in cauliflower

The individual induction of *BoIAA7c*, *BoSWEET13* and *BoSWEET14c* expression using arTALEs enhanced disease symptoms, without promoting bacterial proliferation. This indicates that neither gene accounts for virulence activity of Tal12a alone. This finding is noteworthy, as it diverges from the “one TALE/one *S* gene” model commonly described in *Xanthomonas*–plant interactions, in which the activation of a single *S* gene is typically necessary and sufficient to trigger strong disease phenotypes. A recent exception to this paradigm is Tal7b from *Xanthomonas citri* pv. *malvacearum*, which induces multiple functionally distinct targets (*GhSWEET14a*, *GhSWEET14b*, and *GhPL1*) that act additively to promote lesion development in cotton (Mormile *et al*., 2025). Similarly, our results suggest that Tal12a may act through a network of multiple *S* genes with complementary roles, pointing to a more complex and integrated virulence strategy than previously recognized. This observation raises the possibility that many uncharacterized TALEs contribute quantitatively to pathogenicity, acting in concert with other type III effectors to establish virulence. By elucidating these mechanisms, we pave the way for novel approaches to managing black rot disease in cruciferous crops, through the disruption of Tal12a-dependent susceptibility.

## MATERIALS AND METHODS

### Plant material

Cauliflower plants (*Brassica oleracea* cv. *botrytis* var. Clovis F1, Vilmorin) were grown in a greenhouse. *Nicotiana benthamiana* plants were sown on soil and grown at 24°C under long-day photoperiod (16-h light) with 60% relative humidity. Four-to-five weeks-old plants were used.

### Bacterial strains, plasmids and growth conditions

All bacterial strains used in this study are described in Table **S5**. *Xcc* strains were grown either in MOKA medium (yeast extract 4g/L, casamino acids 8g/L, K_2_HPO_4_ 2g/L, MgSO_4_.7H_2_O 0.3g/L) or in MME medium (K_2_HPO_4_ 10.5 g/L, KH_2_PO_4_ 4.5 g/L, (NH_4_)_2_SO_4_ 1 g/L, casamino acids 1.5g/L, MgSO_4_,7H_2_O 0.3g/L) with the appropriate antibiotics at 28°C. *Agrobacterium tumefaciens* GV3101 strains were grown in YEB medium (yeast extract 1g/L, beef extract 5g/L, peptone 5g/L, sucrose 5g/L, MgSO_4_ 0.4mM pH7.2) with the appropriate antibiotics at 28°C. CN08 wild type strain was isolated by He *et al*., (2007). Rifampicin-resistant and *tal12a* mutant derivatives of CN08 were obtained spontaneously. CN08Δ*tal12a* was complemented with *tal12a* cloned in pSKX1 (Tran *et al*., 2018). The Δ26 polymutant strain derivatives of *Xcc* strain 8004 lacks to coding regions of 26 type III effectors or secreted proteins (european patent EP20305940.7): *XopAC, AvrBs1, XopH, XopX1, XopX2, XopF, HrpW, XopD, XopAN, XopQ, XopK, XopJ5, XopG, AvrXccA1, AvrXccA2, XopAM, XopE2, XopAH, XopR, XopL, XopZ, XopAG, XopP, XopAL1, XopAL2* and *XopN*. TALE cloning vectors were created as described in Mormile *et al*. (2025). *tal* genes were integrated in the Δ26 strain genome between *XC_3385* and *XC_3386* genes as described in Luneau *et al*. (2022). TALE protein accumulation in all strains was assessed by Western blot analysis using a polyclonal antibody developed in rabbit against an *Xcc* TALE (Zárate Chaves *et al*., 2023). Rifampicin (50µg/mL), kanamicin (50µg/mL) and gentamicin (10 and 15µg/mL for *Xanthomonas* and *Agrobacterium*, respectively) were used for antibiotic concentrations.

### Virulence and *in planta* growth assays on cauliflower

Pathogenicity assays were conducted on the second true leaf by piercing the central vein using a needle dipped in a bacterial suspension at 10^8^ CFU/mL. Three to five independent biological repetitions were performed with five to six plants each time. Disease development was scored visually at 4, 7 or 10 dpi, with a disease index scale from 0 (no symptom) to 4 (V-shape like leaf necrosis), as described (Denancé *et al*., 2018).

For virulence assays, mesophyll infiltration was conducted on the second true leaf using a needleless syringe filled with a bacterial suspension at 10^8^ CFU/mL. At least four independent biological repetitions were performed with one to three plants each time. Symptoms were observed between 7 and 8 dpi according to a disease index scale from 0 to 4 (Fig. S3). *In planta* bacterial populations were determined after mesophyll infiltration of the second true leaf using a syringe filled with a bacterial suspension at 10^6^ CFU/mL. One leaf disc (6 mm diameter) was harvested at 0 and 3 dpi and ground with glass beads in 200µL 1mM MgCl_2_. The homogenate was serially diluted and spotted three times on MOKA plates supplemented with pimaricin (30µg/mL) and appropriate antibiotics. Plates were incubated at 28°C for 48h and colonies enumerated in spots containing 1 to 30 colonies. Bacterial densities in leaves were expressed as log CFU/cm^2^. Four independent biological repetitions were performed with six plants.

### *In vitro* bacterial growth assay

Bacterial growth of *Xcc* strains was monitored in MOKA and MME media by measuring optical density à 600nm over 24 hours. Growth rates were calculated during the exponential phase as described (Lauber *et al*., 2024). Three independent biological replicates were performed.

### Tissue sampling, RNA extraction and transcriptomic analyses

Mesophyll infiltrations were carried out on the second leaf of the plant, following previously described methods, using a bacterial suspension at 10□ CFU/mL. Fifteen leaf discs (7-mm diameter) were collected on three plants (five per plant) at 1 dpi and immediately frozen in liquid nitrogen. Three independent biological replicates were performed.

RNA was extracted using either the Macherey-Nagel NucleoSpin RNA Plus Kit or the Qiagen RNeasy Mini Plant Kit, following the manufacturer’s protocols. Global gene expression profiles were determined by oriented paired-end RNA sequencing (2x 150 bp) on an Illumina NovaSeq 6000 using the Illumina TruSeq Stranded mRNA Kit. Raw paired-end reads were processed using the *nf-core/rnaseq* pipeline version 3.0 (doi:10.5281/zenodo.4323183). We used the pseudo-mapping strategy with *salmon* software (version 1.4.0) and reference genome annotation of *Brassica oleracea* (https://ftp.ensemblgenomes.ebi.ac.uk/pub/plants/release-57/gff3/brassica_oleracea). Counts were retrieved at the gene level and were normalized using the EdgeR package and the CPM/TMM method (Robinson & Oshlack, 2010). After quality control, differential analysis was performed using fitted generalized linear models (GLMs) with a design matrix multiple factor (biological repetition and factor of interest). DEGs were called using the GLM likelihood ratio test using a false discovery rate (FDR Benjamini-Hochberg) adjusted p value < 0.01. GOterm were associated with the genes using Blast2GO (Conesa *et al*., 2005). Analysis of GO enrichment was conducted using the topGO package v. 2.52.0 (Alexa *et al*., 2006).

For quantitative real-time-PCR (qRT-PCR) analyses, DNase treatment was carried out using the Invitrogen™ TURBO DNA-free™ Kit, and cDNA was generated from 1µg RNA using the Transcriptor Reverse Transcriptase Kit by Roche. qRT-PCR was then conducted using the Takyon™ No ROX SYBR 2X MasterMix Blue dTTP Kit, following the manufacturer’s protocol. Primers for qRT-PCR used in this study are listed in Table **S6**.

### TALE target prediction

The promoter regions (1000 nt upstream the translation start site) of annotated genes in TO1000 were extracted from *B. oleracea* genome (see above) using the GenomicRanges package in R (Aboyoun *et al*., 2013). Predictions of binding sites were made using Talvez v3.2 (Pérez-Quintero *et al*., 2013) and Preditale (Erkes *et al*., 2019), the top 300 predicted targets according to score were kept for each RVD sequence.

### Promoter sequence cloning and transient co-expression assays in *N. benthamiana*

Promoter sequences were amplified using primers with *attB* sites (Table **S6**) and verified by sequencing (Table **S4**). The sequences were cloned into the pDONR207 vector using Gateway™ BP Clonase™ II Enzyme mix and transferred to the pJawohl11-GW-gus destination vector using Gateway™ LR Clonase™ II Enzyme mix, following the manufacturer’s instructions. The resulting plasmids were electroporated into *Agrobacterium tumefaciens* GV3101. For co-transient expression in *N*. *benthamiana*, *A. tumefaciens* and *Xanthomonas* strains were grown and resuspended in infiltration medium (10 mM MES pH 5.6, 10 mM MgCl_2_, 150 μM acetosyringone) to OD_600nm_ = 0.3 and OD_600nm_ = 0.1, respectively. After a 2 hours incubation at room temperature, the bacterial suspensions were infiltrated into the leaves of four-week-old *N. benthamiana* plants using a needleless syringe. Plants were incubated for two days in growth chambers. To assess GUS activity, 12 mm leaf discs were harvested from the inoculated leaves and vacuum infiltrated with a substrate-detergent solution containing 1 mM 5-bromo-4-chloro-3-indolyl β-D-glucuronide, 0.2% Triton X-100 [v/v], 2 mM K_3_FeCN_6_, 2 mM K_4_FeCN_6_, and 50 mM sodium phosphate buffer, pH 7.2. The discs were incubated overnight at 37 °C, after which they were fixed and destained in 80% ethanol. The intensity of the GUS staining was quantified using ImageJ software. The experiment included four independent biological replicates, each consisting of three plants.

### Construction of designer TAL effectors

The *BoSWEET13*, *BoSWEET14c* and *BoIAA7c* promoter sequences from *Brassica oleracea* were analyzed to design the suitable artificial TAL effectors (arTALE) binding sites as described previously in Streubel *et al*., 2014. The EBEs selected were verified for specificity using Talvez v.3.2 (https://webphim.ird.fr/cgi-bin/talvez/talvez.cgi;Pérez-Quintero et al., 2013). The arTALEs were generated using the Golden TAL technology and expressed under the control of a lac promoter in the Golden Gate-compatible broad host range vector pSKX1. The RVD sequences of the arTALEs used in this study are provided in Table S7.

## ACKNOWLEDGEMENTS

We are grateful to Boris Szurek (PHIM, Montpellier, France) for advice on the design and execution of the study.

## AUTHORS CONTRIBUTIONS

CA, LDN, IF and CB performed preliminary proof-of-concept tests. BC, ZED, AJB, LDN and CA participated in the experimental design and conception of the study. BC carried out all the experiments, performed the statistical analysis, BC and CA wrote the manuscript. CG and BC cloned the *Tal* genes and introduced them in the effectorless *Xanthomonas* strain. CB and IF isolated the CN08 mutant and complemented the strain. SC and BC analyzed and interpreted the transcriptomic data. ALPQ predicted the EBE. ALR performed the imageJ analysis. All co-authors reviewed and corrected the manuscript.

## FUNDING INFORMATIONS

The authors are grateful to the funding sources: B.C. was funded by a PhD grant from the INRAE BAP division. C.G., S.C., C.B., I.F. and L.D.N. were supported by the NEPHRON ANR project (ANR-18-CE20-0020-01). This study is set within the framework of the ‘Laboratoires d’Excellences’ (LABEX) TULIP (ANR-10-LABX-41) and of the ‘Ecole Universitaire de Recherche’ (EUR) TULIP-GS (ANR-18-EURE-0019). Z.D. was supported by a USDA NIFA Pre-doctoral Research Fellowship (2019-67011-29501).

## DATA AVAILABILITY

Raw sequence data for the transcriptomic analysis are available on the Sequence Read Archive (SRA) database under accession number SRP523615.

