## supplementary for "A *Xanthomonas* effector increases virulence quantitatively by inducing at least three minor susceptibility genes"

### SUPPORTING INFORMATION LEGENDS

**Fig. S1:** Identification of TALEs by western blot analysis

**Fig. S2:** Bacterial strains *in vitro* growth analysis

**Fig. S3:** Disease index of cauliflower leaves (cv. Clovis) inoculated with *Xcc* strains by mesophyll infiltration

**Table S1**: Properties of the RNAseq libraries in this study

**Table S2**: GO terms of cellular compartment and molecular function enriched among genes differentially expressed by Tal12a and Tal14c at one dpi

**Table S3:** EBE prediction analysis of Tal12a and Tal14c on cauliflower genome

**Table S4:** Promoter sequences of cauliflower genes used in this study

**Table S5:** Strain and plasmid used in the present study

**Table S6:** Primer used in the present study

**Table S7:** RVD sequences of the arTALEs used in the present study

**Table S8**: Top 300 genes upregulated by Tal12a and Tal14c in cauliflower identified by RNA-seq (FDR<0.01), ranked by log₂ fold change for each TALE.

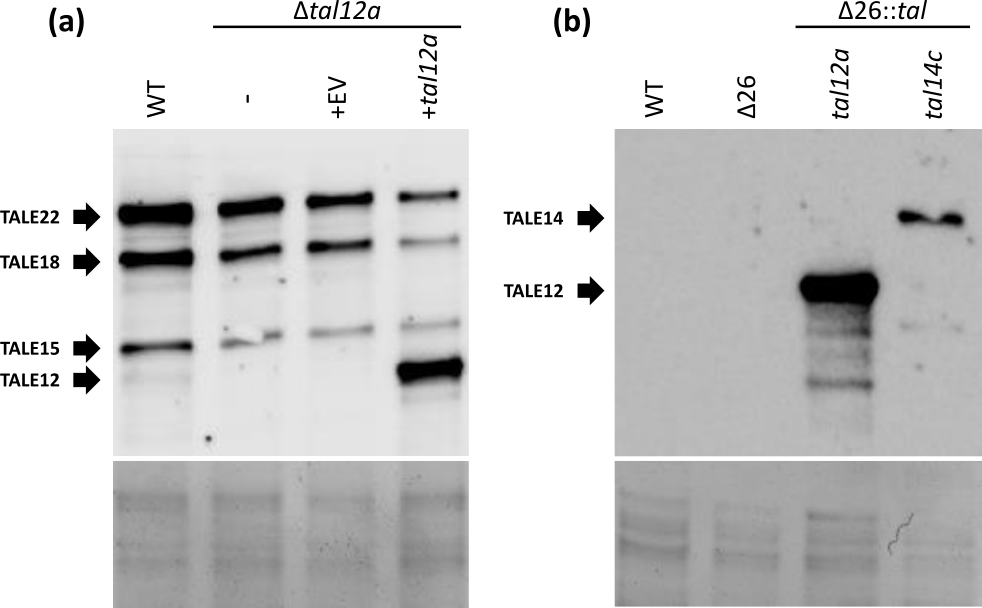

**Fig. S1:** Identification of TALEs by western blot analysis. Western blot analysis was performed using a polyclonal antibody developed against an *Xcc* TALE, as described in Zarate et al. (2023). (**a**) WT CN08 possesses four types of TALEs (Zarate-Chaves et al., 2023; Denancé et al., 2018). The upper panel shows chemiluminescent signals (anti-TALE) in the western blot assay, while the lower panel shows protein loading by Coomassie blue staining. The CN08Δ*tal12a* mutant, which arose spontaneously, was transformed with either the empty vector pSKX1 (EV) or with *tal12a* (+*tal12a*). (**b**) The Δ26 polymutant strain, derived from the 8004 strain, had 26 type III effectors deleted (European patent EP20305940.7). WT 8004 and Δ26 strains naturally carry no TALE gene. The *tal12a* and *tal14c* genes were integrated into the ∆26 strain, resulting in the Δ26::*tal12a* and Δ26::*tal14c* strains, respectively.

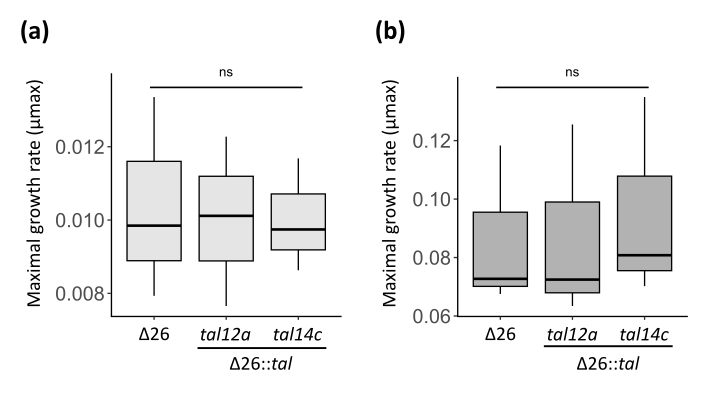

**Fig. S2:** Bacterial strains *in vitro* growth analysis. The Δ26 polymutant strain, derived from the 8004 strain, had 26 type III effectors deleted (European patent EP20305940.7). The ***tal12a*** and ***tal14c*** genes were integrated separately into the ∆26 strain, resulting in the Δ26::***tal12a*** and Δ26::***tal14c*** strains, respectively. Bacterial growth was assessed in both MME minimal medium (**a**) and MOKA rich medium (**b**). For each strain, a bacterial suspension at 1.5x10**^8^**CFU/mL was prepared, and growth was monitored by measuring the optical density at 600 nm (OD600) every 5 minutes over a 24-hour period at 28°C with continuous agitation (700 rpm). The maximum growth rate (µmax) was calculated during the exponential phase using the formula: Ln(2)*slope of growth curve. The experiment was performed with three independent biological replicates. Statistical groups were determined using the nonparametric Kruskal–Wallis test (p < 0.05) and are indicated by different letters.

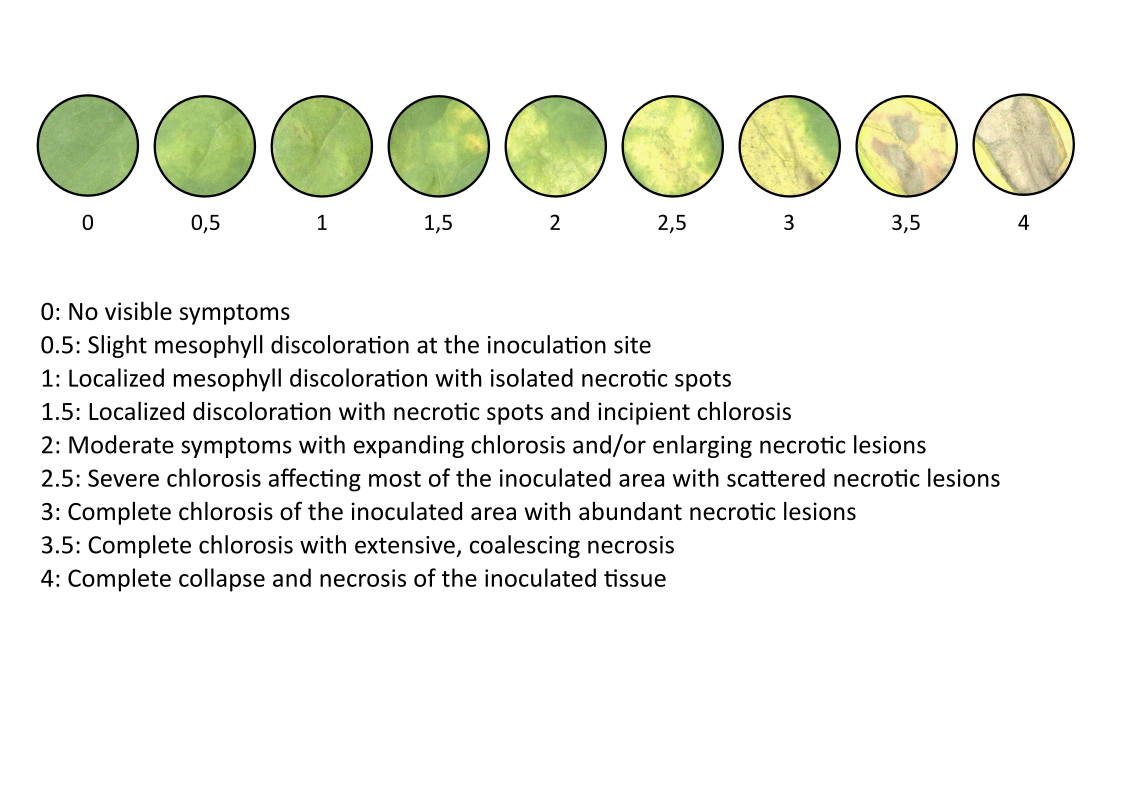

0: no symptoms

0,5: mesophyll decoloration of the infected area

1: mesophyll decoloration of the infected area with small necrotic lesions

1,5: mesophyll decoloration of the infected area with small necrotic lesions and chlorosis

2: mesophyll decoloration of the infected area with extended necrotic lesions or extended chlorosis

2,5: extended chlorosis with small necrotic lesions

3: full chlorosis of the infected area with necrotic lesions

3,5: full chlorosis of the infected area with extended necrosis

4: full necrosis of the infected area

**Fig. S3:** Disease index of cauliflower leaves (cv. Clovis) inoculated with *Xcc* strains by mesophyll infiltration. Symptoms were evaluated at 7–8 dpi and classified according to the disease index scale shown in the figure.

**Table S1**: Properties of the RNAseq libraries in this study

| **Strain** | **Number of raw reads per replicate** | **Mapped reads (%)^a^** | **Mapped reads (millions)^b^** | **Accession^c^** |
| --- | --- | --- | --- | --- |
| Δ26 | 16 068 039 | 34.6 | 5.5 | SRR30080690 |
|  | 16 957 946 | 35.8 | 6.0 | SRR30080692 |
|  | 16 775 385 | 35.5 | 5.9 | SRR30080694 |
| Δ26::*tal12a* | 15 288 503 | 39.0 | 5.9 | SRR30080684 |
|  | 15 699 430 | 35.4 | 5.5 | SRR30080686 |
|  | 14 146 883 | 34.9 | 4.9 | SRR30080688 |
| Δ26::*tal14c* | 14 404 324 | 35.4 | 5.1 | SRR30080678 |
|  | 21 340 180 | 33.8 | 7.2 | SRR30080680 |
|  | 14 814 835 | 35.7 | 5.2 | SRR30080682 |
| Mock | 15 523 042 | 33.6 | 5.2 | SRR30080696 |
|  | 16 337 941 | 34.1 | 5.5 | SRR30080698 |
|  | 16 420 080 | 33.6 | 5.5 | SRR30080700 |

^a^Percentage of reads successfully mapped to the TO1000 *Brassica* *oleracea* genome (<https://ftp.ensemblgenomes.ebi.ac.uk/pub/plants/release-57/gff3/brassica_oleracea>).

^b^Total number of reads in millions, that aligned to the reference *Brassica* *oleracea* genomes.

^c^Sequence Read Archive accession number.

**Table S2**: GO terms of cellular compartment and molecular function enriched among genes differentially expressed by Tal12a and Tal14c at one dpi

|  | **GO term** | **Term** | **Annotated^a^** | **Significant^b^** | **Expected^c^** | **Adjusted *p-*values** | **Fold enrichment^d^** |
| --- | --- | --- | --- | --- | --- | --- | --- |
| **Regulated genes by Tal12a only** | | |  |  |  |  |  |
| Cellular compartment | GO:0005618 | cell wall | 218 | 7 | 1,45 | 6,60E-04 | 4,4 |
|  | GO:0000418 | RNA polymerase IV complex | 18 | 2 | 0,12 | 6,24E-03 | 15,3 |
|  | GO:0000419 | RNA polymerase V complex | 18 | 2 | 0,12 | 6,24E-03 | 15,3 |
| Molecular function | GO:0010295 | (+)-abscisic acid 8'-hydroxylase activity | 9 | 3 | 0,06 | 1,90E-05 | 45,9 |
|  | GO:0042973 | glucan endo-1,3-beta-D-glucosidase activity | 95 | 6 | 0,58 | 2,80E-05 | 8,7 |
|  | GO:1990585 | hydroxyproline O-arabinosyltransferase activity | 5 | 2 | 0,03 | 3,70E-04 | 55,1 |
|  | GO:0004349 | glutamate 5-kinase activity | 5 | 2 | 0,03 | 3,70E-04 | 55,1 |
|  | GO:0004350 | glutamate-5-semialdehyde dehydrogenase activity | 5 | 2 | 0,03 | 3,70E-04 | 55,1 |
|  | GO:0016762 | xyloglucan:xyloglucosyl transferase activity | 58 | 4 | 0,36 | 4,60E-04 | 9,5 |
|  | GO:0008964 | phosphoenolpyruvate carboxylase activity | 26 | 3 | 0,16 | 5,40E-04 | 15,9 |
|  | GO:0061575 | cyclin-dependent protein serine/threonine kinase activator activity | 10 | 2 | 0,06 | 1,64E-03 | 27,5 |
|  | GO:0047372 | acylglycerol lipase activity | 17 | 2 | 0,1 | 4,83E-03 | 16,2 |
|  | GO:0000976 | transcription cis-regulatory region binding | 1298 | 16 | 7,99 | 6,88E-03 | 1,7 |
|  | GO:0003680 | minor groove of adenine-thymine-rich DNA binding | 65 | 3 | 0,4 | 7,59E-03 | 6,4 |
| **Regulated genes by Tal14c only** | | |  |  |  |  |  |
| Biological process | GO:0009611 | response to wounding | 229 | 25 | 3,3 | 5,50E-15 | 7,4 |
|  | GO:0031408 | oxylipin biosynthetic process | 45 | 12 | 0,65 | 1,30E-12 | 18,0 |
|  | GO:2000022 | regulation of jasmonic acid mediated signaling pathway | 57 | 11 | 0,82 | 5,10E-10 | 13,0 |
|  | GO:0009695 | jasmonic acid biosynthetic process | 25 | 8 | 0,36 | 1,50E-09 | 21,6 |
|  | GO:0000162 | tryptophan biosynthetic process | 58 | 10 | 0,84 | 9,90E-09 | 11,6 |
|  | GO:0010951 | negative regulation of endopeptidase activity | 135 | 13 | 1,95 | 8,80E-08 | 6,5 |
|  | GO:0051923 | sulfation | 51 | 8 | 0,74 | 6,50E-07 | 10,6 |
|  | GO:0016126 | sterol biosynthetic process | 91 | 9 | 1,31 | 6,90E-06 | 6,7 |
|  | GO:0009753 | response to jasmonic acid | 263 | 24 | 3,79 | 8,30E-06 | 6,2 |
|  | GO:0016102 | diterpenoid biosynthetic process | 100 | 9 | 1,44 | 1,50E-05 | 6,1 |
|  | GO:0006572 | tyrosine catabolic process | 23 | 5 | 0,33 | 1,70E-05 | 14,7 |
|  | GO:0070814 | hydrogen sulfide biosynthetic process | 13 | 4 | 0,19 | 2,80E-05 | 20,8 |
|  | GO:0031347 | regulation of defense response | 297 | 14 | 4,28 | 1,20E-04 | 3,2 |
|  | GO:0009407 | toxin catabolic process | 56 | 6 | 0,81 | 1,50E-04 | 7,2 |
|  | GO:0006563 | L-serine metabolic process | 58 | 6 | 0,84 | 1,90E-04 | 7,0 |
|  | GO:0002213 | defense response to insect | 23 | 4 | 0,33 | 3,00E-04 | 11,7 |
|  | GO:0019761 | glucosinolate biosynthetic process | 42 | 5 | 0,61 | 3,30E-04 | 8,0 |
|  | GO:0009873 | ethylene-activated signaling pathway | 159 | 9 | 2,29 | 5,30E-04 | 3,8 |
|  | GO:0000105 | histidine biosynthetic process | 48 | 5 | 0,69 | 6,30E-04 | 7,0 |
|  | GO:0009651 | response to salt stress | 521 | 18 | 7,51 | 6,50E-04 | 2,3 |
|  | GO:0006633 | fatty acid biosynthetic process | 303 | 20 | 4,37 | 6,80E-04 | 4,5 |
|  | GO:0009414 | response to water deprivation | 402 | 15 | 5,8 | 8,30E-04 | 2,5 |
|  | GO:0042343 | indole glucosinolate metabolic process | 14 | 3 | 0,2 | 9,60E-04 | 14,5 |
|  | GO:0009850 | auxin metabolic process | 90 | 8 | 1,3 | 1,04E-03 | 6,0 |
|  | GO:0015706 | nitrate transmembrane transport | 54 | 5 | 0,78 | 1,08E-03 | 6,2 |
|  | GO:0000103 | sulfate assimilation | 32 | 4 | 0,46 | 1,11E-03 | 8,4 |
|  | GO:1902358 | sulfate transmembrane transport | 32 | 4 | 0,46 | 1,11E-03 | 8,4 |
|  | GO:0009070 | serine family amino acid biosynthetic process | 81 | 6 | 1,17 | 1,14E-03 | 5,0 |
|  | GO:0031407 | oxylipin metabolic process | 49 | 14 | 0,71 | 1,16E-03 | 19,3 |
|  | GO:0009116 | nucleoside metabolic process | 56 | 5 | 0,81 | 1,27E-03 | 6,0 |
|  | GO:0051865 | protein autoubiquitination | 17 | 3 | 0,25 | 1,74E-03 | 11,9 |
|  | GO:1990961 | xenobiotic detoxification by transmembrane export across the plasma membrane | 123 | 7 | 1,77 | 2,10E-03 | 3,8 |
|  | GO:0080028 | nitrile biosynthetic process | 19 | 3 | 0,27 | 2,43E-03 | 10,7 |
|  | GO:0009694 | jasmonic acid metabolic process | 66 | 12 | 0,95 | 2,68E-03 | 12,3 |
|  | GO:1901606 | alpha-amino acid catabolic process | 193 | 13 | 2,78 | 3,20E-03 | 4,5 |
|  | GO:0050832 | defense response to fungus | 419 | 14 | 6,04 | 3,36E-03 | 2,3 |
|  | GO:1900057 | positive regulation of leaf senescence | 22 | 3 | 0,32 | 3,74E-03 | 9,2 |
|  | GO:0006522 | alanine metabolic process | 22 | 3 | 0,32 | 3,74E-03 | 9,2 |
|  | GO:0000096 | sulfur amino acid metabolic process | 103 | 6 | 1,49 | 3,84E-03 | 3,9 |
|  | GO:0034440 | lipid oxidation | 107 | 6 | 1,54 | 4,63E-03 | 3,8 |
|  | GO:1902652 | secondary alcohol metabolic process | 24 | 3 | 0,35 | 4,81E-03 | 8,4 |
|  | GO:0034599 | cellular response to oxidative stress | 108 | 6 | 1,56 | 4,84E-03 | 3,7 |
|  | GO:0002239 | response to oomycetes | 146 | 7 | 2,11 | 5,41E-03 | 3,2 |
|  | GO:0042762 | regulation of sulfur metabolic process | 25 | 3 | 0,36 | 5,41E-03 | 8,1 |
|  | GO:0019762 | glucosinolate catabolic process | 49 | 4 | 0,71 | 5,42E-03 | 5,5 |
|  | GO:0006730 | one-carbon metabolic process | 78 | 5 | 1,12 | 5,44E-03 | 4,3 |
|  | GO:0043455 | regulation of secondary metabolic process | 79 | 5 | 1,14 | 5,74E-03 | 4,3 |
|  | GO:0006979 | response to oxidative stress | 562 | 20 | 8,1 | 6,19E-03 | 2,4 |
|  | GO:0006520 | amino acid metabolic process | 943 | 47 | 13,6 | 6,24E-03 | 3,4 |
|  | GO:0009063 | amino acid catabolic process | 189 | 8 | 2,73 | 6,37E-03 | 2,9 |
|  | GO:0009901 | anther dehiscence | 27 | 3 | 0,39 | 6,74E-03 | 7,5 |
|  | GO:0009395 | phospholipid catabolic process | 29 | 3 | 0,42 | 8,24E-03 | 7,0 |
|  | GO:0009852 | auxin catabolic process | 10 | 2 | 0,14 | 8,65E-03 | 13,5 |
|  | GO:0036444 | calcium import into the mitochondrion | 10 | 2 | 0,14 | 8,65E-03 | 13,5 |
|  | GO:0042221 | response to chemical | 3691 | 112 | 53,22 | 9,71E-03 | 2,0 |
| Cellular compartment | GO:0005775 | vacuolar lumen | 6 | 2 | 0,08 | 2,40E-03 | 22,5 |
|  | GO:0010168 | ER body | 7 | 2 | 0,09 | 3,40E-03 | 19,3 |
|  | GO:1990246 | uniplex complex | 10 | 2 | 0,13 | 7,10E-03 | 13,5 |
| Molecular function | GO:0005506 | iron ion binding | 713 | 50 | 11,2 | 1,50E-18 | 4,7 |
|  | GO:0020037 | heme binding | 755 | 47 | 11,86 | 1,60E-15 | 4,2 |
|  | GO:0004497 | monooxygenase activity | 638 | 46 | 10,02 | 2,20E-14 | 4,9 |
|  | GO:0016705 | oxidoreductase activity, acting on paired donors, with incorporation or reduction of molecular oxygen | 795 | 49 | 12,49 | 2,90E-10 | 4,2 |
|  | GO:0046423 | allene-oxide cyclase activity | 6 | 5 | 0,09 | 5,60E-09 | 56,2 |
|  | GO:0008146 | sulfotransferase activity | 81 | 11 | 1,27 | 5,90E-08 | 9,2 |
|  | GO:0016702 | oxidoreductase activity, acting on single donors with incorporation of molecular oxygen, incorporation of two atoms of oxygen | 72 | 10 | 1,13 | 1,90E-07 | 9,4 |
|  | GO:0004866 | endopeptidase inhibitor activity | 138 | 13 | 2,17 | 3,10E-07 | 6,4 |
|  | GO:0030170 | pyridoxal phosphate binding | 210 | 15 | 3,3 | 1,40E-06 | 4,8 |
|  | GO:0010333 | terpene synthase activity | 70 | 9 | 1,1 | 1,50E-06 | 8,7 |
|  | GO:0008113 | peptide-methionine (S)-S-oxide reductase activity | 15 | 5 | 0,24 | 2,50E-06 | 22,5 |
|  | GO:0036456 | L-methionine-(S)-S-oxide reductase activity | 9 | 4 | 0,14 | 7,10E-06 | 30,0 |
|  | GO:0004031 | aldehyde oxidase activity | 20 | 5 | 0,31 | 1,20E-05 | 16,9 |
|  | GO:0005544 | calcium-dependent phospholipid binding | 21 | 5 | 0,33 | 1,60E-05 | 16,1 |
|  | GO:0004838 | L-tyrosine:2-oxoglutarate aminotransferase activity | 21 | 5 | 0,33 | 1,60E-05 | 16,1 |
|  | GO:0120091 | jasmonic acid hydrolase | 5 | 3 | 0,08 | 3,80E-05 | 40,5 |
|  | GO:0000287 | magnesium ion binding | 250 | 14 | 3,93 | 4,80E-05 | 3,8 |
|  | GO:0016207 | 4-coumarate-CoA ligase activity | 14 | 4 | 0,22 | 5,30E-05 | 19,3 |
|  | GO:0004020 | adenylylsulfate kinase activity | 15 | 4 | 0,24 | 7,20E-05 | 18,0 |
|  | GO:0051213 | dioxygenase activity | 271 | 24 | 4,26 | 1,20E-04 | 6,0 |
|  | GO:0016161 | beta-amylase activity | 20 | 4 | 0,31 | 2,40E-04 | 13,5 |
|  | GO:0102229 | amylopectin maltohydrolase activity | 20 | 4 | 0,31 | 2,40E-04 | 13,5 |
|  | GO:0004834 | tryptophan synthase activity | 23 | 4 | 0,36 | 4,20E-04 | 11,7 |
|  | GO:0019871 | sodium channel inhibitor activity | 10 | 3 | 0,16 | 4,30E-04 | 20,2 |
|  | GO:0071949 | FAD binding | 149 | 9 | 2,34 | 6,10E-04 | 4,1 |
|  | GO:0004617 | phosphoglycerate dehydrogenase activity | 12 | 3 | 0,19 | 7,60E-04 | 16,9 |
|  | GO:0008171 | O-methyltransferase activity | 97 | 7 | 1,52 | 8,60E-04 | 4,9 |
|  | GO:0004364 | glutathione transferase activity | 97 | 7 | 1,52 | 8,60E-04 | 4,9 |
|  | GO:0008483 | transaminase activity | 163 | 13 | 2,56 | 1,80E-03 | 5,4 |
|  | GO:0008172 | S-methyltransferase activity | 17 | 3 | 0,27 | 2,23E-03 | 11,9 |
|  | GO:0010179 | IAA-Ala conjugate hydrolase activity | 5 | 2 | 0,08 | 2,39E-03 | 27,0 |
|  | GO:0000254 | C-4 methylsterol oxidase activity | 18 | 3 | 0,28 | 2,64E-03 | 11,2 |
|  | GO:0045543 | gibberellin 2-beta-dioxygenase activity | 19 | 3 | 0,3 | 3,10E-03 | 10,7 |
|  | GO:0016844 | strictosidine synthase activity | 39 | 4 | 0,61 | 3,21E-03 | 6,9 |
|  | GO:0070704 | sterol desaturase activity | 6 | 2 | 0,09 | 3,54E-03 | 22,5 |
|  | GO:0003849 | 3-deoxy-7-phosphoheptulonate synthase activity | 6 | 2 | 0,09 | 3,54E-03 | 22,5 |
|  | GO:0016830 | carbon-carbon lyase activity | 229 | 10 | 3,6 | 3,57E-03 | 2,9 |
|  | GO:0016709 | oxidoreductase activity, acting on paired donors, with incorporation or reduction of molecular oxygen, NAD(P)H as one donor, and incorporation of one atom of oxygen | 127 | 7 | 2 | 4,03E-03 | 3,7 |
|  | GO:0004013 | adenosylhomocysteinase activity | 7 | 2 | 0,11 | 4,91E-03 | 19,3 |
|  | GO:0051753 | mannan synthase activity | 45 | 4 | 0,71 | 5,40E-03 | 6,0 |
|  | GO:0016491 | oxidoreductase activity | 3020 | 124 | 47,44 | 6,04E-03 | 2,8 |
|  | GO:0008271 | secondary active sulfate transmembrane transporter activity | 25 | 3 | 0,39 | 6,86E-03 | 8,1 |
|  | GO:0017057 | 6-phosphogluconolactonase activity | 9 | 2 | 0,14 | 8,24E-03 | 15,0 |
|  | GO:0019137 | thioglucosidase activity | 111 | 6 | 1,74 | 8,28E-03 | 3,6 |
|  | GO:0102799 | glucosinolate glucohydrolase activity | 111 | 6 | 1,74 | 8,28E-03 | 3,6 |
|  | GO:0015112 | nitrate transmembrane transporter activity | 51 | 4 | 0,8 | 8,41E-03 | 5,3 |
|  | GO:0004806 | triglyceride lipase activity | 27 | 3 | 0,42 | 8,52E-03 | 7,5 |
|  | GO:0042910 | xenobiotic transmembrane transporter activity | 149 | 7 | 2,34 | 9,45E-03 | 3,2 |
| **Regulated genes by both Tal12a and Tal14c** | | |  |  |  |  |  |
| Cellular compartment | GO:0005576 | extracellular region | 1579 | 6 | 1,16 | 7,90E-04 | 0,5 |
|  | GO:0005764 | lysosome | 83 | 1 | 0,06 | 5,92E-02 | 1,5 |
|  | GO:0000323 | lytic vacuole | 89 | 1 | 0,07 | 6,33E-02 | 1,4 |
|  | GO:0005783 | endoplasmic reticulum | 1409 | 3 | 1,03 | 8,24E-02 | 0,3 |
| Molecular function | GO:0071949 | FAD binding | 149 | 5 | 0,12 | 1,10E-07 | 4,2 |
|  | GO:0016491 | oxidoreductase activity | 3020 | 12 | 2,36 | 1,20E-06 | 0,5 |
|  | GO:0120091 | jasmonic acid hydrolase | 5 | 1 | 0 | 3,90E-03 | 25,1 |
|  | GO:0008757 | S-adenosylmethionine-dependent methyltransferase activity | 544 | 3 | 0,43 | 8,70E-03 | 0,7 |

Note: The lowercase letters a specified

^a^ Number of annoted genes for a GO term,

^b^ Number of up-regulated regulated genes among the genes annotated for a GO term,

^c^ Expected number of up-regulated genes if no significant enrichment,

^d^ Enrichment of the term in the Up-regulated gene list compared to the background population of genes,

**Table S3:** EBE prediction analysis of Tal12a and Tal14c on cauliflower genome

|  |  | | **TALVEZ software** | | | | **PrediTALE software** | | | |
| --- | --- | --- | --- | --- | --- | --- | --- | --- | --- | --- |
| **Gene ID** | **log2FC** | **FDR** | **EBE score** | **EBE rank** | **EBE strand** | **EBE position** | **EBE score** | **EBE rank** | **EBE strand** | **EBE position** |
| **Tal12a EBE prediction analysis** | | | | | | | | | | |
| Bo3g023170 | 8,89 | 1,5E-99 | 10,8 | 33 | + | -150 | 0,55 | 125 | + | 851 |
| Bo3g171090 | 6,55 | 1,5E-27 |  |  |  |  | 0,54 | 176 | + | 896 |
| Bo4g025620 | 6,50 | 5,4E-41 |  |  |  |  | 0,54 | 165 | + | 928 |
| Bo6g112570 | 6,37 | 3,9E-113 | 9,9 | 158 | + | -135 | 0,54 | 163 | + | 866 |
| Bo3g009500 | 5,70 | 6,4E-23 | 9,7 | 231 | - | -798 | 0,54 | 191 | - | 190 |
| Bo3g002110 | 5,48 | 2,8E-78 | 10,3 | 83 | + | -758 |  |  |  |  |
| Bo1g103470 | 4,80 | 8,8E-44 | 9,7 | 207 | + | -101 |  |  |  |  |
| Bo5g093110 | 4,69 | 8,1E-68 | 9,7 | 216 | + | -120 |  |  |  |  |
| Bo5g148490 | 4,33 | 4,8E-37 |  |  |  |  | 0,54 | 192 | + | 309 |
| Bo2g013210 | 4,07 | 2,2E-64 | 10,2 | 96 | + | -110 | 0,55 | 153 | + | 891 |
| Bo7g056390 | 3,85 | 2,2E-11 | 10,9 | 23 | + | -235 | 0,61 | 9 | + | 766 |
| Bo3g081600 | 3,78 | 4,6E-18 | 9,7 | 212 | + | -116 |  |  |  |  |
| Bo9g102490 | 3,76 | 4,8E-39 |  |  |  |  | 0,54 | 295 | + | 956 |
| Bo4g125880 | 3,61 | 4,4E-07 |  |  |  |  | 0,54 | 278 | + | 834 |
| Bo4g059980 | 3,53 | 7,8E-40 | 9,9 | 132 | + | -280 |  |  |  |  |
| Bo7g110790 | 3,43 | 3,2E-08 | 9,6 | 293 | - | -611 |  |  |  |  |
| Bo8g042120 | 3,26 | 2,5E-05 |  |  |  |  | 0,54 | 257 | + | 941 |
| Bo3g055640 | 3,19 | 2,6E-39 |  |  |  |  | 0,54 | 189 | + | 885 |
| Bo9g163920 | 3,10 | 8,8E-04 | 11,2 | 15 | + | -235 | 0,55 | 88 | + | 766 |
| Bo5g123740 | 2,78 | 1,5E-30 | 10,2 | 102 | + | -278 | 0,54 | 170 | + | 723 |
| Bo4g098840 | 2,77 | 2,5E-34 | 9,9 | 133 | + | -204 |  |  |  |  |
| Bo6g047940 | 2,74 | 2,7E-07 | 10,9 | 21 | - | -360 | 0,60 | 22 | - | 628 |
| Bo1g129910 | 2,69 | 7,6E-19 | 11,5 | 2 | - | -59 | 0,61 | 16 | - | 929 |
| Bo9g166010 | 2,64 | 7,9E-03 | 9,6 | 298 | + | -156 |  |  |  |  |
| Bo7g049930 | 2,59 | 4,6E-05 | 10,4 | 79 | + | -82 | 0,59 | 28 | + | 919 |
| Bo6g018460 | 2,59 | 6,5E-09 | 11,5 | 3 | + | -924 | 0,61 | 7 | + | 77 |
| Bo3g035870 | 2,41 | 1,2E-05 | 9,7 | 210 | - | -670 |  |  |  |  |
| Bo5g127080 | 2,39 | 1,2E-30 | 9,7 | 243 | + | -274 |  |  |  |  |
| Bo8g097350 | 2,10 | 7,7E-16 | 10,2 | 104 | + | -68 | 0,54 | 171 | + | 933 |
| **Tal14c EBE prediction analysis** | | | | | | | | | | |
| Bo2g023630 | 8,07 | 3,0E-85 | 12,2 | 34 | + | -209 | 0,43 | 22 | + | 792 |
| Bo6g006880 | 7,74 | 2,5E-60 | 12,2 | 39 | - | -70 | 0,43 | 19 | - | 916 |
| Bo8g077060 | 7,11 | 4,4E-60 | 11,0 | 194 | + | -51 | 0,42 | 28 | + | 950 |
| Bo3g123620 | 5,93 | 2,2E-26 | 11,5 | 94 | - | -138 | 0,40 | 89 | - | 848 |
| Bo2g127540 | 5,55 | 1,1E-28 | 11,5 | 99 | + | -313 | 0,37 | 201 | + | 688 |
| Bo01561s020 | 4,55 | 3,0E-09 | 13,1 | 19 | + | -739 | 0,44 | 14 | + | 262 |
| Bo2g064710 | 4,52 | 3,2E-22 | 11,7 | 68 | - | -324 |  |  |  |  |
| Bo5g108630 | 4,23 | 5,8E-45 |  |  |  |  | 0,38 | 193 | + | 917 |
| Bo6g051490 | 3,95 | 5,8E-50 | 11,9 | 48 | + | -101 | 0,39 | 104 | + | 900 |
| Bo8g106320 | 3,82 | 1,7E-25 |  |  |  |  | 0,40 | 97 | + | 916 |
| Bo3g027380 | 3,21 | 1,8E-29 |  |  |  |  | 0,45 | 13 | + | 970 |
| Bo6g076980 | 3,07 | 8,7E-05 |  |  |  |  | 0,37 | 217 | + | 532 |
| Bo5g126880 | 2,95 | 3,9E-04 | 10,8 | 221 | - | -623 |  |  |  |  |
| Bo8g095240 | 2,88 | 4,4E-04 | 11,1 | 182 | + | -40 | 0,42 | 46 | + | 961 |
| Bo7g114290 | 2,74 | 6,6E-20 | 11,2 | 141 | + | -157 |  |  |  |  |
| Bo2g056230 | 2,70 | 1,6E-28 | 10,5 | 299 | - | -936 |  |  |  |  |
|  | 2,70 | 1,6E-28 | 10,5 | 300 | - | -943 |  |  |  |  |
| Bo1g117730 | 2,63 | 4,3E-03 |  |  |  |  | 0,41 | 58 | + | 437 |
| Bo5g115220 | 2,29 | 5,0E-04 |  |  |  |  | 0,39 | 106 | + | 636 |
|  | 2,29 | 5,0E-04 |  |  |  |  | 0,37 | 295 | + | 454 |
| Bo3g073020 | 2,16 | 1,5E-10 |  |  |  |  | 0,39 | 117 | + | 294 |
| Bo5g008030 | 2,14 | 2,8E-12 | 10,6 | 286 | - | -635 |  |  |  |  |

**Table S4:** Promoter sequences of cauliflower genes used in this study

| **Name** | **Gene** | **Promoter sequence** | **Length** |
| --- | --- | --- | --- |
| *BoSWEET13* | Bo9g097170 | CAGCACAGTTCGCTTTTAGTGTGGGGCTCTGATGGAATCGTTAAGCTCGATCAACTCCCCCTCCTCGCTAATTAGCTATTAGGGAATCAAAAACTTTAAAAGATGATTTGAAGTAAATCAAAATCATAGTGATCTATACTTCTTTAATTGGTAATTAGCTAGAAATATAGCAGCCGAACAACGGGCTTATTGGTTACGCCGCGAAAATACTAAGTAACCACTCATTAGTGACTTTAATAAAACCTCTATCGACCACTCTTTATTCCAATTTCTTTTATTTTTTGTTTTGTTTCTTAAAGAAAGTAATTTTTAGATCATTCGTATGAGTACGATACTACAAATATAAACATAGCTTTGTGGTATAAAGTTTGATTCTAATTTAATAGACACCAAGAGTGCTGCCACATGAAAATATAATACAACTTTTATTTAAAAGAAAACATCAACTGAAAATATCTGAGACCAATAATGTGACCTGTCCTTCTCTCATTACTCTGGAAACGATGTCGTTCTAAACATATTAACACTACACCCCTTCACATGTGAACCCTCTATATATATCTGCTATGGTTGGCATTGTTTCATAGCTAAAACTCACAAAACTAGAGCTTGGCTTTCAAGAGAAAGAAGAGTCTTCAAGAGCTCTATCTCGCTCTTTCACTTGGTCTTAGATATTTCTTTTCCCGTTCATAGTTTTTGTCAATAATCCGACCAAAAAGGAGAGAGAAATAATCAAGAAAGCAGAAACAA | 750 bp |
| *BoSWEET14c* | Bo7g109890 | TCTACCATGCAAACTTTGGAATAGTAAAAAGGAAAATTTTGAAGTATGAATCCAGAAATCATAAGTTTTGAATATATATATATATATATATATGTATGTGTAGGTAAACAAAAATATAAACTATATATTTGGCTAGTAGAATACTGGAAACTTACATGACTGTATAAAACTCGTGAGTCGTGACCAAACGTAGTATTATGGAATATATAGCCATAATCGATATTTATTTGAATGTTTGGCTAGAAAACTAACAGCTTGGCAATGGTTTAATTTATTGGTCATGCCAAAAGAATAAATCACCAATAAGTTTGCTCAATTATTTTTAATTCATTTGGTTTAAAGGAAAAAGAAAGCTCACTTAAATGGCTAAGATAATAATATAAACGAAGGTTTGGCATAAAACAAATATAATACACGCCAAGAGTGCATACCACACAAAATTATCGAATATTTTCCATTTTAATTTTTTTTAGAAATACCTGAGACCGTCAATATGCGATGTTCTTCTCTCTACTAGCCCACGAAAACGACAACGTCCCCAATTTATAAACACTACACCCTTACACATTGAACGCTCTATATATAGTGTGCATCATTGGCCTCGTAGTTAAACTCATATCGATCGAAGAAAACTCAACTAGAGCTAACAAGAGTTTGAAGACCTCCTCTTGTTACCTTGTTCATTTTCTTCGATAATCTTATTCAGAAGTAGAAAGCTCATTTACAAAACAATCAAA | 737 bp |
| *BoIAA7a* | Bo1g103470 | CATTGAATACACGTTTTCACCTAAATAAAACGGTAACATGTTTCGTTGTAATGTCGATGTAATGATTACGACGTATTTCTCATTCCACGTAAATTCGTCGTAAACTTACATGGATTTTACGACGAAAACAATTCGTCGTAAATATTCGTTGTTATAGGCACGTTTTCTTGTAGTGTAATATATAGTCCAACAAACAAATGAATACTCTTATTTCTTACACAATATATATACTACATTCGTATTTTCATATAAAGTTAGATTCAGATTTATAAATGAAAATATTTCGGGCTGATGTGCACGTAAAATTCTAGTTAGTATTACGAAATGTTGATTCTAAGGTTAGGGAGACTATGGACCACTCTTTTTTTTTNGATCAACTGAGAGACTATGGAGCACAACTTCCATTTTTTGGTAATTAAGAAACATCTGTAGGATAAAAGACTCCAATAGTTTTAAACTGCTGGTTATTGTATAAACCTGTTGCAAATTTTAATTCCACCACTTTTTCTTTTCTTACTAAAAGATTATCTCTTGCCAACACCAATCAAAATTTGTTTACCTGAAAAGATTATTTTCCATTATTAAACAAAGAATCCCAGCTGTTGTAGTAAAAGTAATCTATTAACATCTAATCAGTCTCACGTGTCAAAAACCCCACACTAACACATGAAGAAAAAAGAGTAAAGGACATGCACATCCAATTCATACACCCCACCACCATCCTCTCTCCTTATAATTATCAACCCCCCTTCCTCTCCCTACTTCTTCCCCTCTTACCAAATCAACATCTCTAATCTACTACAAC | 805 bp |
| *BoIAA7b* | Bo3g081600 | CGACTCCAAATAATTATCCTTCTAACTGCACTGAAACCTAAGGCTGAAAAAAAAATCCAAACCCGAAAAACTGAACCTAATCCAATCCGAAAAAGTAATATCGAATCTAAACTAAAACTGGTTAAATATTTGAACTGGTTTAAAATTTTGGTATCGAAAGAACTGGAACTGAACCCAACCCAACCCAAACCGTAACATTTCGGGTATCCGAATATATCTGAATCAGATTTATATACTTAAATATACTAACTATTTTTAGATTTAATATCTAAAAAAGTTATTAAAAATATATAAGATATTTGAACTTGTCCAAAAAATATAAAATATTTTTAAGTTGTTCAAAATACTTAAAAATATATACAAATAGTCAAAATTAAATATTTAAAACAATTCAAATGAGTTATTTGAACCCAAACCGTACCGAACCCGCAAAGATTCTAACAGAACCCACGAAGATCCGAACTAAACCGAATCAAAATTTGTAAATACCTGAATATAACTAAAATCTTTTAACTCCGAAAACGCGAAACGAATAGACTCGAAATAACCCAAACGGTTACCCGAATACCTCGCTGAAACCGATTAGAACTTCTAATTCTACCACTTTTTCTTTTATAGCTAAAACAAAAAAAAATTATCTCTTGTCAGCGCCATCAAAATTTCTAACAGAGAATCCCAGCTTTTGTAGCAAAAGAAATATATTAACATCTAATCAATCCCACGTGTCAAAAACCCCACATATGGAGAGAAAAAGAGCAAATGACATGCACATGCATTTCATACACCCCACCACCGTTTTCTCTCCTTATATTGTTTCTGCCCTTTCCTCTCTCTTCTTCTTCCCCTCATACCAAATCAGAGAGAGAAATAACATCTTTAATCTATTAGAAGTAACGACCCAGCTTTCTTGTACAAAGTTG | 920 bp |
| *BoIAA7c* | Bo5g093110 | AGGAATGGTTGTTCGTTTTCAAATTTAACAAGGAATAATTTTGTTCTTTATTTCTCTACAGAAAAAAAAGAATGAGAAATAAAACATTATTCCTTGTGAATGGTGATTTTTTGTAGGAACGTCAGAGAATGCATTATTCCCCATCATTTCTTAGTCACCATTCATTATGACTTTTTAAGATAGTGCAAGAGTAACCAATGTCTAAAATAGTCAAAAATTGGTTTTATGGCAAAAAAGACAATGGGTAGATTAGTTGTTCATTGCTAGGAAATATCCCTTTTAAATTCAGTATATTAGAGTATGAGGAAACTGGGACTATAAAAATTCAAAATTTTTTGGTTATCAGAACAAAAGAAATTTACTAAAATAATCCAAATAGTTTTAAGTTGATACTAATTTTTAATTCCGTCCCTTTACTTTATAGCTAGACAAAATCATTCATTGTCAATCACCAATCAAAAGTTTTAACTAGAAAATTATTGATCATCATGAAACAAAGAACCAAAGCTAGTAAAAATGATTTATTAACATCTAATAATCATAAGTTACAAAGAAAAAAAAACATCTAATAATCCCAAGTGGCAAAAACCCCACACATGAAGAAAAAAAGTGCAAAGGACATGCACATGCAGTTCATACACCCCACCACCATCCTCTCTCCTTATATTATTATTGCCCCCTTCCTCTCCCTTCTTCCCCTCTTACCAAATCAGAGAGAGAGAGAGAAATA | 730 bp |
| *BoERF043* | Bo3g171090 | TAAATAAAAAGATAACAAACTTTTAACAAAAAAAAATCGTAGTTACATCCACCAACCTTTCTCGTAGGTAAGCTACAACTTTATGTGTCCCTGACTCTCTTTCGACCATGTATATAAGTTGATAATACTATGAGGAACCATTGAGAGATTTGGAAACATCTGAGAAGACGAGACTAGAAATTTCATCAAACAATGAAACAAACAAGAACTATCTTTTCAAACTCTTACTACACCCAAACAATGTCGCACTACTTTTGACTTAGAAACTTCTCATTATAATATTCTTTCTTTCTACAACAACACAAAT | 307 bp |
| *BoATHB-X* | Bo6g112570 | TGCGAGACACATGAATAAACTCAGACGTCCAATCAGTGATACTTATGTAATATTTTTTTGTTAAATAATCAACTATATTCAACTATATATGTTTTATTTTTGCTTAAACTATTTTCAACTATGTTACACTATTTACCTTAATATAATTATATAAACTATAGTTTTCGGAACTTTCAAGAAAAATGTATTTATGTTTTAAATGGTACATGATTAGGTTTCAGATATTAAACATCATAGTACTACTTTTTATAATTAATACGGTATTAGTTAATTATACGTTCTACGACATTTATAACAAGATCTTTTAAGCTTAGACCATTCTCGTTTTTTTTCTAAGATTATTGTATTCGTAACGCTCAACGCAAAACAACACTTCGGCCACCCGTTACTGTCCCACCCAAGGCCATAATGAATTCAACAACTCGGAAATTAAAACTGGTAAAACACGTAATAATTGATCATACGAATATGATTTGACTTAGCAACCCTAGTCAAACCCATTTCTTCCGCCAATTACACGTAAATACACCTCTATTTAATTCAATTCCTTTAAAAATCTCATCTTACACTGCCATTTGTTTTTTCAGTAATTTTTGGTTCATTGACCAGCTTCGGACTTTTATCCTTTGTCGTGTTTTGGGACTTTCTTTGAAATTTGATAATTCTGTGAGGCTTAATTCCTTATTCCATATTCCACGCTTTATAACAAAAATTTTCTCCTTTGTCTGGCTTTTTCTCTTTTGTTCTTTCACATGTGTATATACACACAGACATACACACACTTACATTTCTAATATCATATAATACTTCAACTTTACCTTTTCTCTTCCAAAAAAAAACTATTTCTCTTCATCAGATTGTATTTGATCATTCCAGCTTTTCCTCATCCTCCTCT | 895 bp |
| *BoTIC40* | Bo2g013210 | TAAAATAAATAAAAAAATATCTATCGTAATATTAGATAAGTAAATATACAATATAAGAAAAATAAATAATAATTTTATATACAATATAAGAAATAGAAAAACTACATTAAATAACTATCTGTAAATGTATAGTAAAATAAATAATAAGTAAATATACAATATAAAAATTAGAAATATTACTTTCAACATCTATCGTAATATTATCATTTGATATAAAATCAAGAAAAATAAAATAAATTAATATAAAATATAAGAAAAACTAAATAATAAGTAAATATATAATATAAGAAATGAAAAACTACTTTAAACATTTATGTGAAAAATACAAAGTTTTTTCTGGTTTTATCTAAGGATATATGTATAAATATACAATATGTATAATTAAATACAAACATACAGTTCATAATATGACGTCCAAAATAATAGTTATATAGTTAACATTTTGAAATCAAAATAGCAAGCGCGGTTTTACCCTTACTATATTGGGCTTAAAGGTGAAGGGATGACTTTTGATTTATCAAGAAGGAGTTTTTGCACCCTTGGATTACAGCTCAGAGTAAACGATCTCGACGGTCAGATACTCGCCACCTGTAAATCTCCAAGCTTTTTGGTGTACTTGGCTGCACTGGATCCGAGACCGCTACTACACCCAAACCTGATGCTTGAAGGATACAGTAGCTTTAGACCAAAG | 691 bp |
| *BoCYP450* | Bo3g023170 | CAACTGTCCAAATAATAAATTACCGTAAAAACGACGTAACTGCATTTCATATTTTCCATTGGCATTAATATCAAACGGTTAAGATTAACTAAAAAGAAACGGTTATGCAAATTGGCGTTACATAATGTGTCCACTAAGCAAATTACTTAATTAGAAGCTCCTTAAATAAAAGAAAATAAATTGTTGGACATACCCTTTATCAATAGGTCATGTTGCTGTTTTAATAGTATAGATAATTAATAATTAATTCATCCTTTTTCCGTTGGTCAACTGCGTAATTTATCAACTAGATTAAGGGAAAAACAAATATTATTTTGGTGACAAAAAAAATACAGGTAAGTTGTGTTTTAGCTTTTTACCTTTTAAGGATTATAAAATACAACATCCATACATTTATATCACTTTCAGTTATTCATTCAAAAAAGCAGCCGGAGAAACTCCACAAGACTATAGCCAAAAAGGAATAAGAGAGGCATGCTAAATGCCGATAATGTTTGACCAGAAAATGGTCAAGAAACGTTATGAAGCAACATAAAATAAAATTCAACAAATATAATAAATTAAAAGAGAGATGAGACTTTCGTTGGTCCAAGTCTGACATTCTTCTACACCCGAACCTAGAAAACATCTATAAATACCTCATTTGGTTGATTCCCATGTTTCATCTGCATACATTCTGAATACAACATCCAGTATTCCACCAAAGAAACACTTACGCATATATCTATTC | 730 bp |

**Table S5:** Strain and plasmid used in the present study

| **Strain** | **Description** | **Resistance** | **Reference** |
| --- | --- | --- | --- |
| *Xanthomonas campestris* pv. *campestris* | | | |
| 8004 | Wild-type british strain harvested from *Brassica olearacea var. botrytis* in 1912, Rifampicin-resistant derivative of *Xanthomonas campestris* pv. *campestris* 8000 (NCPBB1145), RifR | Rif(50 μg/mL) | Turner and al., 1984 |
| Δ26 | 8004 strain deleted for the *avrBs1, hrpW, xopAC, xopH, xopX1, xopX2, xopF, xopD, xopAN, xopQ, xopK, xopJ, xopG, avrXccA1, avrXccA2, xopAM, xopE2, xopAH, xopR, xopL, xopZ, xopAG, xopP, xopAL1, xopAL2* and *xopN* genes |  | European patent: EP20305940.7 |
| Δ26T3E::*tal12a* | Genomic integration of *tal12a* gene into the background of Δ26 |  | Constructed during this study |
| Δ26T3E::*tal14c* | Genomic integration of *tal14c* gene into the background of Δ26 |  |  |
| CN08 | Wild-type chinease strain harvested from *Raphanus sativus var. longipinnatus* in 2002, Rifampicin-resistant derivative |  | He and al., 2007 |
| CN08Δ*tal12a* | CN08 strain spontaneous mutant for *tal12a* gene |  | Constructed during this study |
| CN08Δ*tal12a* + EV | CN08Δ*tal12a* + empty vector (pSKX1) |  |  |
| CN08Δ*tal12a* + Tal12a | CN08Δ*tal12a* + vector containing Tal12a (pSKX1) |  |  |
| CN08 CRISPRi-RBS | CN08 with CRISPRi construction integrated genomically with single guide RNAs (sgRNAs) silencing *Xcc* CN08 TALEs | Rif(50 μg/mL), Gm(10µg/mL) | Zárate‐Chaves and al., 2023 |
| CN08 CRISPRi-ctrl | CN08 with CRISPRi construction integrated genomically where sgRNA scaffold was replaced by promoterless *uidA* gene |  |  |
| Xca5 | Wild-type american strain coming from M. Daniels, Rifampicin-resistant derivative | Rif(50 μg/mL) | Kay and al., 2005 |
| Xca5 CRISPRi-RBS | Xca5 with CRISPRi construction integrated genomically with single guide RNAs (sgRNAs) silencing *Xcc* CN08 TALEs | Rif(50 μg/mL), Gm(10µg/mL) | Audran, unpublished data |
| Xca5 CRISPRi-ctrl | Xca5 with CRISPRi construction integrated genomically where sgRNA scaffold was replaced by promoterless *uidA* gene |  |  |
| *Agrobacterium tumefaciens* | | | |
| GV3101 containing the plasmid pJawohl11-GW-*gus* below |  | **Strain:** Rif(50µg/mL), Gm(15µg/mL) **Plasmid:** Km(50µg/mL) | **Plasmid:** Serrano and al., 2007 |
| pJawohl11-*mCherry*-*gus* | Integration of *mCherry* gene in pJAWOL11-GW-*gus* plasmid | Rif (50µg/mL), Km(50µg/mL), Gm(15µg/mL) | Constructed during this study |
| pJawohl11-p*BoSWEET13*-*gus* | Integration of *BoSWEET13* promoter in pJAWOL11-GW-*gus* plasmid |  |  |
| pJawohl11-p*BoSWEET14c*-*gus* | Integration of *BoSWEET14b* promoter in pJAWOL11-GW-*gus* plasmid |  |  |
| pJawohl11-p*BoCYP450*-*gus* | Integration of *BoCYP450* promoter in pJAWOL11-GW-*gus* plasmid |  |  |
| pJawohl11-p*BoIAA7a*-*gus* | Integration of *BoIAA7a* promoter in pJAWOL11-GW-*gus* plasmid |  |  |
| pJawohl11-p*BoIAA7b*-*gus* | Integration of *BoIAA7b* promoter in pJAWOL11-GW-*gus* plasmid |  |  |
| pJawohl11-p*BoIAA7c*-*gus* | Integration of *BoIAA7c* promoter in pJAWOL11-GW-*gus* plasmid |  |  |
| pJawohl11-p*BoERF043*-*gus* | Integration of *BoERF043* promoter in pJAWOL11-GW-*gus* plasmid |  |  |
| pJawohl11-p*BoATHB-X*-*gus* | Integration of *BoATHB-X* promoter in pJAWOL11-GW-*gus* plasmid |  |  |
| pJawohl11-p*BoTIC40*-*gus* | Integration of *BoTIC40* promoter in pJAWOL11-GW-*gus* plasmid |  |  |

Rif: rifampicin; Gm: gentamicin; Km: kanamicin; Cm: chloramphenicol

**Table S6**: Primer used in the present study

| **Description** | **Forward (5'-3')** | **Reverse (5'-3')** |
| --- | --- | --- |
| Gene expression | | |
| *BoActin2* | CCGGAGGTCTTGTTCCAGCCATC | GTTCCACCACTGAGCACAATGTTAC |
| *BoSWEET13* | GTTTTGCCGCCATTGTCCTT | GGAGCTGCGAAAACGATGAC |
| *BoSWEET14c* | AGAGCTAACAAGAGTTTGAAGACC | TGATGTTGCCCATGATTCCAAATG |
| *BoCYP450* | GAGGTGAACGGCAGAGTGAT | TCCGGCCTAAACTCGTTGAC |
| *BoIAA7a* | AGATGGCGATTGGATGCCAG | GAGCAAGTCCAATTGCTTCAGA |
| *BoIAA7b* | CAGCTTATGAACCTTAACGCGA | GGCTCTCGACGGCTTTAGTC |
| *BoIAA7c* | GAATCTGGCAAAATCGGCGG | AAGCTGAGGCGACGTTGTTA |
| *BoERF043* | CGGCGTTTTGTCACGAAAGT | TCCTCGTCAATGCTCGTCAC |
| *BoATHB-X* | AAGAACTGCGAGCTCTTAAGCC | CTCCACGGCATTAGAGCCATT |
| *BoTIC40* | TCCTGGATTCCCGACAGGAT | TTATAGGCGTAGGTTGCGGC |
| Promoter cloning | | |
| p*BoSWEET13* | GGGGACAAGTTTGTACAAAAAAGCAGGCTTCCAGCACAGTTCGCTTTTAGTGTGGG | GGGGACCACTTTGTACAAGAAAGCTGGGTCTTGTTTCTGCTTTCTTGATTATTTC |
| p*BoSWEET14c* | GGGGACAAGTTTGTACAAAAAAGCAGGCTTCTCTACCATGCAAACTTTGGAATAG | GGGGACCACTTTGTACAAGAAAGCTGGGTCTTTGATTGTTTTGTAAATGAGCTTTC |
| p*BoCYP450* | GGGGACAAGTTTGTACAAAAAAGCAGGCTTCCATGTATGATTTAATTTTTCGTGGTC | GGGGACCACTTTGTACAAGAAAGCTGGGTCCTCGTATATCTCTTCAAATATGTAG |
| p*BoIAA7a* | GGGGACAAGTTTGTACAAAAAAGCAGGCTTCCATTGAATACACGTTTTCACC | GGGGACCACTTTGTACAAGAAAGCTGGGTCGTTGTAGTAGATTAGAGATGTT |
| p*BoIAA7b* | GGGGACAAGTTTGTACAAAAAAGCAGGCTTCGACTCCAAATAATTATCCTTCT | GGGGACCACTTTGTACAAGAAAGCTGGGTCGTTACTTCTAATAGATTAAAGATG |
| p*BoIAA7c* | GGGGACAAGTTTGTACAAAAAAGCAGGCTTCAGAAGAATAAAATAGCAGGAATGAAA | GGGGACCACTTTGTACAAGAAAGCTGGGTCGTTACCTGTAATAGATTATAGATGT |
| p*BoERF043* | GGGGACAAGTTTGTACAAAAAAGCAGGCTTCCAATATCTCGTGTTCCGCGTTG | GGGGACCACTTTGTACAAGAAAGCTGGGTCTGTGATTCTTGAAATATGATTTTAG |
| p*BoATHB-X* | GGGGACAAGTTTGTACAAAAAAGCAGGCTTCTGCGAGACACATGAATAAAC | GGGGACCACTTTGTACAAGAAAGCTGGGTCAGAGGAGGATGAGGAAAAGC |
| p*BoTIC40* | GGGGACAAGTTTGTACAAAAAAGCAGGCTTCATTAAACATCTATGTAAAAAATGTATAG | GGGGACCACTTTGTACAAGAAAGCTGGGTCGTTTATATGATGAGGAAGAAGAAGA |

**Table S7**: RVD sequences of the arTALEs used in the present study

| Name | Targeted gene | RVD sequence |
| --- | --- | --- |
| arTALE7.1 | *BoIAA7c* | HD-NG-NG-HD-HD-HD-HD-NG-HD-NG-NG-NI-HD-HD-NI-NI-NI-NG |
| arTALE7.2 | *BoIAA7c* | NH-HD-HD-HD-HD-HD-NG-NG-HD-HD-NG-HD-NG-HD-HD-HD-NG-NG |
| arTALE13.1 | *BoSWEET13* | NI-NG-HD-NG-NN-NI-NH-NI-HD-HD-NI-NI-NG-NN-NI-NG-NN-NG |
| arTALE13.2 | *BoSWEET13* | NI-NG-NG-NI-NI-HD-NI-HD-NG-NI-HD-NI-HD-HD-HD-HD-NG-NG |
| arTALE14.1 | *BoSWEET14c* | NI-NG-NI-NG-NI-NG-NI-NN-NG-NN-NG-NH-HD-NI-NG-HD-NI-NG |
| arTALE14.2 | *BoSWEET14c* | NI-HD-NI-HD-NI-NG-NG-NN-NI-NI-HD-NH-HD-NG-HD-NG-NI-NG |

**Table S8**: Top 300 genes upregulated by Tal12a and Tal14c in cauliflower identified by RNA-seq (FDR<0.01), ranked by log₂ fold change for each TALE.

| **TALE** | **Rank** | **Gene name** | **Log_2_FC** | **FDR** | **Gene annotation** | **Putative product** |
| --- | --- | --- | --- | --- | --- | --- |
| Tal12a | 1 | Bo9g097170 | 8,95E+00 | 3,00E-127 | Nodulin MtN3 family protein [Source:TAIR;Acc:AT5G50800](projected from arabidopsis_thaliana,AT5G50800) | sugar transporter SWEET |
| Tal12a | 2 | Bo3g023170 | 8,89E+00 | 1,48E-99 | cytochrome P450, family 715, subfamily A, polypeptide 1 [Source:TAIR;Acc:AT5G52400](projected from arabidopsis_thaliana,AT5G52400) | cytochrome P450 |
| Tal12a | 3 | Bo03924s010 | 8,05E+00 | 1,53E-70 |  | [histone H3]-lysine(4) N-trimethyltransferase chromatin remodeling SET family |
| Tal12a | 4 | Bo2g159060 | 7,89E+00 | 5,46E-76 |  | protein ENHANCED DISEASE RESISTANCE 2 |
| Tal12a | 5 | Bo6g074490 | 7,79E+00 | 2,41E-121 | plasmodesmata callose-binding protein 5 [Source:TAIR;Acc:AT3G58100](projected from arabidopsis_thaliana,AT3G58100) | glucan endo-1,3-beta-D-glucosidase |
| Tal12a | 6 | Bo5g021050 | 7,41E+00 | 2,76E-69 | 2-oxoglutarate (2OG) and Fe(II)-dependent oxygenase superfamily protein [Source:TAIR;Acc:AT1G15540](projected from arabidopsis_thaliana,AT1G15540) | (-)-deoxypodophyllotoxin synthase |
| Tal12a | 7 | Bo9g029530 | 7,25E+00 | 2,91E-50 |  | transcription factor MYB-HB-like family |
| Tal12a | 8 | Bo3g021660 | 7,16E+00 | 2,42E-74 |  | glucan endo-1,3-beta-D-glucosidase |
| Tal12a | 9 | Bo1g104990 | 7,09E+00 | 6,67E-80 |  | bifunctional inhibitor/plant lipid transfer protein/seed storage helical |
| Tal12a | 10 | Bo9g048800 | 7,09E+00 | 1,67E-50 |  | putative protein |
| Tal12a | 11 | Bo20985s010 | 6,86E+00 | 5,35E-06 |  | proteinase inhibitor I3, Kunitz legume, kunitz inhibitor STI-like superfamily |
| Tal12a | 12 | Bo3g171090 | 6,55E+00 | 1,52E-27 |  | transcription factor AP2-EREBP family |
| Tal12a | 13 | Bo4g025620 | 6,50E+00 | 5,45E-41 |  | protein EARLY FLOWERING 4 |
| Tal12a | 14 | Bo5g098520 | 6,49E+00 | 2,53E-40 |  | Gnk2-like domain-containing protein |
| Tal12a | 15 | Bo6g112570 | 6,37E+00 | 3,89E-113 |  | transcription factor Homobox-WOX family |
| Tal12a | 16 | Bo2g024540 | 6,34E+00 | 3,43E-31 |  | transcription factor C2H2 family |
| Tal12a | 17 | Bo8g089750 | 6,20E+00 | 2,42E-91 |  | glucan endo-1,3-beta-D-glucosidase |
| Tal12a | 18 | Bo4g116730 | 6,19E+00 | 9,13E-65 |  | glucan endo-1,3-beta-D-glucosidase |
| Tal12a | 19 | Bo6g125680 | 6,14E+00 | 1,39E-28 |  | Galactose-binding-like domain superfamily |
| Tal12a | 20 | Bo4g070540 | 6,09E+00 | 9,83E-33 |  | (Z)-gamma-bisabolene synthase |
| Tal12a | 21 | Bo7g006000 | 6,08E+00 | 3,33E-91 | zinc ion binding [Source:TAIR;Acc:AT2G19385](projected from arabidopsis_thaliana,AT2G19385) | transcription factor C2H2 family |
| Tal12a | 22 | Bo6g121610 | 5,93E+00 | 2,27E-26 | AGAMOUS-like 67 [Source:TAIR;Acc:AT1G77950](projected from arabidopsis_thaliana,AT1G77950) | transcription factor MADS-type1 family |
| Tal12a | 23 | Bo3g022720 | 5,91E+00 | 6,96E-26 | Leucine-rich repeat protein kinase family protein [Source:TAIR;Acc:AT5G53320](projected from arabidopsis_thaliana,AT5G53320) | protein kinase RLK-Pelle-LRR-III family |
| Tal12a | 24 | Bo2g032390 | 5,77E+00 | 3,98E-24 |  | putative protein |
| Tal12a | 25 | Bo3g009500 | 5,70E+00 | 6,37E-23 |  | transcription factor C2H2 family |
| Tal12a | 26 | Bo2g046150 | 5,56E+00 | 4,57E-61 | RING/U-box superfamily protein [Source:Projected from Arabidopsis thaliana (AT5G53110) TAIR;Acc:AT5G53110] | RING-type E3 ubiquitin transferase transcription factor C2H2 family |
| Tal12a | 27 | Bo2g092980 | 5,55E+00 | 7,33E-25 | Nitrate reductase [Source:UniProtKB/TrEMBL;Acc:A0A0D3ARC1] | nitrate reductase (NADH) |
| Tal12a | 28 | Bo4g106400 | 5,54E+00 | 6,47E-22 |  | S-phase kinase-associated protein |
| Tal12a | 29 | Bo3g002110 | 5,48E+00 | 2,77E-78 |  | F-box protein, plant |
| Tal12a | 30 | Bo01217s020 | 5,43E+00 | 1,49E-14 |  | DnaJ domain, Chaperone J-domain superfamily |
| Tal12a | 31 | Bo6g051130 | 5,37E+00 | 3,07E-20 | Family of unknown function (DUF716) [Source:TAIR;Acc:AT1G55230](projected from arabidopsis_thaliana,AT1G55230) | putative protein |
| Tal12a | 32 | Bo6g015670 | 5,34E+00 | 1,01E-60 |  | GH3 family protein |
| Tal12a | 33 | Bo4g166770 | 5,29E+00 | 4,70E-27 |  | putative protein |
| Tal12a | 34 | Bo4g144150 | 5,24E+00 | 1,20E-41 | brassinosteroid-6-oxidase 1 [Source:TAIR;Acc:AT5G38970](projected from arabidopsis_thaliana,AT5G38970) | cytochrome P450 |
| Tal12a | 35 | Bo7g014600 | 5,21E+00 | 1,09E-73 | 3-hydroxy-3-methylglutaryl-CoA reductase 2 [Source:TAIR;Acc:AT2G17370](projected from arabidopsis_thaliana,AT2G17370) | hydroxymethylglutaryl-CoA reductase (NADPH) |
| Tal12a | 36 | Bo6g010610 | 5,14E+00 | 2,83E-24 | hexokinase 3 [Source:TAIR;Acc:AT1G47840](projected from arabidopsis_thaliana,AT1G47840) | phosphotransferase with an alcohol group as acceptor |
| Tal12a | 37 | Bo4g186390 | 5,13E+00 | 2,42E-33 | UDP-glucosyl transferase 73C2 [Source:TAIR;Acc:AT2G36760](projected from arabidopsis_thaliana,AT2G36760) | hexosyltransferase |
| Tal12a | 38 | Bo5g126380 | 5,06E+00 | 9,71E-15 |  | asparaginase |
| Tal12a | 39 | Bo13741s010 | 5,01E+00 | 5,64E-26 |  | GH3 family protein |
| Tal12a | 40 | Bo9g073020 | 5,00E+00 | 1,50E-23 |  | SPX domain-containing protein |
| Tal12a | 41 | Bo3g058840 | 5,00E+00 | 6,60E-51 |  | heavy metal-associated domain, HMA, heavy metal-associated domain superfamily |
| Tal12a | 42 | Bo5g062720 | 4,92E+00 | 9,95E-92 |  | RNA recognition motif domain, nucleotide-binding alpha-beta plait domain superfamily |
| Tal12a | 43 | Bo2g042250 | 4,91E+00 | 3,66E-15 | homeobox protein 52 [Source:TAIR;Acc:AT5G53980](projected from arabidopsis_thaliana,AT5G53980) | transcription factor Homobox-WOX family |
| Tal12a | 44 | Bo7g104770 | 4,87E+00 | 1,32E-47 |  | transcription factor AP2-EREBP family |
| Tal12a | 45 | Bo1g154800 | 4,86E+00 | 6,82E-16 | UTP--glucose-1-phosphate uridylyltransferase 2 [Source:Projected from Arabidopsis thaliana (AT3G03250) UniProtKB/Swiss-Prot;Acc:Q9M9P3] | UTP--glucose-1-phosphate uridylyltransferase |
| Tal12a | 46 | Bo2g068130 | 4,84E+00 | 3,99E-20 |  | putative protein |
| Tal12a | 47 | Bo1g022580 | 4,82E+00 | 6,71E-80 | Tubulin beta chain [Source:UniProtKB/TrEMBL;Acc:A0A0D3A4N8] | tubulin |
| Tal12a | 48 | Bo1g103470 | 4,80E+00 | 8,84E-44 | Auxin-responsive protein [Source:UniProtKB/TrEMBL;Acc:A0A0D3ABD5] | transcription factor interactor and regulator AUX-IAA family |
| Tal12a | 49 | Bo5g098580 | 4,79E+00 | 3,15E-23 |  | Gnk2-like domain-containing protein |
| Tal12a | 50 | Bo5g093110 | 4,69E+00 | 8,06E-68 | Auxin-responsive protein [Source:UniProtKB/TrEMBL;Acc:A0A0D3CGZ6] | transcription factor interactor and regulator AUX-IAA family |
| Tal12a | 51 | Bo1g150090 | 4,68E+00 | 2,89E-16 |  | putative protein |
| Tal12a | 52 | Bo9g117230 | 4,65E+00 | 1,88E-79 |  | putative protein |
| Tal12a | 53 | Bo4g004680 | 4,57E+00 | 2,94E-22 |  | transcription factor WRKY family |
| Tal12a | 54 | Bo5g098440 | 4,54E+00 | 1,26E-29 |  | putative protein |
| Tal12a | 55 | Bo1g010370 | 4,53E+00 | 3,70E-18 | K+ transporter 5 [Source:TAIR;Acc:AT4G32500](projected from arabidopsis_thaliana,AT4G32500) | cyclic nucleotide-binding domain, potassium channel, voltage-dependent, EAG/ELK/ERG |
| Tal12a | 56 | Bo8g108290 | 4,53E+00 | 2,87E-13 | Hop1 [Source:Projected from Arabidopsis thaliana (AT1G12270) UniProtKB/TrEMBL;Acc:A0A178WM19] | Heat shock chaperonin-binding, tetratricopeptide-like helical domain superfamily |
| Tal12a | 57 | Bo4g139450 | 4,47E+00 | 7,68E-68 | DNA-directed RNA polymerases II and V subunit 6B [Source:Projected from Arabidopsis thaliana (AT2G04630) UniProtKB/Swiss-Prot;Acc:Q9SJ96] | DNA-directed RNA polymerase |
| Tal12a | 58 | Bo4g197330 | 4,46E+00 | 1,55E-12 |  | transcription factor NAM family |
| Tal12a | 59 | Bo2g089880 | 4,46E+00 | 8,36E-15 |  | RING-type E3 ubiquitin transferase transcription factor C2H2 family |
| Tal12a | 60 | Bo00851s030 | 4,44E+00 | 6,76E-10 |  | RNA helicase |
| Tal12a | 61 | Bo4g151230 | 4,44E+00 | 1,43E-39 |  | sinapoylglucose--sinapoylglucose O-sinapoyltransferase |
| Tal12a | 62 | Bo5g042850 | 4,39E+00 | 2,80E-05 | Flavin-dependent oxidoreductase FOX2 [Source:Projected from Arabidopsis thaliana (AT1G26390) UniProtKB/Swiss-Prot;Acc:Q9FZC5] | oxidoreductase |
| Tal12a | 63 | Bo7g063720 | 4,36E+00 | 6,95E-12 |  | putative protein |
| Tal12a | 64 | Bo5g093480 | 4,36E+00 | 1,10E-08 |  | leucine-rich repeat-containing, plant-type, leucine-rich repeat domain superfamily |
| Tal12a | 65 | Bo7g041140 | 4,33E+00 | 8,82E-12 |  | reverse transcriptase zinc-binding domain-containing protein |
| Tal12a | 66 | Bo5g148490 | 4,33E+00 | 4,75E-37 |  | MT-associated protein TORTIFOLIA1/SPIRAL2 |
| Tal12a | 67 | Bo4g088250 | 4,31E+00 | 6,67E-19 | CRIB domain-containing protein RIC9 [Source:Projected from Arabidopsis thaliana (AT1G61795) UniProtKB/Swiss-Prot;Acc:Q1G3Y0] | CRIB domain-containing protein RIC1 |
| Tal12a | 68 | Bo4g115410 | 4,29E+00 | 1,37E-06 |  | flavonol synthase |
| Tal12a | 69 | Bo3g184820 | 4,22E+00 | 3,43E-22 | Dirigent protein [Source:UniProtKB/TrEMBL;Acc:A0A0D3BN47] | dirigent protein |
| Tal12a | 70 | Bo9g084470 | 4,22E+00 | 1,18E-17 | arabinogalactan protein 5 [Source:TAIR;Acc:AT1G35230](projected from arabidopsis_thaliana,AT1G35230) | classical arabinogalactan protein |
| Tal12a | 71 | Bo5g020810 | 4,16E+00 | 4,13E-05 |  | glyoxalase/Bleomycin resistance protein/Dihydroxybiphenyl dioxygenase |
| Tal12a | 72 | Bo1g003710 | 4,16E+00 | 2,81E-11 |  | cytochrome P450 |
| Tal12a | 73 | Bo3g162350 | 4,16E+00 | 2,03E-31 | Tubulin beta chain [Source:UniProtKB/TrEMBL;Acc:A0A0D3BKN7] | tubulin |
| Tal12a | 74 | Bo8g055870 | 4,16E+00 | 3,26E-11 |  | Phytocyanin domain, cupredoxin |
| Tal12a | 75 | Bo8g108840 | 4,07E+00 | 1,13E-21 | Nucleotide/sugar transporter family protein [Source:TAIR;Acc:AT4G03950](projected from arabidopsis_thaliana,AT4G03950) | triose phosphate/phosphoenolpyruvate translocator, sugar phosphate transporter |
| Tal12a | 76 | Bo2g013220 | 4,07E+00 | 3,30E-64 |  | DNA repair protein Rad4 |
| Tal12a | 77 | Bo2g013210 | 4,07E+00 | 2,16E-64 |  | Heat shock chaperonin-binding, STI1/HOP, DP domain-containing protein |
| Tal12a | 78 | Bo7g114260 | 4,04E+00 | 1,07E-58 |  | translocase |
| Tal12a | 79 | Bo9g027350 | 4,04E+00 | 3,71E-57 |  | calmodulin binding protein, central |
| Tal12a | 80 | Bo5g114060 | 4,02E+00 | 1,46E-19 |  | complex 1 LYR protein |
| Tal12a | 81 | Bo5g052440 | 4,01E+00 | 8,53E-40 |  | bifunctional inhibitor/plant lipid transfer protein/seed storage helical |
| Tal12a | 82 | Bo1g006270 | 4,01E+00 | 6,07E-34 |  | vacuolar protein sorting-associated protein Ist1 |
| Tal12a | 83 | Bo1g147150 | 3,97E+00 | 4,58E-71 |  | RNA recognition motif domain, nucleotide-binding alpha-beta plait domain superfamily |
| Tal12a | 84 | Bo4g046230 | 3,96E+00 | 1,23E-14 |  | transcription regulator GNAT family |
| Tal12a | 85 | Bo9g120190 | 3,93E+00 | 1,91E-20 |  | glucan endo-1,3-beta-D-glucosidase |
| Tal12a | 86 | Bo24577s010 | 3,93E+00 | 9,80E-04 |  | Heat shock protein 70kD domain superfamily |
| Tal12a | 87 | Bo3g066850 | 3,93E+00 | 1,32E-23 |  | transcription factor NAM family |
| Tal12a | 88 | Bo4g186440 | 3,93E+00 | 1,70E-09 | Xyloglucan endotransglucosylase/hydrolase [Source:UniProtKB/TrEMBL;Acc:A0A0D3C4M8] | xyloglucan-specific endo-beta-1,4-glucanase |
| Tal12a | 89 | Bo1g051580 | 3,91E+00 | 7,54E-52 | O-fucosyltransferase family protein [Source:TAIR;Acc:AT4G17430](projected from arabidopsis_thaliana,AT4G17430) | putative protein |
| Tal12a | 90 | Bo5g140100 | 3,89E+00 | 6,21E-37 |  | GDP-fucose protein O-fucosyltransferase |
| Tal12a | 91 | Bo6g120020 | 3,88E+00 | 6,62E-13 |  | thioredoxin-like protein |
| Tal12a | 92 | Bo7g056390 | 3,85E+00 | 2,18E-11 | unknown protein; FUNCTIONS IN: molecular_function unknown; LOCATED IN: chloroplast; BEST Arabidopsis thaliana protein match is: unknown protein (TAIR:AT1G70900.1); Has 59 Blast hits to 59 proteins in 15 species: Archae - 0; Bacteria - 0; Metazoa - 0 /.../i - 0; Plants - 59; Viruses - 0; Other Eukaryotes - 0 (source: NCBI BLink). [Source:TAIR;Acc:AT1G23110](projected from arabidopsis_thaliana,AT1G23110) | putative protein |
| Tal12a | 93 | Bo6g081510 | 3,85E+00 | 2,45E-46 | D-mannose binding lectin protein with Apple-like carbohydrate-binding domain [Source:TAIR;Acc:AT1G78820](projected from arabidopsis_thaliana,AT1G78820) | S-locus-specific glycoprotein/EP1 |
| Tal12a | 94 | Bo7g075770 | 3,85E+00 | 2,17E-10 | Transcription factor bHLH28 [Source:Projected from Arabidopsis thaliana (AT5G46830) UniProtKB/Swiss-Prot;Acc:Q9LUK7] | transcription factor MYC/MYB, transcription factor AIB/MYC |
| Tal12a | 95 | Bo01178s040 | 3,84E+00 | 5,73E-06 |  | tetrahydroberberine oxidase |
| Tal12a | 96 | Bo9g020410 | 3,84E+00 | 2,95E-08 |  | rapid ALkalinization Factor |
| Tal12a | 97 | Bo4g148310 | 3,79E+00 | 1,95E-05 |  | putative protein |
| Tal12a | 98 | Bo3g081600 | 3,78E+00 | 4,60E-18 | Auxin-responsive protein [Source:UniProtKB/TrEMBL;Acc:A0A0D3BCV8] | transcription factor interactor and regulator AUX-IAA family |
| Tal12a | 99 | Bo9g102490 | 3,76E+00 | 4,81E-39 |  | putative protein |
| Tal12a | 100 | Bo7g077060 | 3,71E+00 | 9,25E-08 |  | Heat shock protein HSP14.7/HSP23.5/HSP23.6 |
| Tal12a | 101 | Bo2g042150 | 3,68E+00 | 6,54E-39 |  | Band 7 domain, Stomatin/HflK family, STML2-like extension, Band 7/SPFH domain superfamily |
| Tal12a | 102 | Bo3g006950 | 3,65E+00 | 8,22E-09 |  | sugar phosphate transporter domain-containing protein |
| Tal12a | 103 | Bo2g051210 | 3,64E+00 | 7,03E-45 | unknown protein; FUNCTIONS IN: molecular_function unknown; INVOLVED IN: biological_process unknown; LOCATED IN: endoplasmic reticulum, plasma membrane; EXPRESSED IN: 24 plant structures; EXPRESSED DURING: 13 growth stages; Has 149 Blast hits to 149 /.../ns in 49 species: Archae - 0; Bacteria - 0; Metazoa - 98; Fungi - 0; Plants - 47; Viruses - 0; Other Eukaryotes - 4 (source: NCBI BLink). [Source:TAIR;Acc:AT1G65270](projected from arabidopsis_thaliana,AT1G65270) | putative protein |
| Tal12a | 104 | Bo7g071430 | 3,64E+00 | 2,12E-07 |  | (+)-abscisic acid 8'-hydroxylase |
| Tal12a | 105 | Bo3g083900 | 3,61E+00 | 4,08E-13 | cytochrome P450, family 82, subfamily G, polypeptide 1 [Source:TAIR;Acc:AT3G25180](projected from arabidopsis_thaliana,AT3G25180) | cytochrome P450 |
| Tal12a | 106 | Bo5g137450 | 3,61E+00 | 5,68E-11 |  | GAG-pre-integrase domain-containing protein |
| Tal12a | 107 | Bo4g125880 | 3,61E+00 | 4,40E-07 |  | nucleic acid-binding protein |
| Tal12a | 108 | Bo3g108110 | 3,60E+00 | 6,59E-43 |  | transcription factor HSF-type-DNA-binding family |
| Tal12a | 109 | Bo4g041250 | 3,55E+00 | 2,26E-08 |  | RING-type E3 ubiquitin transferase transcription factor C2H2 family |
| Tal12a | 110 | Bo3g109400 | 3,55E+00 | 3,19E-48 | Transducin/WD40 repeat-like superfamily protein [Source:TAIR;Acc:AT3G49660](projected from arabidopsis_thaliana,AT3G49660) | transcription factor WD40-like family |
| Tal12a | 111 | Bo2g024810 | 3,54E+00 | 1,05E-06 |  | fasciclin-like arabinogalactan protein, group A |
| Tal12a | 112 | Bo4g059980 | 3,53E+00 | 7,76E-40 |  | Acyl-lipid omega-3 desaturase (cytochrome b5), endoplasmic reticulum |
| Tal12a | 113 | Bo7g048390 | 3,52E+00 | 4,52E-14 |  | glutathione transferase transcription factor MYB family |
| Tal12a | 114 | Bo7g109890 | 3,52E+00 | 8,25E-07 | Bidirectional sugar transporter SWEET [Source:UniProtKB/TrEMBL;Acc:A0A0D3DFZ0] | sugar transporter SWEET |
| Tal12a | 115 | Bo2g002610 | 3,50E+00 | 3,90E-06 |  | very-long-chain 3-oxoacyl-CoA reductase |
| Tal12a | 116 | Bo7g117780 | 3,49E+00 | 1,66E-06 |  | small auxin-up RNA |
| Tal12a | 117 | Bo02914s010 | 3,46E+00 | 1,07E-05 |  | transmembrane protein |
| Tal12a | 118 | Bo7g114890 | 3,46E+00 | 3,69E-48 | ARM repeat superfamily protein [Source:TAIR;Acc:AT4G30990](projected from arabidopsis_thaliana,AT4G30990) | U3 small nucleolar RNA-associated protein |
| Tal12a | 119 | Bo2g126880 | 3,45E+00 | 4,83E-14 |  | disease resistance protein |
| Tal12a | 120 | Bo7g110790 | 3,43E+00 | 3,21E-08 | myo-inositol oxygenase 4 [Source:TAIR;Acc:AT4G26260](projected from arabidopsis_thaliana,AT4G26260) | inositol oxygenase |
| Tal12a | 121 | Bo8g100750 | 3,42E+00 | 1,29E-10 | VQ motif-containing protein [Source:TAIR;Acc:AT2G22880](projected from arabidopsis_thaliana,AT2G22880) | VQ motif-containing protein |
| Tal12a | 122 | Bo6g108930 | 3,42E+00 | 7,94E-32 |  | transcription factor NAM family |
| Tal12a | 123 | Bo4g137570 | 3,42E+00 | 2,65E-08 |  | putative protein |
| Tal12a | 124 | Bo9g090870 | 3,41E+00 | 3,91E-41 | alpha/beta-Hydrolases superfamily protein [Source:TAIR;Acc:AT4G10955](projected from arabidopsis_thaliana,AT4G10955) | fungal lipase-like domain, alpha/Beta hydrolase |
| Tal12a | 125 | Bo8g058230 | 3,37E+00 | 4,97E-07 |  | plant self-incompatibility S1 |
| Tal12a | 126 | Bo00689s040 | 3,35E+00 | 6,28E-06 |  | nucleic acid-binding protein |
| Tal12a | 127 | Bo04487s010 | 3,35E+00 | 2,11E-08 |  | Ctr copper transporter |
| Tal12a | 128 | Bo2g149730 | 3,35E+00 | 4,05E-36 |  | transcription factor WD40-like family |
| Tal12a | 129 | Bo8g081980 | 3,34E+00 | 3,11E-54 | unknown protein; FUNCTIONS IN: molecular_function unknown; INVOLVED IN: biological_process unknown; LOCATED IN: endomembrane system; EXPRESSED IN: 10 plant structures; EXPRESSED DURING: LP.04 four leaves visible, 4 anthesis, petal differentiation an /.../nsion stage; Has 28 Blast hits to 28 proteins in 7 species: Archae - 0; Bacteria - 0; Metazoa - 0; Fungi - 0; Plants - 28; Viruses - 0; Other Eukaryotes - 0 (source: NCBI BLink). [Source:TAIR;Acc:AT3G52480](projected from arabidopsis_thaliana,AT3G52480) | transmembrane protein |
| Tal12a | 130 | Bo2g155190 | 3,33E+00 | 7,02E-06 |  | hypothetical protein |
| Tal12a | 131 | Bo6g101010 | 3,33E+00 | 2,24E-27 | plastid movement impaired 2 [Source:TAIR;Acc:AT1G66480](projected from arabidopsis_thaliana,AT1G66480) | PADRE domain-containing protein |
| Tal12a | 132 | Bo3g130310 | 3,33E+00 | 1,73E-05 |  | transcription factor C2H2 family |
| Tal12a | 133 | Bo9g174300 | 3,33E+00 | 6,21E-37 |  | transcription factor & chromatin remodeling DDT family |
| Tal12a | 134 | Bo9g175080 | 3,32E+00 | 1,32E-12 |  | nepenthesin |
| Tal12a | 135 | Bo2g156160 | 3,32E+00 | 6,27E-12 |  | Phytocyanin domain, cupredoxin |
| Tal12a | 136 | Bo8g100320 | 3,32E+00 | 3,69E-11 |  | PADRE domain-containing protein |
| Tal12a | 137 | Bo3g023910 | 3,32E+00 | 6,72E-24 | NADPH/respiratory burst oxidase protein D [Source:TAIR;Acc:AT5G51060](projected from arabidopsis_thaliana,AT5G51060) | NAD(P)H oxidase (H2O2-forming) |
| Tal12a | 138 | Bo1g151660 | 3,31E+00 | 4,52E-07 |  | microtubule-binding protein TANGLED1 |
| Tal12a | 139 | Bo2g055820 | 3,29E+00 | 1,16E-35 |  | rubber elongation factor |
| Tal12a | 140 | Bo6g099030 | 3,29E+00 | 1,08E-57 |  | transcription factor CG1-CAMTA family |
| Tal12a | 141 | Bo2g011150 | 3,28E+00 | 3,27E-04 |  | transcription factor TIFY family |
| Tal12a | 142 | Bo04611s010 | 3,28E+00 | 8,64E-03 |  | triacylglycerol lipase |
| Tal12a | 143 | Bo9g025770 | 3,27E+00 | 1,05E-13 | P-loop containing nucleoside triphosphate hydrolases superfamily protein [Source:TAIR;Acc:AT1G03030](projected from arabidopsis_thaliana,AT1G03030) | P-loop containing nucleoside triphosphate hydrolase |
| Tal12a | 144 | Bo8g095590 | 3,27E+00 | 1,96E-11 |  | fasciclin-like arabinogalactan protein |
| Tal12a | 145 | Bo8g042120 | 3,26E+00 | 2,51E-05 | ARM repeat superfamily protein [Source:TAIR;Acc:AT4G15830](projected from arabidopsis_thaliana,AT4G15830) | armadillo-like helical protein |
| Tal12a | 146 | Bo1g011370 | 3,25E+00 | 3,89E-05 |  | flowering-promoting factor 1 |
| Tal12a | 147 | Bo5g130500 | 3,25E+00 | 1,81E-06 |  | carboxylesterase |
| Tal12a | 148 | Bo9g175640 | 3,25E+00 | 2,22E-05 |  | EF-Hand 1, calcium-binding protein |
| Tal12a | 149 | Bo9g032670 | 3,24E+00 | 3,81E-05 | P-loop containing nucleoside triphosphate hydrolases superfamily protein [Source:TAIR;Acc:AT1G64110](projected from arabidopsis_thaliana,AT1G64110) | AAA+ ATPase domain, ATPase, AAA-type, core, AAA ATPase, AAA+ lid domain-containing protein |
| Tal12a | 150 | Bo1g119260 | 3,22E+00 | 1,61E-06 | F-box and associated interaction domains-containing protein [Source:TAIR;Acc:AT3G17710](projected from arabidopsis_thaliana,AT3G17710) | F-box domain-containing protein |
| Tal12a | 151 | Bo8g050070 | 3,21E+00 | 2,85E-07 |  | small auxin-up RNA |
| Tal12a | 152 | Bo2g047970 | 3,21E+00 | 1,30E-07 |  | putative protein |
| Tal12a | 153 | Bo6g027580 | 3,20E+00 | 4,25E-05 |  | jacalin-like lectin domain-containing protein |
| Tal12a | 154 | Bo3g055640 | 3,19E+00 | 2,64E-39 | Pentatricopeptide repeat (PPR) superfamily protein [Source:TAIR;Acc:AT3G02490](projected from arabidopsis_thaliana,AT3G02490) | pentatricopeptide repeat-containing protein BIR6 |
| Tal12a | 155 | Bo2g117970 | 3,18E+00 | 3,40E-43 |  | actinidain |
| Tal12a | 156 | Bo1g129760 | 3,17E+00 | 4,69E-10 |  | putative protein |
| Tal12a | 157 | Bo01233s010 | 3,17E+00 | 2,50E-24 | embryo defective 2737 [Source:TAIR;Acc:AT5G53860](projected from arabidopsis_thaliana,AT5G53860) | putative protein |
| Tal12a | 158 | Bo3g169090 | 3,16E+00 | 7,76E-22 | Glycosyltransferase [Source:UniProtKB/TrEMBL;Acc:A0A0D3BLG5] | UDP-glucuronosyl/UDP-glucosyltransferase, UDP-glycosyltransferase family |
| Tal12a | 159 | Bo5g043440 | 3,14E+00 | 1,82E-04 |  | putative protein |
| Tal12a | 160 | Bo9g079550 | 3,13E+00 | 3,24E-16 | unknown protein; BEST Arabidopsis thaliana protein match is: unknown protein (TAIR:AT4G22560.1); Has 380 Blast hits to 380 proteins in 23 species: Archae - 0; Bacteria - 0; Metazoa - 1; Fungi - 4; Plants - 374; Viruses - 0; Other Eukaryotes - 1 (sou /.../CBI BLink). [Source:TAIR;Acc:AT4G12450](projected from arabidopsis_thaliana,AT4G12450) | putative protein |
| Tal12a | 161 | Bo8g114250 | 3,13E+00 | 8,06E-34 |  | Clp, repeat (R) domain, P-loop containing nucleoside triphosphate hydrolase |
| Tal12a | 162 | Bo2g131930 | 3,13E+00 | 2,33E-25 |  | RING-type E3 ubiquitin transferase transcription factor C2H2 family |
| Tal12a | 163 | Bo8g102350 | 3,12E+00 | 9,39E-05 |  | H/ACA ribonucleoprotein complex, subunit Nop10 |
| Tal12a | 164 | Bo5g071060 | 3,12E+00 | 4,79E-10 |  | protein-serine/threonine phosphatase |
| Tal12a | 165 | Bo7g065020 | 3,12E+00 | 8,28E-14 | ATP binding microtubule motor family protein [Source:TAIR;Acc:AT5G66310](projected from arabidopsis_thaliana,AT5G66310) | plus-end-directed kinesin ATPase |
| Tal12a | 166 | Bo9g058400 | 3,11E+00 | 7,86E-46 | Protein of unknown function, DUF547 [Source:TAIR;Acc:AT5G42690](projected from arabidopsis_thaliana,AT5G42690) | ternary complex factor MIP1, leucine-zipper |
| Tal12a | 167 | Bo9g163920 | 3,10E+00 | 8,78E-04 |  | protein SPIRAL1 |
| Tal12a | 168 | Bo7g043700 | 3,10E+00 | 1,60E-06 |  | Chaperone J-domain superfamily |
| Tal12a | 169 | Bo3g061400 | 3,10E+00 | 6,69E-12 |  | transcription factor C2C2-CO-like family |
| Tal12a | 170 | Bo5g078660 | 3,10E+00 | 3,23E-22 |  | protein kinase ULK-Fused family |
| Tal12a | 171 | Bo5g009380 | 3,09E+00 | 4,06E-19 | Nuclear transcription factor Y subunit C-3 [Source:Projected from Arabidopsis thaliana (AT1G54830) UniProtKB/Swiss-Prot;Acc:Q9ZVL3] | transcription factor Hap3/NF-YB family |
| Tal12a | 172 | Bo1g052890 | 3,08E+00 | 1,75E-04 |  | putative protein |
| Tal12a | 173 | Bo7g063740 | 3,08E+00 | 1,94E-04 |  | transcription factor interactor and regulator LisH family |
| Tal12a | 174 | Bo1g059360 | 3,07E+00 | 4,61E-32 |  | transcription factor C3H family |
| Tal12a | 175 | Bo5g115220 | 3,06E+00 | 9,58E-07 | Cysteine proteinases superfamily protein [Source:TAIR;Acc:AT3G43960](projected from arabidopsis_thaliana,AT3G43960) | actinidain |
| Tal12a | 176 | Bo4g154690 | 3,06E+00 | 1,24E-07 |  | putative protein |
| Tal12a | 177 | Bo3g036400 | 3,06E+00 | 2,22E-05 |  | transcription factor bHLH family |
| Tal12a | 178 | Bo4g041200 | 3,04E+00 | 4,18E-21 |  | trichome birefringence-like domain, PC-Esterase |
| Tal12a | 179 | Bo8g104890 | 3,04E+00 | 1,88E-10 |  | transcription factor TIFY family |
| Tal12a | 180 | Bo3g052640 | 3,04E+00 | 2,85E-04 |  | glutathione transferase transcription factor MYB-HB-like family |
| Tal12a | 181 | Bo3g081320 | 3,03E+00 | 2,05E-04 |  | bifunctional inhibitor/plant lipid transfer protein/seed storage helical |
| Tal12a | 182 | Bo3g165820 | 3,00E+00 | 3,24E-04 |  | hypothetical protein |
| Tal12a | 183 | Bo6g018670 | 2,99E+00 | 2,68E-04 |  | transcription factor MADS-type1 family |
| Tal12a | 184 | Bo3g053050 | 2,99E+00 | 1,89E-19 | tolB protein-related [Source:TAIR;Acc:AT4G01870](projected from arabidopsis_thaliana,AT4G01870) | transcription factor WD40-like family |
| Tal12a | 185 | Bo9g151700 | 2,98E+00 | 9,65E-05 |  | hypothetical protein |
| Tal12a | 186 | Bo7g080870 | 2,98E+00 | 7,52E-07 |  | ribonuclease T2 |
| Tal12a | 187 | Bo4g023860 | 2,96E+00 | 2,20E-13 |  | RING-type E3 ubiquitin transferase chromatin regulator PHD family |
| Tal12a | 188 | Bo2g007480 | 2,96E+00 | 1,27E-04 | Sulfotransferase [Source:UniProtKB/TrEMBL;Acc:A0A0D3AI14] | hydroxyjasmonate sulfotransferase |
| Tal12a | 189 | Bo8g033750 | 2,95E+00 | 2,54E-30 |  | chlorophyll(ide) b reductase |
| Tal12a | 190 | Bo6g080700 | 2,95E+00 | 6,34E-04 |  | sister chromatid cohesion protein Pds5 |
| Tal12a | 191 | Bo7g098530 | 2,92E+00 | 4,02E-08 |  | putative protein |
| Tal12a | 192 | Bo1g155130 | 2,90E+00 | 5,44E-15 |  | protein-tyrosine-phosphatase |
| Tal12a | 193 | Bo3g022080 | 2,90E+00 | 7,26E-04 |  | GH3 family protein |
| Tal12a | 194 | Bo7g105020 | 2,89E+00 | 2,55E-04 |  | putative protein |
| Tal12a | 195 | Bo5g122540 | 2,89E+00 | 1,17E-05 | Molecular chaperone Hsp40/DnaJ family protein [Source:TAIR;Acc:AT3G17830](projected from arabidopsis_thaliana,AT3G17830) | Heat shock protein DnaJ, cysteine-rich |
| Tal12a | 196 | Bo00834s100 | 2,88E+00 | 5,61E-03 |  | DNA/RNA-binding protein Alba |
| Tal12a | 197 | Bo6g119050 | 2,87E+00 | 3,87E-08 | RING/U-box superfamily protein [Source:TAIR;Acc:AT1G74870](projected from arabidopsis_thaliana,AT1G74870) | RING-type E3 ubiquitin transferase transcription factor C2H2 family |
| Tal12a | 198 | Bo4g184570 | 2,87E+00 | 1,01E-03 | Heavy metal transport/detoxification superfamily protein [Source:TAIR;Acc:AT2G35730](projected from arabidopsis_thaliana,AT2G35730) | heavy metal-associated domain, HMA, heavy metal-associated domain superfamily |
| Tal12a | 199 | Bo6g119180 | 2,86E+00 | 7,33E-04 | GNS1/SUR4 membrane protein family [Source:TAIR;Acc:AT1G75000](projected from arabidopsis_thaliana,AT1G75000) | ELO family protein |
| Tal12a | 200 | Bo2g041160 | 2,85E+00 | 2,79E-29 |  | glucan endo-1,3-beta-D-glucosidase |
| Tal12a | 201 | Bo5g009840 | 2,85E+00 | 3,36E-06 |  | dihydroflavanol 4-reductase |
| Tal12a | 202 | Bo9g020340 | 2,84E+00 | 1,75E-04 |  | putative protein |
| Tal12a | 203 | Bo7g054170 | 2,83E+00 | 1,75E-03 |  | hypothetical protein |
| Tal12a | 204 | Bo6g061060 | 2,83E+00 | 2,28E-07 |  | ribosomal RNA-processing protein |
| Tal12a | 205 | Bo3g027080 | 2,83E+00 | 6,40E-14 | phosphate transporter 1;5 [Source:TAIR;Acc:AT2G32830](projected from arabidopsis_thaliana,AT2G32830) | major facilitator, sugar transporter, major facilitator superfamily |
| Tal12a | 206 | Bo9g160000 | 2,83E+00 | 1,32E-03 |  | putative protein |
| Tal12a | 207 | Bo4g157500 | 2,80E+00 | 7,35E-06 |  | hydroxyproline O-arabinosyltransferase |
| Tal12a | 208 | Bo3g015460 | 2,80E+00 | 1,56E-07 | Glycosyl hydrolase family protein [Source:TAIR;Acc:AT5G20940](projected from arabidopsis_thaliana,AT5G20940) | glucan 1,3-beta-glucosidase |
| Tal12a | 209 | Bo4g125390 | 2,79E+00 | 4,28E-05 | RING-type E3 ubiquitin transferase [Source:UniProtKB/TrEMBL;Acc:A0A0D3BYI3] | RING-type E3 ubiquitin transferase transcription regulator Rcd1-like family |
| Tal12a | 210 | Bo3g011970 | 2,79E+00 | 7,03E-16 |  | LRAT domain-containing protein |
| Tal12a | 211 | Bo3g103500 | 2,78E+00 | 2,35E-03 |  | putative protein |
| Tal12a | 212 | Bo4g154660 | 2,78E+00 | 6,79E-05 |  | cytochrome P450 |
| Tal12a | 213 | Bo5g123740 | 2,78E+00 | 1,47E-30 |  | condensin II complex subunit H2, scpA-like, condensin II complex subunit H2, middle |
| Tal12a | 214 | Bo4g198550 | 2,77E+00 | 1,18E-17 |  | long-chain-fatty-acid--CoA ligase |
| Tal12a | 215 | Bo6g035930 | 2,77E+00 | 2,66E-05 | unknown protein; BEST Arabidopsis thaliana protein match is: unknown protein (TAIR:AT3G12970.1); Has 3011 Blast hits to 958 proteins in 192 species: Archae - 0; Bacteria - 193; Metazoa - 479; Fungi - 286; Plants - 158; Viruses - 8; Other Eukaryotes /.../ (source: NCBI BLink). [Source:TAIR;Acc:AT1G56020](projected from arabidopsis_thaliana,AT1G56020) | putative protein |
| Tal12a | 216 | Bo4g098840 | 2,77E+00 | 2,46E-34 |  | protein-serine/threonine phosphatase |
| Tal12a | 217 | Bo2g010640 | 2,77E+00 | 1,52E-13 |  | alpha/Beta hydrolase |
| Tal12a | 218 | Bo3g032100 | 2,76E+00 | 4,28E-05 |  | GDP-fucose protein O-fucosyltransferase |
| Tal12a | 219 | Bo3g162530 | 2,75E+00 | 3,65E-13 |  | organic solute transporter subunit alpha/Transmembrane protein |
| Tal12a | 220 | Bo9g168900 | 2,75E+00 | 3,63E-03 |  | transcription factor TIFY family |
| Tal12a | 221 | Bo5g021160 | 2,74E+00 | 3,95E-19 | Cobalamin biosynthesis CobW-like protein [Source:TAIR;Acc:AT1G15730](projected from arabidopsis_thaliana,AT1G15730) | cobW/HypB/UreG, nucleotide-binding domain, Zinc chaperone CobW-like protein |
| Tal12a | 222 | Bo6g047940 | 2,74E+00 | 2,71E-07 | RING/U-box superfamily protein [Source:TAIR;Acc:AT1G33480](projected from arabidopsis_thaliana,AT1G33480) | RING-type E3 ubiquitin transferase transcription factor C2H2 family |
| Tal12a | 223 | Bo9g160140 | 2,74E+00 | 2,82E-15 |  | maltose excess protein |
| Tal12a | 224 | Bo2g092490 | 2,73E+00 | 6,93E-05 |  | aminocyclopropanecarboxylate oxidase |
| Tal12a | 225 | Bo3g031950 | 2,73E+00 | 1,31E-06 |  | oxidoreductase |
| Tal12a | 226 | Bo1g017350 | 2,72E+00 | 1,50E-06 | ROP-interactive CRIB motif-containing protein 7 [Source:TAIR;Acc:AT4G28560](projected from arabidopsis_thaliana,AT4G28560) | leucine-rich repeat domain superfamily |
| Tal12a | 227 | Bo9g025750 | 2,72E+00 | 1,62E-12 | Diacylglycerol kinase [Source:UniProtKB/TrEMBL;Acc:A0A0D3E378] | diacylglycerol kinase (ATP) |
| Tal12a | 228 | Bo6g029520 | 2,72E+00 | 1,84E-05 |  | phosphoenolpyruvate carboxylase |
| Tal12a | 229 | Bo4g198690 | 2,71E+00 | 2,54E-08 | Integrase-type DNA-binding superfamily protein [Source:TAIR;Acc:AT2G47520](projected from arabidopsis_thaliana,AT2G47520) | transcription factor AP2-EREBP family |
| Tal12a | 230 | Bo9g007570 | 2,70E+00 | 8,37E-12 | Protein phosphatase 2C family protein [Source:TAIR;Acc:AT4G03415](projected from arabidopsis_thaliana,AT4G03415) | protein-serine/threonine phosphatase |
| Tal12a | 231 | Bo8g101030 | 2,70E+00 | 8,68E-06 | S-adenosyl-L-methionine-dependent methyltransferases superfamily protein [Source:TAIR;Acc:AT3G44840](projected from arabidopsis_thaliana,AT3G44840) | juvenile hormone-III synthase |
| Tal12a | 232 | Bo1g129910 | 2,69E+00 | 7,65E-19 |  | trichome birefringence-like domain, PC-Esterase |
| Tal12a | 233 | Bo9g148400 | 2,69E+00 | 3,89E-17 | Fatty acyl-CoA reductase [Source:UniProtKB/TrEMBL;Acc:A0A0D3EDD1] | oxidoreductase |
| Tal12a | 234 | Bo6g029510 | 2,69E+00 | 1,13E-38 |  | phosphoenolpyruvate carboxylase |
| Tal12a | 235 | Bo3g066460 | 2,69E+00 | 2,76E-06 |  | protein gravitropic in the light 1 |
| Tal12a | 236 | Bo6g049370 | 2,68E+00 | 2,44E-04 | Protein kinase superfamily protein [Source:TAIR;Acc:AT1G33260](projected from arabidopsis_thaliana,AT1G33260) | protein kinase RLK-Pelle-RLCK-XIII family |
| Tal12a | 237 | Bo6g044230 | 2,68E+00 | 3,75E-07 |  | carboxylic ester hydrolase |
| Tal12a | 238 | Bo1g105730 | 2,66E+00 | 3,62E-06 |  | multi antimicrobial extrusion protein |
| Tal12a | 239 | Bo4g025890 | 2,66E+00 | 2,84E-20 | Delta-1-pyrroline-5-carboxylate synthase [Source:UniProtKB/TrEMBL;Acc:A0A0D3BQG1] | oxidoreductase, glutamate 5-kinase |
| Tal12a | 240 | Bo1g020370 | 2,66E+00 | 5,45E-03 |  | (+)-abscisic acid 8'-hydroxylase |
| Tal12a | 241 | Bo10177s010 | 2,66E+00 | 5,71E-05 |  | non-specific serine/threonine protein kinase |
| Tal12a | 242 | Bo7g058300 | 2,65E+00 | 1,52E-13 |  | F-box protein AUF1 |
| Tal12a | 243 | Bo1g005100 | 2,65E+00 | 5,72E-03 | Patatin [Source:UniProtKB/TrEMBL;Acc:A0A0D3A1D4] | phospholipase A2 |
| Tal12a | 244 | Bo8g080780 | 2,65E+00 | 4,95E-04 |  | putative protein |
| Tal12a | 245 | Bo7g104790 | 2,65E+00 | 2,37E-04 |  | carboxylic ester hydrolase |
| Tal12a | 246 | Bo9g166010 | 2,64E+00 | 7,93E-03 |  | putative protein |
| Tal12a | 247 | Bo5g025160 | 2,64E+00 | 9,89E-34 |  | Golgin subfamily A member 5 protein |
| Tal12a | 248 | Bo9g176510 | 2,64E+00 | 9,66E-03 |  | sucrose/H+ symporter, plant, MFS transporter superfamily |
| Tal12a | 249 | Bo3g107930 | 2,63E+00 | 1,28E-23 | unknown protein; BEST Arabidopsis thaliana protein match is: unknown protein (TAIR:AT5G51850.1); Has 381 Blast hits to 359 proteins in 81 species: Archae - 0; Bacteria - 16; Metazoa - 101; Fungi - 21; Plants - 99; Viruses - 3; Other Eukaryotes - 141 /.../ce: NCBI BLink). [Source:TAIR;Acc:AT5G62170](projected from arabidopsis_thaliana,AT5G62170) | putative protein |
| Tal12a | 250 | Bo3g093980 | 2,63E+00 | 3,44E-16 |  | transcription factor bZIP family |
| Tal12a | 251 | Bo4g119580 | 2,62E+00 | 2,66E-25 |  | non-specific serine/threonine protein kinase |
| Tal12a | 252 | Bo1g019970 | 2,62E+00 | 3,15E-33 | AAA-type ATPase family protein [Source:TAIR;Acc:AT4G18820](projected from arabidopsis_thaliana,AT4G18820) | DNA-directed DNA polymerase |
| Tal12a | 253 | Bo4g125450 | 2,62E+00 | 2,55E-15 | FASCICLIN-like arabinogalactan protein 15 precursor [Source:TAIR;Acc:AT3G52370](projected from arabidopsis_thaliana,AT3G52370) | fasciclin-like arabinogalactan protein 15/16/17/18 |
| Tal12a | 254 | Bo6g031100 | 2,61E+00 | 7,83E-14 |  | legume lectin domain, concanavalin A-like lectin/glucanase domain superfamily |
| Tal12a | 255 | Bo9g151710 | 2,61E+00 | 3,50E-04 |  | glutathione transferase transcription factor MYB-HB-like family |
| Tal12a | 256 | Bo1g022550 | 2,61E+00 | 4,88E-17 |  | tetrahydroberberine oxidase |
| Tal12a | 257 | Bo1g039280 | 2,60E+00 | 5,61E-19 |  | palmitoyl-protein hydrolase |
| Tal12a | 258 | Bo3g147640 | 2,60E+00 | 7,54E-04 |  | transcription factor TIFY family |
| Tal12a | 259 | Bo3g068560 | 2,60E+00 | 6,62E-06 |  | transcription factor C2C2-GATA family |
| Tal12a | 260 | Bo7g109750 | 2,59E+00 | 9,85E-05 | Pectate lyase [Source:UniProtKB/TrEMBL;Acc:A0A0D3DFX6] | pectate lyase |
| Tal12a | 261 | Bo7g049930 | 2,59E+00 | 4,59E-05 |  | CRIB domain-containing protein RIC2/4 |
| Tal12a | 262 | Bo6g018460 | 2,59E+00 | 6,51E-09 |  | phosphoenolpyruvate carboxylase |
| Tal12a | 263 | Bo9g117710 | 2,59E+00 | 4,46E-30 |  | transcription factor WD40-like family |
| Tal12a | 264 | Bo1g023850 | 2,58E+00 | 4,88E-09 |  | tetrahydroberberine oxidase |
| Tal12a | 265 | Bo2g034070 | 2,57E+00 | 2,17E-09 | ATP-dependent 6-phosphofructokinase [Source:UniProtKB/TrEMBL;Acc:A0A0D3ALN7] | 6-phosphofructokinase |
| Tal12a | 266 | Bo2g010300 | 2,57E+00 | 3,96E-04 | Glutamate receptor [Source:UniProtKB/TrEMBL;Acc:A0A0D3AIJ7] | periplasmic binding protein-like I |
| Tal12a | 267 | Bo3g012520 | 2,56E+00 | 2,51E-09 | Glutamate decarboxylase [Source:UniProtKB/TrEMBL;Acc:A0A0D3B1M0] | glutamate decarboxylase |
| Tal12a | 268 | Bo03992s010 | 2,56E+00 | 1,92E-03 |  | putative protein |
| Tal12a | 269 | Bo9g005710 | 2,56E+00 | 6,44E-10 | Protein of unknown function, DUF538 [Source:TAIR;Acc:AT4G02360](projected from arabidopsis_thaliana,AT4G02360) | Putative-like superfamily protein |
| Tal12a | 270 | Bo9g117350 | 2,56E+00 | 1,69E-15 |  | transcription factor MYB-HB-like family |
| Tal12a | 271 | Bo6g030940 | 2,55E+00 | 1,42E-11 | NAC domain containing protein 19 [Source:TAIR;Acc:AT1G52890](projected from arabidopsis_thaliana,AT1G52890) | transcription factor NAM family |
| Tal12a | 272 | Bo8g016830 | 2,55E+00 | 1,85E-08 |  | transmembrane protein |
| Tal12a | 273 | Bo7g114400 | 2,55E+00 | 4,45E-03 | Xyloglucan endotransglucosylase/hydrolase [Source:UniProtKB/TrEMBL;Acc:A0A0D3DGZ2] | xyloglucan:xyloglucosyl transferase |
| Tal12a | 274 | Bo7g003210 | 2,55E+00 | 5,25E-11 |  | mannan endo-1,4-beta-mannosidase |
| Tal12a | 275 | Bo01213s020 | 2,55E+00 | 4,92E-04 |  | protein WAVE-DAMPENED 2 |
| Tal12a | 276 | Bo3g012560 | 2,54E+00 | 9,38E-03 |  | PADRE domain-containing protein |
| Tal12a | 277 | Bo6g101070 | 2,53E+00 | 8,37E-03 |  | juvenile hormone-III synthase |
| Tal12a | 278 | Bo5g002460 | 2,53E+00 | 5,61E-05 | 1-amino-cyclopropane-1-carboxylate synthase 2 [Source:TAIR;Acc:AT1G01480](projected from arabidopsis_thaliana,AT1G01480) | 1-aminocyclopropane-1-carboxylate synthase |
| Tal12a | 279 | Bo3g057330 | 2,52E+00 | 7,07E-04 |  | C2 domain-containing protein |
| Tal12a | 280 | Bo7g092740 | 2,51E+00 | 5,41E-34 |  | transcription factor WD40-like family |
| Tal12a | 281 | Bo4g046030 | 2,51E+00 | 5,03E-04 |  | mitogen-activated protein kinase kinase kinase STE-STE11 family |
| Tal12a | 282 | Bo4g167920 | 2,51E+00 | 2,29E-17 |  | rlpA-like protein, double-psi beta-barrel |
| Tal12a | 283 | Bo2g119360 | 2,51E+00 | 1,07E-12 | actin-related protein 9 [Source:TAIR;Acc:AT5G43500](projected from arabidopsis_thaliana,AT5G43500) | Actin family, ATPase, nucleotide binding domain-containing protein |
| Tal12a | 284 | Bo3g179730 | 2,50E+00 | 1,83E-22 |  | sulfurtransferase |
| Tal12a | 285 | Bo6g077870 | 2,50E+00 | 1,16E-10 | Glucose-6-phosphate 1-epimerase [Source:UniProtKB/TrEMBL;Acc:A0A0D3CV70] | isomerase |
| Tal12a | 286 | Bo8g030170 | 2,49E+00 | 8,23E-06 | Xyloglucan endotransglucosylase/hydrolase [Source:UniProtKB/TrEMBL;Acc:A0A0D3DKY0] | xyloglucan:xyloglucosyl transferase, xyloglucan-specific endo-beta-1,4-glucanase |
| Tal12a | 287 | Bo9g036870 | 2,48E+00 | 3,67E-19 |  | pectinesterase inhibitor domain-containing protein |
| Tal12a | 288 | Bo3g018270 | 2,48E+00 | 8,67E-04 | cyclin T1;1 [Source:TAIR;Acc:AT1G35440](projected from arabidopsis_thaliana,AT1G35440) | cyclin domain-containing protein |
| Tal12a | 289 | Bo6g081600 | 2,47E+00 | 3,55E-08 |  | VQ motif-containing protein 1/10 |
| Tal12a | 290 | Bo8g002860 | 2,47E+00 | 3,62E-09 |  | non-specific serine/threonine protein kinase |
| Tal12a | 291 | Bo4g157490 | 2,46E+00 | 9,74E-06 |  | hydroxyproline O-arabinosyltransferase |
| Tal12a | 292 | Bo7g064080 | 2,45E+00 | 7,99E-03 | mitogen-activated protein kinase kinase kinase 19 [Source:Projected from Arabidopsis thaliana (AT5G67080) TAIR;Acc:AT5G67080] | mitogen-activated protein kinase kinase kinase STE-STE11 family |
| Tal12a | 293 | Bo8g110850 | 2,45E+00 | 1,84E-03 |  | transcription factor TGA like domain-containing protein |
| Tal12a | 294 | Bo7g109130 | 2,44E+00 | 8,04E-15 | TRICHOME BIREFRINGENCE-LIKE 24 [Source:TAIR;Acc:AT4G23790](projected from arabidopsis_thaliana,AT4G23790) | trichome birefringence-like domain, PC-Esterase |
| Tal12a | 295 | Bo2g152920 | 2,43E+00 | 2,65E-26 |  | AT-hook motif nuclear-localized protein |
| Tal12a | 296 | Bo6g122430 | 2,43E+00 | 6,12E-11 | Mitochondrial transcription termination factor family protein [Source:TAIR;Acc:AT1G78930](projected from arabidopsis_thaliana,AT1G78930) | transcription regulator mTERF family |
| Tal12a | 297 | Bo9g135020 | 2,42E+00 | 4,67E-11 | P-loop containing nucleoside triphosphate hydrolases superfamily protein [Source:TAIR;Acc:AT5G58370](projected from arabidopsis_thaliana,AT5G58370) | small GTP-binding protein |
| Tal12a | 298 | Bo3g035870 | 2,41E+00 | 1,23E-05 |  | putative protein |
| Tal12a | 299 | Bo8g110860 | 2,41E+00 | 4,55E-15 | Glutamyl-tRNA reductase [Source:UniProtKB/TrEMBL;Acc:A0A0D3DY87] | glutamyl-tRNA reductase |
| Tal12a | 300 | Bo1g022520 | 2,41E+00 | 8,67E-04 |  | tetrahydroberberine oxidase |
| Tal14c | 1 | Bo01007s020 | 1,01E+01 | 1,01E-122 |  | cytochrome P450 |
| Tal14c | 2 | Bo9g058330 | 9,79E+00 | 2,14E-86 |  | acetylajmaline esterase |
| Tal14c | 3 | Bo3g176080 | 9,41E+00 | 1,25E-59 |  | nucleotide diphosphatase |
| Tal14c | 4 | Bo10610s010 | 9,18E+00 | 1,85E-65 |  | nucleotide diphosphatase |
| Tal14c | 5 | Bo3g153450 | 8,56E+00 | 5,84E-68 |  | cytochrome P450 |
| Tal14c | 6 | Bo9g075840 | 8,48E+00 | 3,46E-89 |  | allene-oxide cyclase |
| Tal14c | 7 | Bo3g034260 | 8,30E+00 | 1,83E-29 |  | jacalin-like lectin domain-containing protein |
| Tal14c | 8 | Bo2g023630 | 8,07E+00 | 2,98E-85 | Integrase-type DNA-binding superfamily protein [Source:TAIR;Acc:AT5G21960](projected from arabidopsis_thaliana,AT5G21960) | transcription factor AP2-EREBP family |
| Tal14c | 9 | Bo9g017680 | 7,87E+00 | 3,23E-61 |  | jasmonoyl-L-amino acid 12-hydroxylase |
| Tal14c | 10 | Bo6g006880 | 7,74E+00 | 2,54E-60 |  | triacylglycerol lipase |
| Tal14c | 11 | Bo6g028110 | 7,56E+00 | 8,49E-31 |  | putative protein |
| Tal14c | 12 | Bo20985s010 | 7,40E+00 | 1,51E-06 |  | proteinase inhibitor I3, Kunitz legume, kunitz inhibitor STI-like superfamily |
| Tal14c | 13 | Bo7g082020 | 7,35E+00 | 7,78E-69 |  | allene-oxide cyclase |
| Tal14c | 14 | Bo3g022260 | 7,32E+00 | 1,75E-51 |  | START domain-containing protein |
| Tal14c | 15 | Bo3g064960 | 7,27E+00 | 8,43E-46 |  | chitinase |
| Tal14c | 16 | Bo8g077060 | 7,11E+00 | 4,43E-60 |  | putative protein |
| Tal14c | 17 | Bo4g187470 | 7,08E+00 | 2,63E-23 |  | flavanone 3-dioxygenase |
| Tal14c | 18 | Bo4g115410 | 7,01E+00 | 1,22E-16 |  | flavonol synthase |
| Tal14c | 19 | Bo4g021650 | 7,00E+00 | 8,19E-26 |  | neprosin |
| Tal14c | 20 | Bo3g004370 | 7,00E+00 | 6,00E-59 |  | peptide-methionine (S)-S-oxide reductase |
| Tal14c | 21 | Bo9g010640 | 6,96E+00 | 3,36E-43 | Kinesin-like protein [Source:UniProtKB/TrEMBL;Acc:A0A0D3E159] | minus-end-directed kinesin ATPase |
| Tal14c | 22 | Bo3g178190 | 6,95E+00 | 1,27E-24 |  | acid phosphatase |
| Tal14c | 23 | Bo7g054590 | 6,92E+00 | 1,38E-57 |  | glutathione transferase |
| Tal14c | 24 | Bo9g176740 | 6,92E+00 | 5,39E-35 |  | phenylacetaldehyde oxime monooxygenase |
| Tal14c | 25 | Bo4g194410 | 6,79E+00 | 1,79E-45 |  | scorpion long chain toxin/defensin, knottin, scorpion toxin-like superfamily |
| Tal14c | 26 | Bo6g028090 | 6,75E+00 | 1,56E-31 |  | beta-glucosidase |
| Tal14c | 27 | Bo7g082010 | 6,70E+00 | 1,50E-44 |  | allene-oxide cyclase |
| Tal14c | 28 | Bo4g117140 | 6,56E+00 | 7,23E-43 | ABC-2 type transporter family protein [Source:TAIR;Acc:AT3G55100](projected from arabidopsis_thaliana,AT3G55100) | ABC-type molybdate transporter |
| Tal14c | 29 | Bo4g187150 | 6,55E+00 | 1,07E-30 |  | fatty acid hydroxylase |
| Tal14c | 30 | Bo01178s040 | 6,55E+00 | 3,97E-24 |  | tetrahydroberberine oxidase |
| Tal14c | 31 | Bo6g006870 | 6,53E+00 | 1,38E-57 |  | triacylglycerol lipase |
| Tal14c | 32 | Bo4g187140 | 6,47E+00 | 7,76E-21 |  | very-long-chain aldehyde decarbonylase CER1-like protein |
| Tal14c | 33 | Bo5g148850 | 6,46E+00 | 1,36E-38 | non-specific phospholipase C3 [Source:TAIR;Acc:AT3G03520](projected from arabidopsis_thaliana,AT3G03520) | phospholipase C |
| Tal14c | 34 | Bo8g099700 | 6,46E+00 | 2,87E-26 | Nucleotide-diphospho-sugar transferases superfamily protein [Source:TAIR;Acc:AT5G30500](projected from arabidopsis_thaliana,AT5G30500) | inositol 3-alpha-galactosyltransferase |
| Tal14c | 35 | Bo8g101030 | 6,44E+00 | 1,59E-27 | S-adenosyl-L-methionine-dependent methyltransferases superfamily protein [Source:TAIR;Acc:AT3G44840](projected from arabidopsis_thaliana,AT3G44840) | juvenile hormone-III synthase |
| Tal14c | 36 | Bo27042s010 | 6,41E+00 | 7,65E-38 |  | nucleotide diphosphatase |
| Tal14c | 37 | Bo2g011150 | 6,39E+00 | 2,70E-16 |  | transcription factor TIFY family |
| Tal14c | 38 | Bo2g159220 | 6,32E+00 | 9,33E-57 |  | acid phosphatase |
| Tal14c | 39 | Bo5g042850 | 6,21E+00 | 1,05E-11 | Flavin-dependent oxidoreductase FOX2 [Source:Projected from Arabidopsis thaliana (AT1G26390) UniProtKB/Swiss-Prot;Acc:Q9FZC5] | oxidoreductase |
| Tal14c | 40 | Bo9g168900 | 6,12E+00 | 7,39E-15 |  | transcription factor TIFY family |
| Tal14c | 41 | Bo2g007480 | 6,10E+00 | 2,18E-21 | Sulfotransferase [Source:UniProtKB/TrEMBL;Acc:A0A0D3AI14] | hydroxyjasmonate sulfotransferase |
| Tal14c | 42 | Bo5g020510 | 6,08E+00 | 2,05E-05 |  | gamma-glutamyl-gamma-aminobutyrate hydrolase |
| Tal14c | 43 | Bo5g021790 | 6,05E+00 | 3,64E-15 |  | major facilitator, sugar transporter, major facilitator superfamily |
| Tal14c | 44 | Bo3g004380 | 6,03E+00 | 1,29E-16 |  | peptide-methionine (S)-S-oxide reductase |
| Tal14c | 45 | Bo1g004980 | 5,98E+00 | 2,56E-42 | mitogen-activated protein kinase kinase kinase 21 [Source:TAIR;Acc:AT4G36950](projected from arabidopsis_thaliana,AT4G36950) | mitogen-activated protein kinase kinase kinase STE-STE11 family |
| Tal14c | 46 | Bo00722s140 | 5,96E+00 | 8,08E-23 |  | harbinger transposase-derived protein |
| Tal14c | 47 | Bo4g198550 | 5,94E+00 | 4,20E-82 |  | long-chain-fatty-acid--CoA ligase |
| Tal14c | 48 | Bo5g110090 | 5,93E+00 | 1,61E-39 |  | cytochrome P450 |
| Tal14c | 49 | Bo3g123620 | 5,93E+00 | 2,23E-26 |  | transcription factor C2H2 family |
| Tal14c | 50 | Bo8g100840 | 5,92E+00 | 5,83E-10 | HXXXD-type acyl-transferase family protein [Source:TAIR;Acc:AT5G47950](projected from arabidopsis_thaliana,AT5G47950) | anthocyanidin 3-O-glucoside 6''-O-acyltransferase |
| Tal14c | 51 | Bo2g075340 | 5,92E+00 | 8,04E-25 |  | transferase |
| Tal14c | 52 | Bo4g169210 | 5,90E+00 | 2,98E-13 |  | oxidoreductase |
| Tal14c | 53 | Bo1g141170 | 5,83E+00 | 3,17E-20 |  | anthocyanidin synthase |
| Tal14c | 54 | Bo7g116150 | 5,75E+00 | 1,71E-48 |  | galacturonan 1,4-alpha-galacturonidase |
| Tal14c | 55 | Bo5g134680 | 5,75E+00 | 1,69E-18 |  | leucine-rich repeat-containing, plant-type, leucine-rich repeat domain superfamily |
| Tal14c | 56 | Bo9g082080 | 5,74E+00 | 1,76E-14 |  | putative protein |
| Tal14c | 57 | Bo7g101110 | 5,73E+00 | 1,95E-32 |  | xanthoxin dehydrogenase |
| Tal14c | 58 | Bo6g032890 | 5,72E+00 | 3,17E-24 |  | GDSL lipase/esterase, SGNH hydrolase superfamily |
| Tal14c | 59 | Bo4g039150 | 5,70E+00 | 1,55E-18 |  | leucine-rich repeat-containing, plant-type, leucine-rich repeat domain superfamily |
| Tal14c | 60 | Bo1g057020 | 5,69E+00 | 3,66E-23 |  | anthocyanidin 3-O-glucoside 6''-O-acyltransferase |
| Tal14c | 61 | Bo2g107500 | 5,65E+00 | 1,30E-22 | unknown protein; FUNCTIONS IN: molecular_function unknown; INVOLVED IN: biological_process unknown; LOCATED IN: chloroplast; BEST Arabidopsis thaliana protein match is: unknown protein (TAIR:AT4G11910.1); Has 30201 Blast hits to 17322 proteins in 78 /.../ies: Archae - 12; Bacteria - 1396; Metazoa - 17338; Fungi - 3422; Plants - 5037; Viruses - 0; Other Eukaryotes - 2996 (source: NCBI BLink). [Source:TAIR;Acc:AT4G11911](projected from arabidopsis_thaliana,AT4G11911) | staygreen protein |
| Tal14c | 62 | Bo6g083740 | 5,64E+00 | 1,57E-33 |  | O-methyltransferase domain, plant methyltransferase dimerisation domain-containing protein |
| Tal14c | 63 | Bo6g116660 | 5,59E+00 | 1,67E-48 |  | proteinase inhibitor I3, Kunitz legume, kunitz inhibitor STI-like superfamily |
| Tal14c | 64 | Bo9g021840 | 5,58E+00 | 3,64E-15 |  | transcription factor interactor and regulator LIM family |
| Tal14c | 65 | Bo6g101070 | 5,57E+00 | 1,25E-15 |  | juvenile hormone-III synthase |
| Tal14c | 66 | Bo3g004090 | 5,56E+00 | 3,69E-44 |  | leucine-rich repeat-containing, plant-type, leucine-rich repeat domain superfamily |
| Tal14c | 67 | Bo2g127540 | 5,55E+00 | 1,15E-28 |  | hypothetical protein |
| Tal14c | 68 | Bo2g075280 | 5,54E+00 | 1,09E-22 |  | transferase |
| Tal14c | 69 | Bo9g175500 | 5,52E+00 | 5,67E-29 |  | peptide-methionine (S)-S-oxide reductase |
| Tal14c | 70 | Bo8g100690 | 5,44E+00 | 4,48E-16 |  | PLAT/LH2 domain-containing protein |
| Tal14c | 71 | Bo2g023440 | 5,40E+00 | 4,18E-22 |  | Late embryogenesis abundant protein, LEA_2 subgroup |
| Tal14c | 72 | Bo5g020810 | 5,40E+00 | 4,88E-09 |  | glyoxalase/Bleomycin resistance protein/Dihydroxybiphenyl dioxygenase |
| Tal14c | 73 | Bo6g120830 | 5,39E+00 | 2,86E-33 |  | O-methyltransferase domain, plant methyltransferase dimerisation domain-containing protein |
| Tal14c | 74 | Bo3g153850 | 5,38E+00 | 9,92E-27 | Heterodimeric geranylgeranyl pyrophosphate synthase large subunit 1, chloroplastic [Source:Projected from Arabidopsis thaliana (AT4G36810) UniProtKB/Swiss-Prot;Acc:P34802] | transferase |
| Tal14c | 75 | Bo5g149890 | 5,37E+00 | 2,73E-31 |  | (Z)-3-hexen-1-ol acetyltransferase |
| Tal14c | 76 | Bo9g161390 | 5,37E+00 | 8,29E-20 | Glutamate decarboxylase [Source:UniProtKB/TrEMBL;Acc:A0A0D3EET8] | glutamate decarboxylase |
| Tal14c | 77 | Bo9g175060 | 5,35E+00 | 3,07E-12 | Sulfotransferase [Source:UniProtKB/TrEMBL;Acc:A0A0D3EH11] | hydroxyjasmonate sulfotransferase |
| Tal14c | 78 | Bo9g161480 | 5,34E+00 | 4,00E-25 |  | glutathione transferase |
| Tal14c | 79 | Bo6g122000 | 5,34E+00 | 1,01E-27 |  | 9-cis-epoxycarotenoid dioxygenase |
| Tal14c | 80 | Bo5g111090 | 5,29E+00 | 3,97E-17 |  | cytochrome P450 |
| Tal14c | 81 | Bo8g067210 | 5,28E+00 | 5,77E-22 | Lipoxygenase [Source:UniProtKB/TrEMBL;Acc:A0A0D3DQ07] | linoleate 13S-lipoxygenase |
| Tal14c | 82 | Bo9g075870 | 5,27E+00 | 3,30E-19 | allene oxide cyclase 3 [Source:TAIR;Acc:AT3G25780](projected from arabidopsis_thaliana,AT3G25780) | allene-oxide cyclase |
| Tal14c | 83 | Bo1g014720 | 5,27E+00 | 3,38E-29 |  | transcription factor bHLH family |
| Tal14c | 84 | Bo5g009840 | 5,21E+00 | 1,49E-29 |  | dihydroflavanol 4-reductase |
| Tal14c | 85 | Bo8g090360 | 5,21E+00 | 2,03E-12 |  | flavonol synthase |
| Tal14c | 86 | Bo6g079880 | 5,19E+00 | 3,78E-14 | Lipoxygenase [Source:UniProtKB/TrEMBL;Acc:A0A0D3CVH2] | linoleate 13S-lipoxygenase |
| Tal14c | 87 | Bo9g154850 | 5,16E+00 | 6,95E-18 | Terpenoid cyclases/Protein prenyltransferases superfamily protein [Source:TAIR;Acc:AT5G44630](projected from arabidopsis_thaliana,AT5G44630) | lyase |
| Tal14c | 88 | Bo5g015010 | 5,16E+00 | 2,49E-41 | arogenate dehydratase 1 [Source:TAIR;Acc:AT1G11790](projected from arabidopsis_thaliana,AT1G11790) | lyase |
| Tal14c | 89 | Bo2g139040 | 5,15E+00 | 4,77E-27 |  | thymidine kinase |
| Tal14c | 90 | Bo3g024650 | 5,14E+00 | 1,07E-11 |  | trans-cinnamate 4-monooxygenase |
| Tal14c | 91 | Bo2g014290 | 5,12E+00 | 1,23E-14 |  | transmembrane protein |
| Tal14c | 92 | Bo6g116570 | 5,06E+00 | 5,52E-14 |  | proteinase inhibitor I3, Kunitz legume, kunitz inhibitor STI-like superfamily |
| Tal14c | 93 | Bo4g149550 | 5,06E+00 | 2,12E-26 | cytochrome P450, family 79, subfamily B, polypeptide 3 [Source:TAIR;Acc:AT2G22330](projected from arabidopsis_thaliana,AT2G22330) | tryptophan N-monooxygenase |
| Tal14c | 94 | Bo3g076030 | 5,05E+00 | 4,71E-38 |  | MATH/TRAF domain-containing protein |
| Tal14c | 95 | Bo3g009050 | 5,05E+00 | 9,76E-19 |  | transcription factor TIFY family |
| Tal14c | 96 | Bo2g002340 | 5,04E+00 | 1,34E-16 | HXXXD-type acyl-transferase family protein [Source:TAIR;Acc:AT5G01210](projected from arabidopsis_thaliana,AT5G01210) | shikimate O-hydroxycinnamoyltransferase |
| Tal14c | 97 | Bo7g104580 | 5,00E+00 | 4,89E-13 | cellulose synthase-like A01 [Source:TAIR;Acc:AT4G16590](projected from arabidopsis_thaliana,AT4G16590) | glucomannan 4-beta-mannosyltransferase |
| Tal14c | 98 | Bo6g116530 | 4,99E+00 | 5,37E-54 | CONTAINS InterPro DOMAIN/s: Nucleoporin protein Ndc1-Nup (InterPro:IPR019049); Has 36 Blast hits to 36 proteins in 17 species: Archae - 0; Bacteria - 0; Metazoa - 1; Fungi - 0; Plants - 35; Viruses - 0; Other Eukaryotes - 0 (source: NCBI BLink). [Source:TAIR;Acc:AT1G73240](projected from arabidopsis_thaliana,AT1G73240) | nucleoporin protein Ndc1-Nup |
| Tal14c | 99 | Bo5g005220 | 4,99E+00 | 6,01E-20 | mitogen-activated protein kinase kinase kinase 18 [Source:TAIR;Acc:AT1G05100](projected from arabidopsis_thaliana,AT1G05100) | mitogen-activated protein kinase kinase kinase STE-STE11 family |
| Tal14c | 100 | Bo7g063950 | 4,93E+00 | 6,75E-19 |  | putative protein |
| Tal14c | 101 | Bo3g041960 | 4,90E+00 | 8,56E-21 | tyrosine aminotransferase 3 [Source:Projected from Arabidopsis thaliana (AT2G24850) TAIR;Acc:AT2G24850] | aminotransferase, class-I, pyridoxal-phosphate-binding, aminotransferase, class I/classII |
| Tal14c | 102 | Bo5g103450 | 4,86E+00 | 8,80E-27 |  | 4-coumarate--CoA ligase |
| Tal14c | 103 | Bo4g136980 | 4,86E+00 | 3,93E-13 | HXXXD-type acyl-transferase family protein [Source:TAIR;Acc:AT5G38130](projected from arabidopsis_thaliana,AT5G38130) | shikimate O-hydroxycinnamoyltransferase |
| Tal14c | 104 | Bo3g004360 | 4,86E+00 | 7,39E-30 | PMSR2 [Source:Projected from Arabidopsis thaliana (AT5G07460) UniProtKB/TrEMBL;Acc:A0A178UBC0] | peptide-methionine (S)-S-oxide reductase |
| Tal14c | 105 | Bo3g064930 | 4,84E+00 | 2,55E-21 |  | chitinase |
| Tal14c | 106 | Bo1g151860 | 4,82E+00 | 7,74E-16 | ARM repeat superfamily protein [Source:Projected from Arabidopsis thaliana (AT3G05040) UniProtKB/TrEMBL;Acc:A0A1I9LRW7] | importin-beta domain, armadillo-like helical, exportin-1/Importin-beta, exportin-1/5 |
| Tal14c | 107 | Bo3g003910 | 4,81E+00 | 9,75E-10 |  | carboxylesterase |
| Tal14c | 108 | Bo4g187790 | 4,79E+00 | 1,79E-15 | Annexin [Source:UniProtKB/TrEMBL;Acc:A0A0D3C513] | annexin |
| Tal14c | 109 | Bo2g075330 | 4,78E+00 | 5,80E-15 |  | transferase |
| Tal14c | 110 | Bo4g086590 | 4,78E+00 | 1,32E-16 |  | geranyllinalool synthase |
| Tal14c | 111 | Bo2g050630 | 4,75E+00 | 6,25E-13 | Lipoxygenase [Source:UniProtKB/TrEMBL;Acc:A0A0D3AN05] | linoleate 13S-lipoxygenase |
| Tal14c | 112 | Bo3g104710 | 4,75E+00 | 1,04E-13 |  | calcineurin-like phosphoesterase domain, ApaH type, metallo-dependent phosphatase |
| Tal14c | 113 | Bo7g063820 | 4,70E+00 | 1,03E-23 |  | cytochrome P450 |
| Tal14c | 114 | Bo9g054490 | 4,70E+00 | 2,42E-20 | Pectin lyase-like superfamily protein [Source:TAIR;Acc:AT1G56710](projected from arabidopsis_thaliana,AT1G56710) | endo-polygalacturonase |
| Tal14c | 115 | Bo5g139630 | 4,70E+00 | 5,46E-41 | Protein SCO1 homolog 1, mitochondrial [Source:Projected from Arabidopsis thaliana (AT3G08950) UniProtKB/Swiss-Prot;Acc:Q8VYP0] | copper chaperone SCO1/SenC, Thioredoxin domain, Thioredoxin-like superfamily |
| Tal14c | 116 | Bo6g097400 | 4,69E+00 | 1,31E-09 | Lipoxygenase [Source:UniProtKB/TrEMBL;Acc:A0A0D3CXY3] | linoleate 13S-lipoxygenase |
| Tal14c | 117 | Bo2g002440 | 4,68E+00 | 1,81E-14 | MD-2-related lipid recognition domain-containing protein [Source:TAIR;Acc:AT2G26370](projected from arabidopsis_thaliana,AT2G26370) | sterol transport protein NPC2 |
| Tal14c | 118 | Bo9g028200 | 4,66E+00 | 1,58E-15 |  | tyrosine decarboxylase |
| Tal14c | 119 | Bo7g116730 | 4,64E+00 | 3,06E-33 | D-3-phosphoglycerate dehydrogenase [Source:TAIR;Acc:AT4G34200](projected from arabidopsis_thaliana,AT4G34200) | phosphoglycerate dehydrogenase |
| Tal14c | 120 | Bo2g116210 | 4,63E+00 | 5,08E-33 | allene oxide synthase [Source:TAIR;Acc:AT5G42650](projected from arabidopsis_thaliana,AT5G42650) | hydroperoxide dehydratase |
| Tal14c | 121 | Bo5g024040 | 4,61E+00 | 9,27E-20 | Leucine-rich repeat receptor-like protein kinase PEPR2 [Source:Projected from Arabidopsis thaliana (AT1G17750) UniProtKB/Swiss-Prot;Acc:Q9FZ59] | protein kinase RLK-Pelle-LRR-XI-1 family |
| Tal14c | 122 | Bo01561s020 | 4,55E+00 | 2,95E-09 |  | geranyllinalool synthase |
| Tal14c | 123 | Bo01172s040 | 4,54E+00 | 3,74E-21 |  | cytochrome P450 |
| Tal14c | 124 | Bo6g101200 | 4,53E+00 | 1,78E-16 |  | juvenile hormone-III synthase |
| Tal14c | 125 | Bo3g101000 | 4,53E+00 | 4,50E-15 | WAT1-related protein [Source:UniProtKB/TrEMBL;Acc:A0A0D3BF10] | EamA domain-containing protein |
| Tal14c | 126 | Bo2g064710 | 4,52E+00 | 3,23E-22 |  | actin-crosslinking |
| Tal14c | 127 | Bo6g094620 | 4,50E+00 | 2,00E-42 |  | transcription factor TIFY family |
| Tal14c | 128 | Bo3g032760 | 4,50E+00 | 5,08E-42 |  | Annexin superfamily |
| Tal14c | 129 | Bo6g051550 | 4,50E+00 | 5,02E-29 |  | serine O-acetyltransferase |
| Tal14c | 130 | Bo6g116560 | 4,49E+00 | 5,56E-10 | Similar to Alpha-amylase/subtilisin inhibitor (RASI) [Source: Projected from Oryza sativa (Os04g0526600)] | proteinase inhibitor I3, Kunitz legume, kunitz inhibitor STI-like superfamily |
| Tal14c | 131 | Bo1g050940 | 4,49E+00 | 1,32E-28 | vacuolar iron transporter (VIT) family protein [Source:TAIR;Acc:AT4G27860](projected from arabidopsis_thaliana,AT4G27860) | putative protein |
| Tal14c | 132 | Bo5g078730 | 4,48E+00 | 2,68E-10 |  | MATH/TRAF domain-containing protein |
| Tal14c | 133 | Bo9g172170 | 4,45E+00 | 1,56E-28 | Protein DETOXIFICATION [Source:UniProtKB/TrEMBL;Acc:A0A0D3EGH1] | multi antimicrobial extrusion protein |
| Tal14c | 134 | Bo4g154660 | 4,45E+00 | 8,91E-14 |  | cytochrome P450 |
| Tal14c | 135 | Bo9g058960 | 4,44E+00 | 4,45E-13 |  | protein kinase RLK-Pelle-DLSV family |
| Tal14c | 136 | Bo4g106230 | 4,44E+00 | 6,93E-04 |  | beta-glucosidase |
| Tal14c | 137 | Bo3g083900 | 4,43E+00 | 3,94E-19 | cytochrome P450, family 82, subfamily G, polypeptide 1 [Source:TAIR;Acc:AT3G25180](projected from arabidopsis_thaliana,AT3G25180) | cytochrome P450 |
| Tal14c | 138 | Bo3g052090 | 4,43E+00 | 2,85E-09 |  | (-)-deoxypodophyllotoxin synthase |
| Tal14c | 139 | Bo1g023950 | 4,42E+00 | 1,07E-12 |  | gibberellin 2beta-dioxygenase |
| Tal14c | 140 | Bo1g022520 | 4,42E+00 | 1,11E-10 |  | tetrahydroberberine oxidase |
| Tal14c | 141 | Bo4g116900 | 4,41E+00 | 6,07E-11 |  | 3-oxoacyl-[acyl-carrier-protein] reductase |
| Tal14c | 142 | Bo9g004350 | 4,40E+00 | 7,13E-12 |  | chalcone synthase |
| Tal14c | 143 | Bo4g078590 | 4,39E+00 | 3,80E-10 |  | triacylglycerol lipase |
| Tal14c | 144 | Bo01100s010 | 4,36E+00 | 2,56E-10 |  | proteinase inhibitor I3, Kunitz legume, kunitz inhibitor STI-like superfamily |
| Tal14c | 145 | Bo9g054540 | 4,33E+00 | 3,08E-17 | unknown protein; Has 665200 Blast hits to 205811 proteins in 4684 species: Archae - 3320; Bacteria - 107592; Metazoa - 249086; Fungi - 76753; Plants - 38542; Viruses - 3008; Other Eukaryotes - 186899 (source: NCBI BLink). [Source:TAIR;Acc:AT1G56660](projected from arabidopsis_thaliana,AT1G56660) | putative protein |
| Tal14c | 146 | Bo6g103700 | 4,32E+00 | 8,96E-16 |  | aldehyde oxidase |
| Tal14c | 147 | Bo7g097340 | 4,31E+00 | 3,82E-11 |  | alpha-copaene synthase |
| Tal14c | 148 | Bo14869s010 | 4,29E+00 | 5,30E-10 |  | cytochrome P450 |
| Tal14c | 149 | Bo3g024720 | 4,27E+00 | 3,18E-14 |  | thioredoxin-disulfide reductase |
| Tal14c | 150 | Bo8g100000 | 4,24E+00 | 2,23E-32 |  | lyase |
| Tal14c | 151 | Bo3g003410 | 4,24E+00 | 1,33E-10 | Uncharacterized protein (Fragment) [Source:UniProtKB/TrEMBL;Acc:A0A0D3AZV6] | anthocyanidin synthase |
| Tal14c | 152 | Bo2g092580 | 4,23E+00 | 1,85E-18 |  | O-methyltransferase domain, plant methyltransferase dimerisation domain-containing protein |
| Tal14c | 153 | Bo5g108630 | 4,23E+00 | 5,80E-45 |  | hypothetical protein |
| Tal14c | 154 | Bo1g121630 | 4,22E+00 | 2,54E-18 |  | stress-response A/B barrel domain-containing protein HS1/DABB1 |
| Tal14c | 155 | Bo3g062370 | 4,22E+00 | 3,30E-08 | PYRIMIDINE 4 [Source:TAIR;Acc:AT3G08860](projected from arabidopsis_thaliana,AT3G08860) | alanine--glyoxylate transaminase |
| Tal14c | 156 | Bo8g105040 | 4,21E+00 | 1,26E-29 |  | glutathione transferase |
| Tal14c | 157 | Bo2g028660 | 4,20E+00 | 2,52E-07 |  | protein Networked (NET), actin-binding (NAB) |
| Tal14c | 158 | Bo4g021130 | 4,18E+00 | 1,49E-25 |  | thiol S-methyltransferase |
| Tal14c | 159 | Bo1g002960 | 4,18E+00 | 6,43E-50 | Adenylyl-sulfate kinase [Source:UniProtKB/TrEMBL;Acc:A0A0D3A119] | adenylyl-sulfate kinase |
| Tal14c | 160 | Bo4g194980 | 4,17E+00 | 7,83E-13 | Beta-glucosidase 15 [Source:Projected from Arabidopsis thaliana (AT2G44450) UniProtKB/Swiss-Prot;Acc:O64879] | beta-glucosidase |
| Tal14c | 161 | Bo5g014860 | 4,17E+00 | 3,21E-21 | Tryptophan synthase [Source:UniProtKB/TrEMBL;Acc:A0A0D3C9C1] | tryptophan synthase |
| Tal14c | 162 | Bo4g024110 | 4,17E+00 | 1,37E-05 | alpha/beta-Hydrolases superfamily protein [Source:TAIR;Acc:AT2G44810](projected from arabidopsis_thaliana,AT2G44810) | phospholipase A1 |
| Tal14c | 163 | Bo9g148400 | 4,14E+00 | 2,47E-37 | Fatty acyl-CoA reductase [Source:UniProtKB/TrEMBL;Acc:A0A0D3EDD1] | oxidoreductase |
| Tal14c | 164 | Bo7g114110 | 4,13E+00 | 4,38E-08 |  | transcription factor bHLH family |
| Tal14c | 165 | Bo1g065080 | 4,13E+00 | 9,83E-06 | High affinity nitrate transporter 2.6 [Source:Projected from Arabidopsis thaliana (AT3G45060) UniProtKB/Swiss-Prot;Acc:Q9LXH0] | major facilitator superfamily, MFS transporter superfamily |
| Tal14c | 166 | Bo3g042930 | 4,11E+00 | 4,63E-08 |  | small auxin-up RNA |
| Tal14c | 167 | Bo5g062250 | 4,11E+00 | 7,31E-10 |  | gibberellin 2beta-dioxygenase |
| Tal14c | 168 | Bo00722s060 | 4,09E+00 | 3,55E-11 |  | proteinase inhibitor I3, Kunitz legume, kunitz inhibitor STI-like superfamily |
| Tal14c | 169 | Bo3g162840 | 4,09E+00 | 2,40E-15 |  | transcription factor bHLH family |
| Tal14c | 170 | Bo9g177260 | 4,08E+00 | 1,03E-09 |  | phenylalanine N-monooxygenase |
| Tal14c | 171 | Bo4g086600 | 4,06E+00 | 5,15E-24 |  | geranyllinalool synthase |
| Tal14c | 172 | Bo7g065090 | 4,06E+00 | 9,72E-11 | Uncharacterized protein (Fragment) [Source:UniProtKB/TrEMBL;Acc:A0A0D3D8J2] | thiosulfate sulfurtransferase |
| Tal14c | 173 | Bo4g136080 | 4,05E+00 | 4,48E-07 |  | ABC-type xenobiotic transporter |
| Tal14c | 174 | Bo9g176940 | 4,05E+00 | 7,42E-47 |  | anthranilate synthase |
| Tal14c | 175 | Bo01163s010 | 4,04E+00 | 2,66E-08 | Lipoxygenase [Source:UniProtKB/TrEMBL;Acc:A0A0D2ZTZ5] | linoleate 13S-lipoxygenase |
| Tal14c | 176 | Bo3g004390 | 4,02E+00 | 1,61E-12 |  | peptide-methionine (S)-S-oxide reductase |
| Tal14c | 177 | Bo3g149810 | 4,02E+00 | 2,54E-32 |  | agmatine N(4)-coumaroyltransferase |
| Tal14c | 178 | Bo3g044760 | 4,01E+00 | 1,36E-11 |  | flavanone 3-dioxygenase |
| Tal14c | 179 | Bo1g016130 | 4,01E+00 | 2,79E-26 |  | nucleotide diphosphatase |
| Tal14c | 180 | Bo3g147640 | 4,01E+00 | 6,73E-12 |  | transcription factor TIFY family |
| Tal14c | 181 | Bo9g180100 | 3,99E+00 | 2,09E-25 |  | methyltransferase |
| Tal14c | 182 | Bo2g009330 | 3,97E+00 | 4,73E-07 |  | triacylglycerol lipase |
| Tal14c | 183 | Bo3g003970 | 3,96E+00 | 2,67E-14 |  | transcription factor Homobox-WOX family |
| Tal14c | 184 | Bo6g051490 | 3,95E+00 | 5,79E-50 |  | cellulose synthase (UDP-forming) |
| Tal14c | 185 | Bo5g025610 | 3,95E+00 | 1,62E-59 | sulfotransferase 17 [Source:TAIR;Acc:AT1G18590](projected from arabidopsis_thaliana,AT1G18590) | sulfotransferase |
| Tal14c | 186 | Bo9g150050 | 3,94E+00 | 8,75E-41 |  | oxidoreductase |
| Tal14c | 187 | Bo4g020890 | 3,94E+00 | 1,11E-09 |  | scorpion long chain toxin/defensin, knottin, scorpion toxin |
| Tal14c | 188 | Bo01243s020 | 3,94E+00 | 1,64E-12 |  | glutaredoxin, Thioredoxin-like superfamily |
| Tal14c | 189 | Bo7g095340 | 3,93E+00 | 1,32E-06 |  | ubiquitinyl hydrolase 1 |
| Tal14c | 190 | Bo5g113720 | 3,92E+00 | 1,32E-06 |  | branched-chain-amino-acid transaminase |
| Tal14c | 191 | Bo7g031500 | 3,92E+00 | 1,58E-44 |  | beta-glucosidase |
| Tal14c | 192 | Bo1g139550 | 3,91E+00 | 7,98E-14 | Leucine-rich repeat (LRR) family protein [Source:Projected from Arabidopsis thaliana (AT3G12145) TAIR;Acc:AT3G12145] | leucine-rich repeat-containing, plant-type, leucine-rich repeat domain superfamily |
| Tal14c | 193 | Bo4g094360 | 3,91E+00 | 2,58E-23 | Sulfotransferase [Source:UniProtKB/TrEMBL;Acc:A0A0D3BVV8] | flavonol 3-sulfotransferase |
| Tal14c | 194 | Bo1g011680 | 3,90E+00 | 8,09E-08 |  | palmitoyl-protein hydrolase |
| Tal14c | 195 | Bo3g064990 | 3,90E+00 | 1,76E-17 | sulfate transporter 4;2 [Source:TAIR;Acc:AT3G12520](projected from arabidopsis_thaliana,AT3G12520) | SLC26A/SulP transporter, STAS domain, sulphate anion transporter, STAS domain superfamily |
| Tal14c | 196 | Bo06739s010 | 3,90E+00 | 1,71E-16 |  | geranyllinalool synthase |
| Tal14c | 197 | Bo4g046030 | 3,88E+00 | 4,62E-10 |  | mitogen-activated protein kinase kinase kinase STE-STE11 family |
| Tal14c | 198 | Bo8g099540 | 3,87E+00 | 4,10E-08 | Probable aminotransferase TAT3 [Source:Projected from Arabidopsis thaliana (AT2G24850) UniProtKB/Swiss-Prot;Acc:Q9SK47] | aminotransferase, class-I, pyridoxal-phosphate-binding, aminotransferase, class I/classII |
| Tal14c | 199 | Bo7g087380 | 3,87E+00 | 5,48E-10 |  | putative protein |
| Tal14c | 200 | Bo8g076760 | 3,86E+00 | 7,42E-11 |  | flavonol synthase |
| Tal14c | 201 | Bo7g077450 | 3,86E+00 | 3,51E-20 | nudix hydrolase homolog 8 [Source:TAIR;Acc:AT5G47240](projected from arabidopsis_thaliana,AT5G47240) | hydrolase |
| Tal14c | 202 | Bo1g023960 | 3,86E+00 | 5,74E-08 |  | gibberellin 2beta-dioxygenase |
| Tal14c | 203 | Bo5g126180 | 3,86E+00 | 8,86E-18 |  | transcription factor MYB-HB-like family |
| Tal14c | 204 | Bo4g188070 | 3,85E+00 | 5,93E-34 |  | jacalin-like lectin domain-containing protein |
| Tal14c | 205 | Bo5g098740 | 3,85E+00 | 2,01E-09 | Glycosyltransferase [Source:UniProtKB/TrEMBL;Acc:A0A0D3CHG3] | flavonol 3-O-glucosyltransferase |
| Tal14c | 206 | Bo6g116740 | 3,85E+00 | 1,78E-10 | Cytochrome P450 superfamily protein [Source:TAIR;Acc:AT1G73340](projected from arabidopsis_thaliana,AT1G73340) | abieta-7,13-dien-18-ol hydroxylase |
| Tal14c | 207 | Bo2g012130 | 3,84E+00 | 6,15E-07 |  | NAD-dependent epimerase/dehydratase, NAD(P)-binding domain superfamily |
| Tal14c | 208 | Bo8g040630 | 3,84E+00 | 2,75E-12 | Beta-amylase [Source:UniProtKB/TrEMBL;Acc:A0A0D3DLY3] | beta-amylase |
| Tal14c | 209 | Bo3g052200 | 3,84E+00 | 1,58E-15 |  | like-Sm (LSM) domain containing protein, LSm4/SmD1/SmD3 |
| Tal14c | 210 | Bo2g092470 | 3,84E+00 | 3,80E-10 |  | RNA-directed DNA polymerase |
| Tal14c | 211 | Bo8g050630 | 3,83E+00 | 8,41E-09 |  | thebaine 6-O-demethylase |
| Tal14c | 212 | Bo6g072600 | 3,83E+00 | 7,58E-24 |  | strictosidine synthase transcription factor WD40-like family |
| Tal14c | 213 | Bo8g106320 | 3,82E+00 | 1,72E-25 | 10 kDa chaperonin, mitochondrial [Source:Projected from Arabidopsis thaliana (AT1G14980) UniProtKB/Swiss-Prot;Acc:P34893] | GroES-like superfamily, chaperonin GroES, groES chaperonin family |
| Tal14c | 214 | Bo5g030280 | 3,81E+00 | 9,84E-09 | SKU5 similar 7 [Source:Projected from Arabidopsis thaliana (AT1G21860) UniProtKB/TrEMBL;Acc:Q9SFF1] | L-ascorbate oxidase |
| Tal14c | 215 | Bo5g103380 | 3,81E+00 | 4,45E-25 |  | 4-coumarate--CoA ligase |
| Tal14c | 216 | Bo8g025760 | 3,80E+00 | 1,50E-24 | Annexin [Source:UniProtKB/TrEMBL;Acc:A0A0D3DKI6] | peroxidase |
| Tal14c | 217 | Bo00722s100 | 3,80E+00 | 5,13E-11 |  | proteinase inhibitor I3, Kunitz legume, kunitz inhibitor STI-like superfamily |
| Tal14c | 218 | Bo00921s010 | 3,80E+00 | 1,18E-22 |  | geranyllinalool synthase |
| Tal14c | 219 | Bo1g101970 | 3,79E+00 | 3,85E-16 |  | cytochrome P450 |
| Tal14c | 220 | Bo4g020910 | 3,78E+00 | 8,99E-11 |  | scorpion long chain toxin/defensin, knottin, scorpion toxin |
| Tal14c | 221 | Bo5g013240 | 3,78E+00 | 3,64E-36 | phosphoribosyl pyrophosphate (PRPP) synthase 3 [Source:TAIR;Acc:AT1G10700](projected from arabidopsis_thaliana,AT1G10700) | ribose-phosphate diphosphokinase |
| Tal14c | 222 | Bo3g181210 | 3,78E+00 | 7,80E-10 |  | jacalin-like lectin domain, kelch-type beta propeller |
| Tal14c | 223 | Bo2g165690 | 3,77E+00 | 7,34E-15 |  | jasmonoyl-L-amino acid 12-hydroxylase |
| Tal14c | 224 | Bo8g104870 | 3,77E+00 | 2,02E-20 | Lipoxygenase [Source:UniProtKB/TrEMBL;Acc:A0A0D3DWT7] | linoleate 13S-lipoxygenase |
| Tal14c | 225 | Bo7g064080 | 3,76E+00 | 6,29E-07 | mitogen-activated protein kinase kinase kinase 19 [Source:Projected from Arabidopsis thaliana (AT5G67080) TAIR;Acc:AT5G67080] | mitogen-activated protein kinase kinase kinase STE-STE11 family |
| Tal14c | 226 | Bo8g118250 | 3,75E+00 | 4,51E-06 |  | transcription factor AP2-EREBP family |
| Tal14c | 227 | Bo5g143400 | 3,75E+00 | 3,35E-15 |  | N-acetylneuraminate 7-O(or 9-O)-acetyltransferase |
| Tal14c | 228 | Bo6g083780 | 3,73E+00 | 3,46E-17 |  | EF-hand domain-containing protein |
| Tal14c | 229 | Bo1g144550 | 3,72E+00 | 1,08E-11 | Sulfotransferase [Source:UniProtKB/TrEMBL;Acc:A0A0D3AEX2] | flavonol 3-sulfotransferase |
| Tal14c | 230 | Bo9g055870 | 3,71E+00 | 2,26E-03 | Late embryogenesis abundant (LEA) protein-related [Source:TAIR;Acc:AT1G54890](projected from arabidopsis_thaliana,AT1G54890) | putative protein |
| Tal14c | 231 | Bo7g040400 | 3,71E+00 | 6,51E-06 |  | geraniol 8-hydroxylase |
| Tal14c | 232 | Bo1g138910 | 3,71E+00 | 1,54E-09 | Auxin-responsive protein SAUR72 [Source:Projected from Arabidopsis thaliana (AT3G12830) UniProtKB/Swiss-Prot;Acc:Q9LTV3] | small auxin-up RNA |
| Tal14c | 233 | Bo7g118850 | 3,69E+00 | 1,23E-38 | Adenylyl-sulfate kinase [Source:UniProtKB/TrEMBL;Acc:A0A0D3DI87] | adenylyl-sulfate kinase |
| Tal14c | 234 | Bo2g130260 | 3,68E+00 | 6,65E-11 |  | aluminum-activated malate transporter |
| Tal14c | 235 | Bo7g089560 | 3,68E+00 | 8,89E-13 |  | tetratricopeptide repeat protein POLLENLESS 3/SULFUR DEFICIENCY-INDUCED 1 |
| Tal14c | 236 | Bo6g039700 | 3,68E+00 | 4,33E-24 |  | beta-glucosidase |
| Tal14c | 237 | Bo3g017440 | 3,67E+00 | 4,20E-07 | Concanavalin A-like lectin protein kinase family protein [Source:TAIR;Acc:AT5G60320](projected from arabidopsis_thaliana,AT5G60320) | protein kinase RLK-Pelle-L-LEC family |
| Tal14c | 238 | Bo7g045650 | 3,67E+00 | 1,74E-07 |  | glutathione transferase |
| Tal14c | 239 | Bo1g139500 | 3,64E+00 | 1,88E-23 | Serine carboxypeptidase-like 17 [Source:Projected from Arabidopsis thaliana (AT3G12203) UniProtKB/Swiss-Prot;Acc:Q9C7D6] | sinapoylglucose--sinapoylglucose O-sinapoyltransferase |
| Tal14c | 240 | Bo6g026320 | 3,64E+00 | 6,07E-35 |  | jasmonoyl-L-amino acid hydrolase |
| Tal14c | 241 | Bo9g157290 | 3,64E+00 | 1,41E-11 | Aquaporin-like superfamily protein [Source:TAIR;Acc:AT5G18290](projected from arabidopsis_thaliana,AT5G18290) | major intrinsic protein |
| Tal14c | 242 | Bo4g198560 | 3,64E+00 | 1,78E-11 |  | transcription factor WRKY family |
| Tal14c | 243 | Bo7g063790 | 3,64E+00 | 8,01E-08 |  | cytochrome P450 |
| Tal14c | 244 | Bo6g039690 | 3,63E+00 | 7,39E-30 |  | putative protein |
| Tal14c | 245 | Bo2g075620 | 3,63E+00 | 6,93E-04 |  | proteinase inhibitor I3, Kunitz legume, kunitz inhibitor STI-like superfamily |
| Tal14c | 246 | Bo4g139360 | 3,62E+00 | 5,89E-05 | P-loop containing nucleoside triphosphate hydrolases superfamily protein [Source:TAIR;Acc:AT5G40000](projected from arabidopsis_thaliana,AT5G40000) | AAA+ ATPase domain, ATPase, AAA-type, core, AAA-type ATPase domain-containing protein |
| Tal14c | 247 | Bo9g028760 | 3,62E+00 | 1,06E-29 | basic helix-loop-helix (bHLH) DNA-binding superfamily protein [Source:TAIR;Acc:AT1G62975](projected from arabidopsis_thaliana,AT1G62975) | transcription factor bHLH family |
| Tal14c | 248 | Bo1g055140 | 3,62E+00 | 3,05E-07 |  | putative protein |
| Tal14c | 249 | Bo8g040610 | 3,62E+00 | 4,17E-10 | Beta-amylase [Source:UniProtKB/TrEMBL;Acc:A0A0D3DLY1] | beta-amylase |
| Tal14c | 250 | Bo3g049860 | 3,62E+00 | 4,09E-33 |  | cytochrome P450, NPH3 domain, cytochrome P450 superfamily, NPH3/RPT2-like family |
| Tal14c | 251 | Bo4g186450 | 3,62E+00 | 2,46E-32 | S-adenosylmethionine synthase [Source:UniProtKB/TrEMBL;Acc:A0A0D3C4M9] | methionine adenosyltransferase |
| Tal14c | 252 | Bo6g032990 | 3,61E+00 | 5,99E-09 |  | galactose oxidase/kelch, beta-propeller, kelch-type beta propeller |
| Tal14c | 253 | Bo5g027180 | 3,61E+00 | 6,79E-27 |  | carboxylesterase |
| Tal14c | 254 | Bo8g102890 | 3,60E+00 | 2,58E-16 |  | transcription factor TIFY family |
| Tal14c | 255 | Bo3g019580 | 3,60E+00 | 5,22E-08 |  | putative protein |
| Tal14c | 256 | Bo8g100510 | 3,59E+00 | 6,94E-10 |  | P-type Ca(2+) transporter |
| Tal14c | 257 | Bo01291s010 | 3,59E+00 | 2,83E-06 | Glycosyltransferase [Source:UniProtKB/TrEMBL;Acc:A0A0D2ZUJ7] | flavonol 3-O-glucosyltransferase |
| Tal14c | 258 | Bo00722s130 | 3,59E+00 | 6,23E-05 |  | proteinase inhibitor I3, Kunitz legume, kunitz inhibitor STI-like superfamily |
| Tal14c | 259 | Bo1g007550 | 3,57E+00 | 5,38E-25 | D-3-phosphoglycerate dehydrogenase [Source:UniProtKB/TrEMBL;Acc:A0A0D3A229] | phosphoglycerate dehydrogenase |
| Tal14c | 260 | Bo6g119630 | 3,56E+00 | 5,24E-13 | cytokinin oxidase 5 [Source:TAIR;Acc:AT1G75450](projected from arabidopsis_thaliana,AT1G75450) | cytokinin dehydrogenase |
| Tal14c | 261 | Bo1g156260 | 3,56E+00 | 1,56E-08 |  | delta(7)-sterol 5(6)-desaturase |
| Tal14c | 262 | Bo2g093210 | 3,56E+00 | 1,06E-10 |  | RNA-directed DNA polymerase |
| Tal14c | 263 | Bo6g003330 | 3,56E+00 | 7,27E-18 | IAA-leucine resistant (ILR)-like gene 6 [Source:TAIR;Acc:AT1G44350](projected from arabidopsis_thaliana,AT1G44350) | jasmonoyl-L-amino acid hydrolase |
| Tal14c | 264 | Bo6g099590 | 3,56E+00 | 5,82E-43 | Isocitrate dehydrogenase [NADP] [Source:UniProtKB/TrEMBL;Acc:A0A0D3CYA3] | isocitrate dehydrogenase (NADP(+)) |
| Tal14c | 265 | Bo1g036850 | 3,55E+00 | 3,82E-18 | Tyrosine transaminase family protein [Source:TAIR;Acc:AT4G23600](projected from arabidopsis_thaliana,AT4G23600) | cysteine-S-conjugate beta-lyase |
| Tal14c | 266 | Bo3g168910 | 3,54E+00 | 7,17E-11 |  | transcription factor AP2-EREBP family |
| Tal14c | 267 | Bo2g023330 | 3,54E+00 | 6,44E-06 |  | oxidoreductase |
| Tal14c | 268 | Bo3g078640 | 3,53E+00 | 1,05E-17 | 1-deoxy-D-xylulose 5-phosphate synthase 1 [Source:TAIR;Acc:AT3G21500](projected from arabidopsis_thaliana,AT3G21500) | 1-deoxy-D-xylulose-5-phosphate synthase |
| Tal14c | 269 | Bo2g161650 | 3,52E+00 | 3,77E-11 |  | stress up-regulated Nod 19 |
| Tal14c | 270 | Bo4g013290 | 3,51E+00 | 1,34E-09 |  | PLAT/LH2 domain superfamily protein |
| Tal14c | 271 | Bo03100s010 | 3,49E+00 | 1,62E-16 | Beta-amylase [Source:UniProtKB/TrEMBL;Acc:A0A0D2ZWI6] | beta-amylase |
| Tal14c | 272 | Bo1g116240 | 3,49E+00 | 1,68E-08 |  | RING-type E3 ubiquitin transferase transcription factor C2H2 family |
| Tal14c | 273 | Bo9g094320 | 3,49E+00 | 3,75E-10 |  | O-methyltransferase domain, plant methyltransferase dimerisation domain-containing protein |
| Tal14c | 274 | Bo3g032770 | 3,48E+00 | 7,62E-21 | Annexin [Source:UniProtKB/TrEMBL;Acc:A0A0D3B5Q1] | annexin |
| Tal14c | 275 | Bo6g030940 | 3,46E+00 | 6,58E-21 | NAC domain containing protein 19 [Source:TAIR;Acc:AT1G52890](projected from arabidopsis_thaliana,AT1G52890) | transcription factor NAM family |
| Tal14c | 276 | Bo7g099340 | 3,46E+00 | 3,24E-24 | methyl esterase 10 [Source:TAIR;Acc:AT3G50440](projected from arabidopsis_thaliana,AT3G50440) | alpha/beta hydrolase-1, methylesterase/Alpha-hydroxynitrile lyase |
| Tal14c | 277 | Bo8g050000 | 3,46E+00 | 2,82E-06 |  | purine permease, plant |
| Tal14c | 278 | Bo6g080740 | 3,45E+00 | 6,85E-12 |  | transcription factor WRKY family |
| Tal14c | 279 | Bo3g093960 | 3,45E+00 | 3,95E-08 |  | cytochrome P450 |
| Tal14c | 280 | Bo4g139430 | 3,44E+00 | 2,15E-05 | Thioredoxin superfamily protein [Source:TAIR;Acc:AT4G12170](projected from arabidopsis_thaliana,AT4G12170) | monodehydroascorbate reductase (NADH) |
| Tal14c | 281 | Bo4g181640 | 3,44E+00 | 5,82E-06 | vesicle-associated membrane protein 723 [Source:TAIR;Acc:AT2G33110](projected from arabidopsis_thaliana,AT2G33110) | Longin domain, v-SNARE, coiled-coil domain-containing protein |
| Tal14c | 282 | Bo3g051010 | 3,43E+00 | 2,91E-17 |  | cytochrome P450 |
| Tal14c | 283 | Bo3g143920 | 3,42E+00 | 2,11E-06 |  | precursor of CEP13/CEP14 |
| Tal14c | 284 | Bo4g186980 | 3,42E+00 | 4,14E-03 |  | trans-2,3-dihydro-3-hydroxyanthranilate isomerase |
| Tal14c | 285 | Bo5g103390 | 3,42E+00 | 2,38E-08 |  | 4-coumarate--CoA ligase |
| Tal14c | 286 | Bo6g079770 | 3,42E+00 | 1,01E-07 |  | carboxylesterase |
| Tal14c | 287 | Bo8g104890 | 3,41E+00 | 5,96E-13 |  | transcription factor TIFY family |
| Tal14c | 288 | Bo2g092530 | 3,41E+00 | 3,91E-16 |  | 2-acylglycerol O-acyltransferase |
| Tal14c | 289 | Bo3g003600 | 3,41E+00 | 1,17E-05 |  | bursehernin 5'-monooxygenase |
| Tal14c | 290 | Bo2g021960 | 3,41E+00 | 1,81E-08 |  | germin, rmlC-like cupin domain superfamily, rmlC-like jelly roll |
| Tal14c | 291 | Bo9g165610 | 3,40E+00 | 1,52E-04 | Glycosyltransferase [Source:UniProtKB/TrEMBL;Acc:A0A0D3EFG2] | flavanone 7-O-beta-glucosyltransferase |
| Tal14c | 292 | Bo9g169450 | 3,40E+00 | 5,92E-09 |  | PADRE domain-containing protein |
| Tal14c | 293 | Bo4g195230 | 3,40E+00 | 1,20E-12 | unknown protein; CONTAINS InterPro DOMAIN/s: Uncharacterised protein family UPF0565 (InterPro:IPR018881); BEST Arabidopsis thaliana protein match is: unknown protein (TAIR:AT2G45380.1); Has 138 Blast hits to 138 proteins in 53 species: Archae - 0; B /.../a - 0; Metazoa - 73; Fungi - 0; Plants - 62; Viruses - 0; Other Eukaryotes - 3 (source: NCBI BLink). [Source:TAIR;Acc:AT2G44850](projected from arabidopsis_thaliana,AT2G44850) | mitochondrial protein C2orf69 |
| Tal14c | 294 | Bo9g062900 | 3,40E+00 | 8,02E-11 |  | proton-dependent oligopeptide transporter family, MFS transporter superfamily |
| Tal14c | 295 | Bo9g029180 | 3,39E+00 | 4,35E-05 |  | Gnk2-like domain-containing protein |
| Tal14c | 296 | Bo5g145170 | 3,38E+00 | 9,34E-46 | Plant invertase/pectin methylesterase inhibitor superfamily [Source:TAIR;Acc:AT3G05620](projected from arabidopsis_thaliana,AT3G05620) | pectinesterase |
| Tal14c | 297 | Bo1g156330 | 3,38E+00 | 1,31E-06 | Regulator of chromosome condensation (RCC1) family protein [Source:TAIR;Acc:AT3G02510](projected from arabidopsis_thaliana,AT3G02510) | regulator of chromosome condensation 1/beta-lactamase-inhibitor protein II |
| Tal14c | 298 | Bo2g023390 | 3,37E+00 | 6,22E-32 |  | oxidoreductase |
| Tal14c | 299 | Bo7g117250 | 3,37E+00 | 2,96E-06 |  | acetylserotonin O-methyltransferase |
| Tal14c | 300 | Bo8g028740 | 3,37E+00 | 1,29E-34 | 2Fe-2S ferredoxin-like superfamily protein [Source:TAIR;Acc:AT1G32550](projected from arabidopsis_thaliana,AT1G32550) | 2Fe-2S ferredoxin-type iron-sulfur binding domain, ferredoxin [2Fe-2S], plant |
